# Diverse bacterial pattern recognition receptors sense the core phage proteome

**DOI:** 10.64898/2026.01.04.697583

**Authors:** Hyunbin Lee, Sofia Luengo-Woods, Jianxiu Zhang, Kira S. Makarova, Yuri I. Wolf, Collin Chiu, Simone A. Evans, Junyi Chen, Haopeng Xiao, Liang Feng, Eugene V. Koonin, Alex Gao

**Affiliations:** Department of Biochemistry, Stanford University, Stanford, CA 94305, USA; Department of Microbiology and Immunology, Stanford University, Stanford, CA 94305, USA; Department of Molecular and Cellular Physiology, Stanford University, Stanford, CA 94305, USA; Computational Biology Branch, Division of Intramural Research, National Library of Medicine, National Institutes of Health, Bethesda, MD 20894, USA; Department of Genetics, Stanford University, Stanford, CA 94305, USA; Stanford Cancer Institute, Stanford University, Stanford, CA 94305, USA

## Abstract

Recognition of foreign molecules inside cells is critical for immunity across all domains of life. Proteins of the STAND NTPase superfamily^1,2^, including eukaryotic NOD-like receptors (NLRs), play a central role in this process^3,4^. In bacteria and archaea, although several STAND families sense phage proteins^5–9^, their functional diversity remains largely unexplored. We conducted a systematic phylogenetic analysis of prokaryotic STAND NTPases and identified at least 90 structurally distinct families associated with antiviral defense. We first show that the uncharacterized Avs7 family recognizes the major capsid protein (MCP) of tailed phages. Three cryo-EM structures of *Salmonella enterica* Avs7 reveal an asymmetric, butterfly-shaped tetramer that assembles stepwise via large, MCP-induced conformational changes, incorporating bacterial elongation factor Tu (EF-Tu) as a structural component that enhances defense. Using genetic screens with a library of 687 phage genes, we further show that 13 additional STAND families sense 13 conserved phage proteins, encompassing most of the core structural and replicative components of tailed phages. These include two distinct MCP-sensing families—Avs8 (PD-λ-4) and Avs10 (Erebus/Hypnos/bNACHT64)—and 11 others (Avs11–21) recognizing the portal, portal adaptor, tail nozzle, head–tail connector, tail terminator, tail tube protein, tail assembly chaperone, tape measure protein, DNA polymerase, helicase/RecA-type ATPase, and single-stranded DNA annealing protein, respectively. Together, our findings highlight structurebased pattern recognition and host factor repurposing as fundamental strategies of bacterial immunity.

## Introduction

Intracellular surveillance for foreign molecules is a cornerstone of host defense against pathogens. In animals and plants, a major part of this process is orchestrated by NLRs, which belong to the STAND (Signal Transduction ATPases with Numerous Domains) NTPase superfamily. STANDs typically have a tripartite architecture consisting of a central NTPase domain, a C-terminal sensor, and an N-terminal effector^1,2^. Immune-related members of this superfamily, including NLRs, sense pathogen-associated molecular patterns (PAMPs) and oligomerize to initiate inflammation or programmed cell death^3^. Mammalian NLRs include NAIP, which detects bacterial flagellin and type III secretion systems^10,11^, and NOD1/2, which recognize peptidoglycan fragments of bacterial cell walls^12,13^.

STAND NTPases are also abundant in bacteria and archaea^1,2^, and some of these, dubbed antiviral STANDs (Avs), provide protection against phage infection^4^. We previously demonstrated that four families, Avs1–4, recognize the folds of two conserved phage proteins, the large terminase subunit and portal, through direct binding^5^. Recent studies have revealed other STAND activation mechanisms, including recognition of multiple proteins by a single sensor^6,7^, a jumbophage protein^9^, and host chaperone perturbation^8^. Several additional STANDs have been reported to confer anti-phage protection^4,14–17^, and one branch—the NACHT clade—has been systematically analyzed in the phylogenetic context^16^, but on the whole, the vast repertoire of prokaryotic STANDs remains poorly characterized. Here, we combine large-scale genome mining with structural, biochemical, and genetic analyses to define the landscape of prokaryotic defense-associated STAND proteins and reveal mechanistic principles of their activation.

### Systematic analysis of defense-associated STAND proteins in prokaryotes

To investigate the diversity of the highly divergent defense-associated STANDs, we employed a two-step, systematic approach (Fig. 1a). We first identified a set of 656 ‘seed’ STAND proteins, including known immune-related STANDs^2,4,16–18^ and additional candidates detected by their localization in genomic proximity to other defense-related genes (Supplementary Table 1)^4,19,20^. We then performed relaxed PSI-BLAST searches using the conserved NTPase domains of these seeds as queries, and after several rounds of curation (see Methods), expanded the set to 359,879 diverse representatives (clustered at 75% sequence identity) distributed across 589 STAND clades (Fig. 1b and Supplementary Table 2; see Methods). These sequences included known animal and plant NLRs, although many of these were not included among the seeds, supporting the sensitivity of this approach.

**Figure 1.**
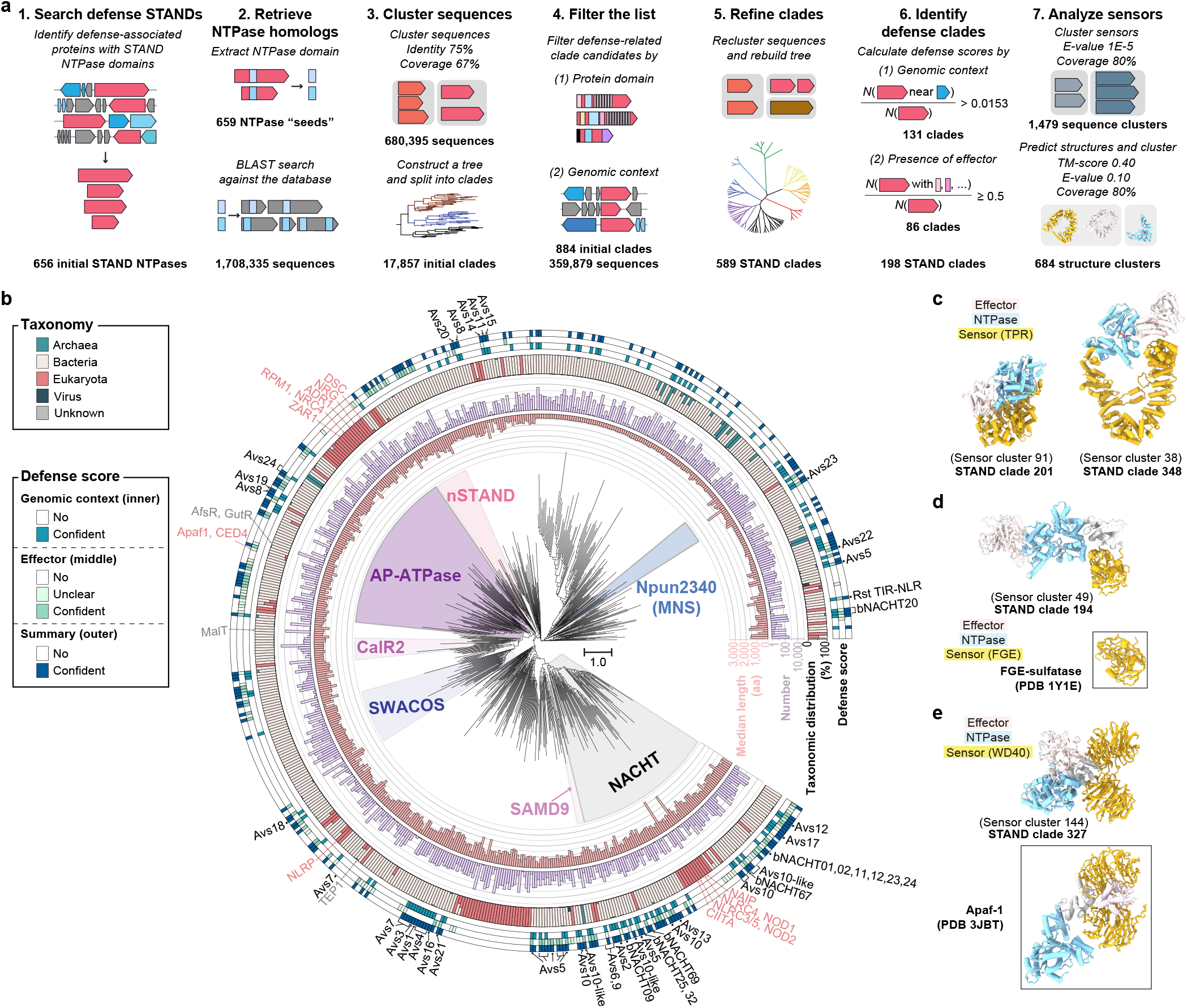
Phylogeny of defense-associated STAND NTPases in prokaryotes. **a**, Schematic of the workflow to identify, analyze, and classify STAND NTPases. **b**, Maximum likelihood tree of the NTPase domains of 589 clades. MNS, NACHT, SWACOS, CalR2, AP–ATPase, and nSTAND refer to previously defined clades. Eukaryotic STANDs are annotated in red. Defense scores were assigned based on two criteria: the frequency of known defense genes in the genomic neighborhood of STAND genes (genomic context) and proportion of STANDs carrying immune-related effector domains (effector; Supplementary Table 3). Clades were classified as confident defense-associated if they satisfied either criterion (see Methods). **c–e**, Representative AlphaFold3 structural models of representatives with helical bundles (**c**), FGE-sulfatase (**d**), or Apaf1-like WD40 β-propellers (**e**) as sensors. Numbers in parentheses represent structural clusters (1–684). Avs, antiviral STAND; TPR, tetratricopeptide repeat; FGE, formylglycine-generating enzyme; Apaf-1, apoptotic protease activating factor 1; bNACHT, bacterial NACHT; SAMD9, sterile alpha motif domain containing 9.

We classified 198 of the 589 STAND clades as defenseassociated based on their proximity to known defense systems and/or the presence of a defense-related effector domain (see Methods; Fig. 1a, Extended Data Fig. 1a, Supplementary Tables 3–4). Our analysis identified CRISPR-associated STANDs^21^ and prophage-encoded STANDs^22^, consistent with their established roles in anti-phage defense (Supplementary Fig. 1a–b). Defense-associated clades are interspersed with non-defense STAND clades, suggestive of multiple functional switches throughout the evolution of the STAND superfamily (Fig. 1b). Known NLRs from animals^23–26^, plants^27,28^, and bacteria^4,5,15–17^ were found in distinct clades, alongside nonimmune STANDs such as transcriptional regulators^29–31^, the telomerase-associated protein TEP1, and the primary cilia protein NPHP3, consistent with earlier analyses^2,32^.

To further explore the diversity of C-terminal sensor domains of the STANDs, which are usually substantially larger than typical defense proteins (Supplementary Fig. 1c), we grouped defense-associated sensors of ≥400 amino acids into 684 clusters by sequence and structural similarity (Extended Data Fig. 1b–f and Supplementary Table 5). Most defense-associated clades were dominated by a single sensor type (Extended Data Fig. 1f). Analysis of the 90 most prevalent clusters revealed contrasting evolutionary patterns among the sensors: some were clade-specific (*e.g*., Avs2), whereas others spanned multiple clades (*e.g*., sensor structural cluster 39; Fig. 1c–e, Extended Data Fig. 1f, Supplementary Fig. 2–4). The majority of the sensors were composed of tetratricopeptide repeat (TPR)-like α-helical repeats (Fig. 1c, Supplementary Fig. 2, 3a, 4), although we also identified other types, including WD40 β-propeller repeats and formylglycine-generating enzyme (FGE) sulfatase-like domains (Fig. 1d–e, Supplementary Fig. 3b–c). This diversity mirrors the sensor repertoire of eukaryotic NLRs^3^.

We identified specific structural similarity between several bacterial sensors and animal proteins involved in immunity. The Avs5 sensor clustered with the SAMD9 family of animal antiviral restriction factors, consistent with recent observations^33,34^ (Supplementary Fig. 2a). We also detected bacterial orthologs of Apaf–1, the core component of the eukaryotic apoptosome^26^, all sharing structurally similar β-propeller sensors (Fig. 1e and Supplementary Fig. 3b). Several bacterial variants encode an N-terminal caspase-like protease in place of the eukaryotic CARD domain that recruits procaspase-9, suggesting a more direct effector mechanism than that in the animal counterparts. The closest Apaf-1 homologs were found in cyanobacteria, consistent with parallels between the defense systems in multicellular bacteria and eukaryotic immunity^35^.

### A shared sensor domain in Avs7 and Dsr1 recognizes phage major capsid proteins

We first selected an uncharacterized STAND family, Avs7, given the consistent presence of its sensor domain across multiple STAND clades, its association with at least 20 distinct N-terminal effector domains, and evidence of extensive horizontal gene transfer (Extended Data Fig. 1f, Supplementary Fig. 5, and Supplementary Table 6). Unexpectedly, the Avs7 sensor is shared by unrelated defense systems lacking the STAND NTPase domain, including defense-associated sirtuin 1 (Dsr1)^4^ (Fig. 2a and Supplementary Table 8). AlphaFold3 models of SeAvs7, *Escherichia coli* Dsr1 (EcDsr1), and a *Streptomyces* HEPN-containing protein revealed a conserved bilobed architecture formed by a sharply bent sensor, suggesting a common recognition mechanism (Extended Data Fig. 2a).

**Figure 2.**
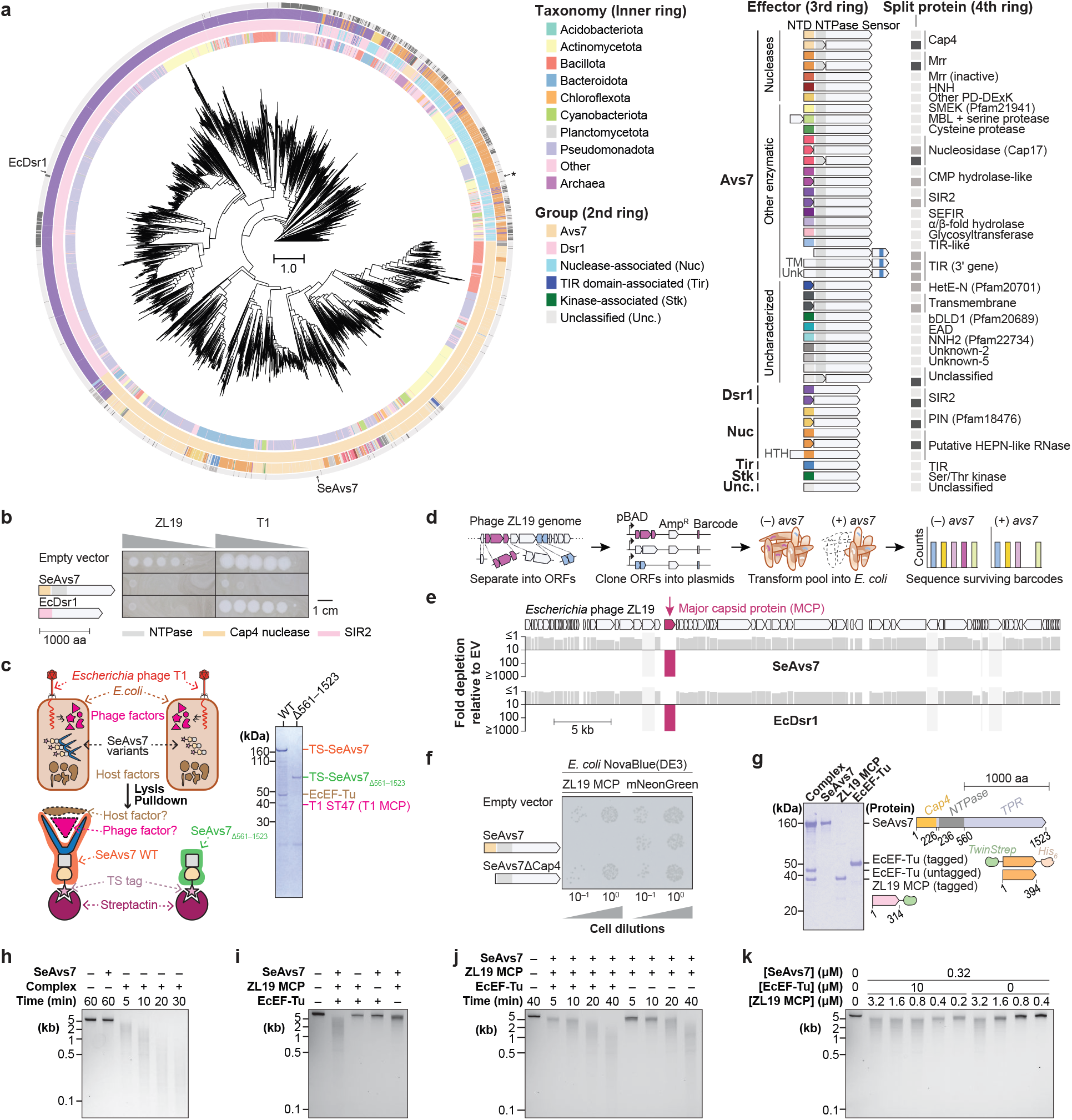
Homologous sensor domains in Avs7 and Dsr1 recognize phage major capsid proteins. **a**, Maximum likelihood tree of the homologous sensor domains of Avs7, Dsr1, and other defense proteins (*n* = 4,969 representatives). The inner-most ring represents taxonomy, and the three outer rings denote the type and domain architecture of the respective proteins. The asterisk indicates a HEPN-like RNase-containing protein from *Streptomyces sp*. NBC_00271. Unknown-2 is also fused to several Avs2 homologs. MBL, metallo-β-lactamase; EAD, effector-associated domain; TM, transmembrane; Unk, unknown. **b**, Plaque assays of *E. coli* expressing an empty vector, SeAvs7, or EcDsr1 against *Escherichia* phages ZL19 and T1. **c**, Schematic and SDS-PAGE analysis of an SeAvs7 affinity pulldown. TS, Twin-Strep tag. **d–e**, Synthetic lethal genetic screen to identify phage triggers that induce SeAvs7- and EcDsr1-mediated cell death. Candidate genes were cloned downstream of an arabinose-inducible pBAD promoter with a unique 20-nt barcode. AmpR, ampicillin resistance gene. **f**, Representative photographs of E. coli transformation assays. **g**, SDS-PAGE analysis of purified recombinant proteins and the oligomeric complex. **h–k**, Agarose gel analysis of *in vitro* nuclease activity assays with a linear dsDNA substrate.

SeAvs7 conferred robust defense in *E. coli* against nine *Drexlerviridae* family coliphages, whereas EcDsr1 was active against only a few (Fig. 2b and Extended Data Fig. 2b). Proteomic profiling of tagged SeAvs7 isolated from T1-infected cells revealed significant enrichment of the T1 major capsid protein (MCP; T1 ST47) and host translation elongation factor Tu (EcEF-Tu) bound to SeAvs7, compared to a sensor-deleted mutant (Fig. 2c, Extended Data Fig. 2c–f, and Supplementary Table 9). A co-expression toxicity screen using a barcoded library of 73 genes from phage ZL19 identified the MCP as the unique activator of both SeAvs7- and EcDsr1-mediated cell death (Fig. 2d–e and Supplementary Table 10), and direct coexpression of ZL19 MCP with SeAvs7 in *E. coli* caused pronounced cell death (Fig. 2f). Together, these results establish MCP as the trigger for Avs7 sensors across distinct defense systems.

### The SeAvs7–MCP complex incorporates elongation factor Tu

To investigate the activation of SeAvs7, we recombinantly purified SeAvs7 and ZL19 MCP and incubated them *in vitro* (Supplementary Fig. 6a). Size-exclusion chromatography (SEC) demonstrated that, in the presence of ATP and Mg^2+^, these proteins formed a large multimeric complex (Fig. 2g and Supplementary Fig. 6b). This complex efficiently degraded double-stranded DNA (dsDNA) substrates, whereas SeAvs7 alone did not (Fig. 2h). EcEF-Tu co-purified with SeAvs7 even in the absence of phage infection and co-eluted with the SeAvs7–ZL19 MCP complex during SEC (Supplementary Fig. 6b). To rule out nonspecific binding, we reconstituted the complex *in vitro* using individually purified SeAvs7, ZL19 MCP, and EcEF-Tu proteins (Fig. 2g and Supplementary Fig. 6c–d). SEC analysis confirmed that EcEF-Tu was stably incorporated into the multimeric complex at an approximately 1:1:1 molar ratio, demonstrating its role as a core structural component (Supplementary Fig. 6e).

To clarify the role of each protein, we assessed nuclease activity across different combinations of the individually purified proteins (Fig. 2g and Supplementary Fig. 6c). DNA degradation strictly required both SeAvs7 and ZL19 MCP and was substantially enhanced by EcEF-Tu (Fig. 2i). Although cleavage occurred without EcEF-Tu after prolonged incubation, its presence notably increased the sensitivity of SeAvs7 to ZL19 MCP (Fig. 2j–k and Extended Data Fig. 2i, and Supplementary Fig. 7). Given the high cellular concentration of EF-Tu (comprising 6−10% of the total *E. coli* proteome^36^) and its sequence conservation between *E. coli* and *Salmonella enterica* (a single amino acid difference), this enhancer role is likely conserved in the native host. The elimination of both defense and cellular toxicity of SeAvs7 by inactivation of the Cap4 nuclease indicates that the defense activity of SeAvs7 is not driven by EF-Tu sequestration (Extended Data Fig. 2g–h).

### SeAvs7 assembles into an asymmetric tetramer upon MCP binding

To understand how ZL19 MCP and EcEF-Tu interact with SeAvs7, we determined the structures of the oligomeric complex by cryo-electron microscopy (cryo-EM) (Supplementary Fig. 8, and Supplementary Table 11). Refinement revealed two distinct structures: an approximately 1 MDa butterfly-shaped asymmetric tetrameric complex at 3.43 Å resolution containing four copies each of SeAvs7, ZL19 MCP, and EcEF-Tu, and a smaller ∼250 kDa complex at 3.40 Å resolution with a single copy of each protein (Fig. 3a–b). The protomers of the tetramer are structurally similar except for the orientation of the SeAvs7 Cap4 nuclease domain (Fig. 3a and Extended Data Fig. 3a). In contrast to the tetramer, the smaller monomeric complex lacked density for the Cap4 nuclease and the N-terminal part of the NTPase, suggesting that these regions are flexible in the monomeric state but are stabilized upon tetramerization (Fig. 3b). The remaining parts of the monomeric complex closely resemble a protomer of the tetramer (Extended Data Fig. 3a).

**Figure 3.**
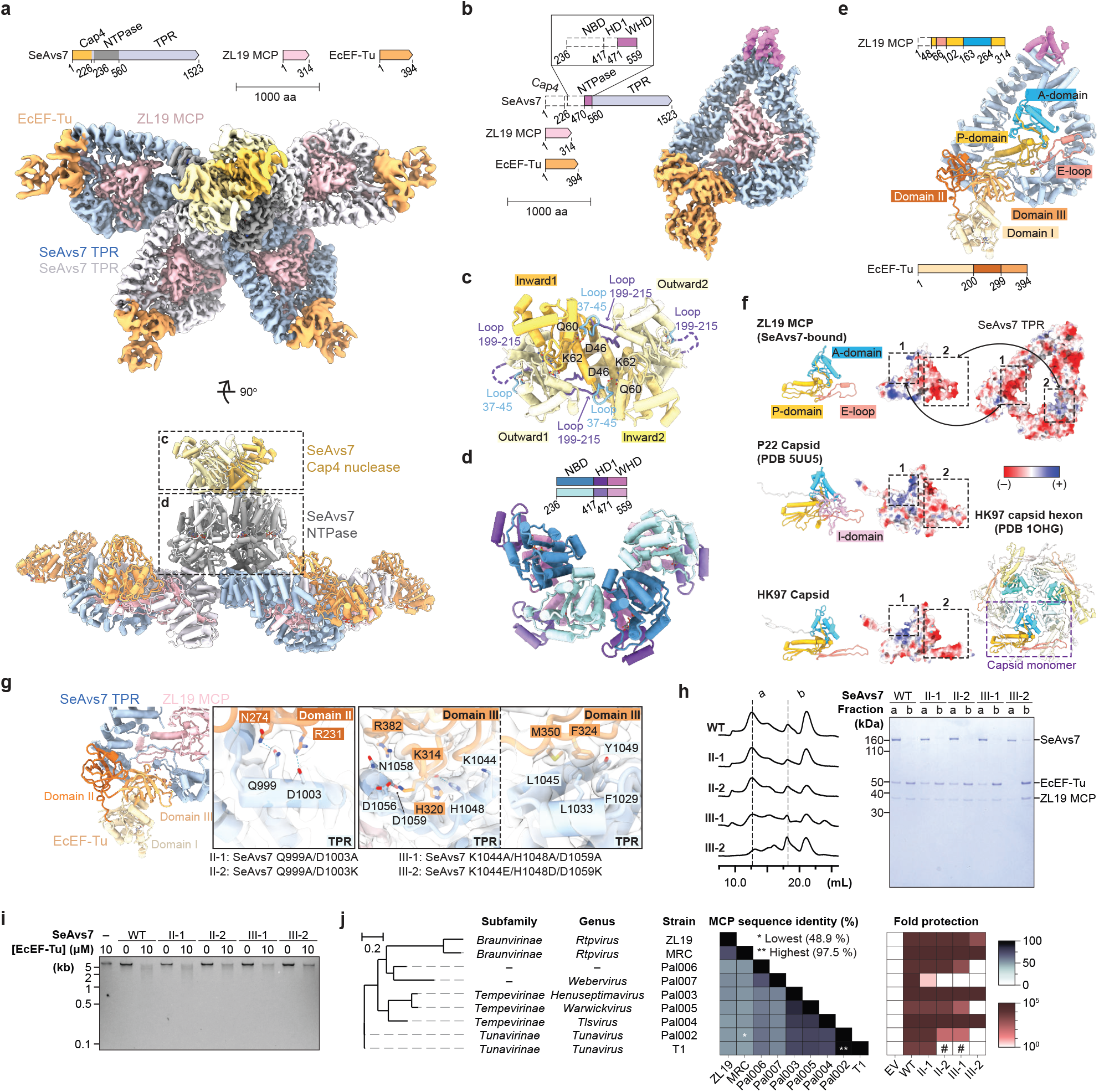
EcEF-Tu enhances SeAvs7 complex assembly and the immune response. **a**, Composite map (top) and structure of the tetrameric SeAvs7–ZL19 MCP–EcEF-Tu complex. **b**, Cryo-EM map of the monomeric complex. **c–d**, Structures of the Cap4 nuclease (**c**) and NTPase tetramers (**d**). **e**, Structure of the monomeric SeAvs7–ZL19 MCP–EcEF-Tu complex. **f**, Structural comparison of major capsid proteins (MCPs) from diverse tailed phages, alongside the electrostatic surface potentials of the MCPs and the SeAvs7 TPR domain. **g**, Interface between the SeAvs7 TPR domain and EcEF-Tu domains II–III. Mutants: II-1, SeAvs7^Q999A/D1003A^; II-2, SeAvs7^Q999A/D1003K^; III-1, SeAvs7_K1044A/H1048A/D1059A_; III-2, SeAvs7_K1044E/H1048D/D1059K_. **h**, Sizeexclusion chromatograms of the *in vitro* assembled complex and SDS-PAGE analysis of selected fractions. **i**, Agarose gel analysis of the dsDNA nuclease activity of the wild-type complex and mutants. **j**, Phylogeny of the nine *Drexlerviridae* phages evaluated (left), pairwise sequence identities of their respective MCPs (middle), and corresponding anti-phage activity (right). The hash (#) indicates reduced plaque size without a change in the efficiency of plating. EV, empty vector; WT, wild-type.

The Cap4 nuclease domains mediate tetramerization by forming a C2-symmetric arrangement with two inward- and two outward-facing protomers (Fig. 3c). The inward protomers contact each other through helix–helix interfaces, whereas the outward protomers engage in close contacts with the inward ones, forming two hydrogen bonds and burying ∼12% (∼1300 Å^2^) of their solvent-accessible surface area (Extended Data Fig. 3b). In contrast, the Cap4 tetramer of SeAvs3 (ref.^5^) adopts a distinct arrangement and buries only ∼2% (∼280 Å^2^) of the surface area of its outward Cap4 protomers (Extended Data Fig. 3c). AlphaFold3 modeling placed dsDNA in the SeAvs7 Cap4 PD-(D/E)xK active site (Supplementary Fig. 9a), suggesting that the inward protomers are accessible for DNA binding and likely form the active nuclease. Finally, the Cap4 domain is regulated by a flexible loop (residues 41–60) in SeAvs3, whereas SeAvs7 uses a different extended loop (residues 199–215) for the regulation (Fig. 3c).

The overall asymmetry of the SeAvs7 complex is established through close contacts between the NTPase domains of adjacent protomers (Fig. 3d). Specifically, the nucleotide-binding (NBD) and winged-helix (WHD) subdomains of adjacent protomers interact across a ∼1,100 Å^2^ interface stabilized by three hydrogen bonds, whereas helical domain 1 (HD1) sits outside this primary interface (Extended Data Fig. 3d–e). Each protomer coordinates ATP and Mg^2+^ through conserved residues (Thr291, Lys292, Asp358, and Arg395) (Extended Data Fig. 3e), consistent with the active conformations of other characterized STAND NTPase oligomers^5,24,26,28^. Notably, although the NTPase tetramer could potentially accommodate a fifth subunit (Supplementary Fig. 9b), the tetramer is sufficient to fully scaffold the assembled Cap4 nuclease, and pentamers were not observed experimentally. These findings indicate that the asymmetric tetramer represents the fully assembled immune complex rather than a partial intermediate.

The SeAvs7 TPR sensor adopts a bilobed structure that envelops the ZL19 MCP, burying 3,700 Å^2^ of its 15,800 Å^2^ solvent-accessible surface (Fig. 3e). Binding is driven by the shape and charge complementarity between the sensor and the HK97-fold of MCP, including the peripheral (P) domain, axial (A) domain, and extended loop (E-loop; Fig. 3f)^37–39^. The sensor extensively engages with the conserved penton/hexon-forming region of the A-domain, while making surface contacts with the variable P-domain and E-loop (Fig. 3f)^37,38,40,41^. AlphaFold3 (ref.^42^) predictions for other Avs7–MCP pairs suggest this broad interface is a conserved feature of Avs7 sensors (Extended Data Fig. 5d and Supplementary Fig. 9c). Although we identified five residues potentially involved in hydrogen bond network formation, mutating three of these (N1204, Y1289, and N1361) did not abolish defense against ZL19, suggesting that recognition depends on extensive structural complementarity rather than on specific contacts (Extended Data Fig. 3f–g).

### EF-Tu binding enhances SeAvs7 complex assembly, nuclease activity, and defense

While ZL19 MCP is enveloped by the TPR sensor of SeAvs7, EcEF-Tu binds the outer surface of the flexible sensor region and adjacent helices through its tRNA-binding domains (Domain II and Domain III)^43^. This interaction buries ∼1,450 Å^2^ of solvent-accessible surface and is also facilitated by shape complementarity (Fig. 3g). Within this complex, EcEF-Tu adopts the open, inactive conformation typically bound to GDP (Extended Data Fig. 4a). Although the bound nucleotide could not be resolved, structural superposition indicates that the closed, GTP-bound active state would sterically clash with the TPR N-lobe (Extended Data Fig. 4b)^44^, suggesting SeAvs7 specifically co-opts the inactive form of EF-Tu.

The SeAvs7–EcEF-Tu interface consists of a network of contacting residues, including several that are conserved within one Avs7 clade (Extended Data Fig. 5a–b). Consistent with this observation, AlphaFold3 confidently predicted Avs7–MCP–EF-Tu complexes exclusively for members of this clade (Extended Data Fig. 5c–d and Supplementary Table 12). To test the functional importance of the interaction, we generated four SeAvs7 mutants targeting conserved charged residues (Q999, D1003, K1044, H1048, D1059; Fig. 3g). None of these mutants co-purified with detectable amounts of EcEF-Tu when expressed alone, indicating substantially reduced affinity (Supplementary Fig. 9d).

Upon reconstitution of the multimeric complex, mutants II-1 (SeAvs7^Q999A/D1003A^) and II-2 (SeAvs7^Q999A/D1003K^) formed EF-Tu-containing complexes at levels comparable to WT; mutant III-1 (SeAvs7^K1044A/H1048A/D1059A^) formed less complex; and mutant III-2 (SeAvs7^K1044E/H1048D/D1059K^) failed to incorporate a detectable amount of EcEF-Tu (Fig. 3h). EF-Tu-mediated enhancement of nuclease activity directly paralleled this assembly efficiency, with WT and mutant II-1 strongly enhanced, and mutants II-2, III-1, and III-2 showing progressively weaker effects (Fig. 3i). Plaque assays against nine *Drexlerviridae* family phages (with pairwise sequence identities of MCPs ranging from 48.9% to 97.5%) further corroborated the functional link, showing progressive loss of defense from mutant II-1 to III-2 against five phages, mirroring the trends in complex formation efficiency and nuclease activity enhancement (Fig. 3j and Extended Data Fig. 5e–f). The loss of defense activity by the most disruptive mutant (III-2) did not correlate with either MCP sequence similarity or phage phylogeny, indicating the promiscuity and variable dispensability of the Avs7–EF-Tu interaction across the diversity of phages (Fig. 3j). These findings establish EcEF-Tu as a key partner of Avs7 that optimizes phage defense by promoting the formation of the complex between Avs7 and MCP, thereby enhancing nuclease activity.

### MCP binding triggers structural rearrangements for SeAvs7 oligomerization

To understand the molecular basis of SeAvs7 activation, we determined the structure of apo SeAvs7 at 2.72 Å resolution (Fig. 4a, Supplementary Fig. 10, and Supplementary Table 11). Although EcEF-Tu copurified with SeAvs7, no corresponding cryo-EM density was observed, suggesting weak association in the absence of MCP (Supplementary Fig. 10a). Compared to the assembled tetramer, the apo structure shows a ∼90° rotation of the NBD and HD1 relative to the WHD, with none of these domains resolved in the monomeric intermediate (Fig. 4b). Notably, despite ATP being supplied during purification, ADP occupies the NTPase active site and stabilizes a closed conformation (Extended Data Fig. 6a). In contrast, the active tetramer exclusively contains ATP, suggesting ADP-to-ATP exchange during activation. Recognition of both nucleotides involves largely the same residues, except that an Arg395–ATP γ-phosphate contact replaces a His545–ADP β-phosphate contact (Extended Data Fig. 6a). Consistent with this observation, DNA cleavage was slower in the presence of ADP or the nonhydrolyzable ATP analog adenylyl-imidodiphosphate (AMP-PNP), and absent without a nucleotide, indicating that nucleotide binding promotes activation through a conformational change rather than through ATP hydrolysis (Extended Data Fig. 6b).

**Figure 4.**
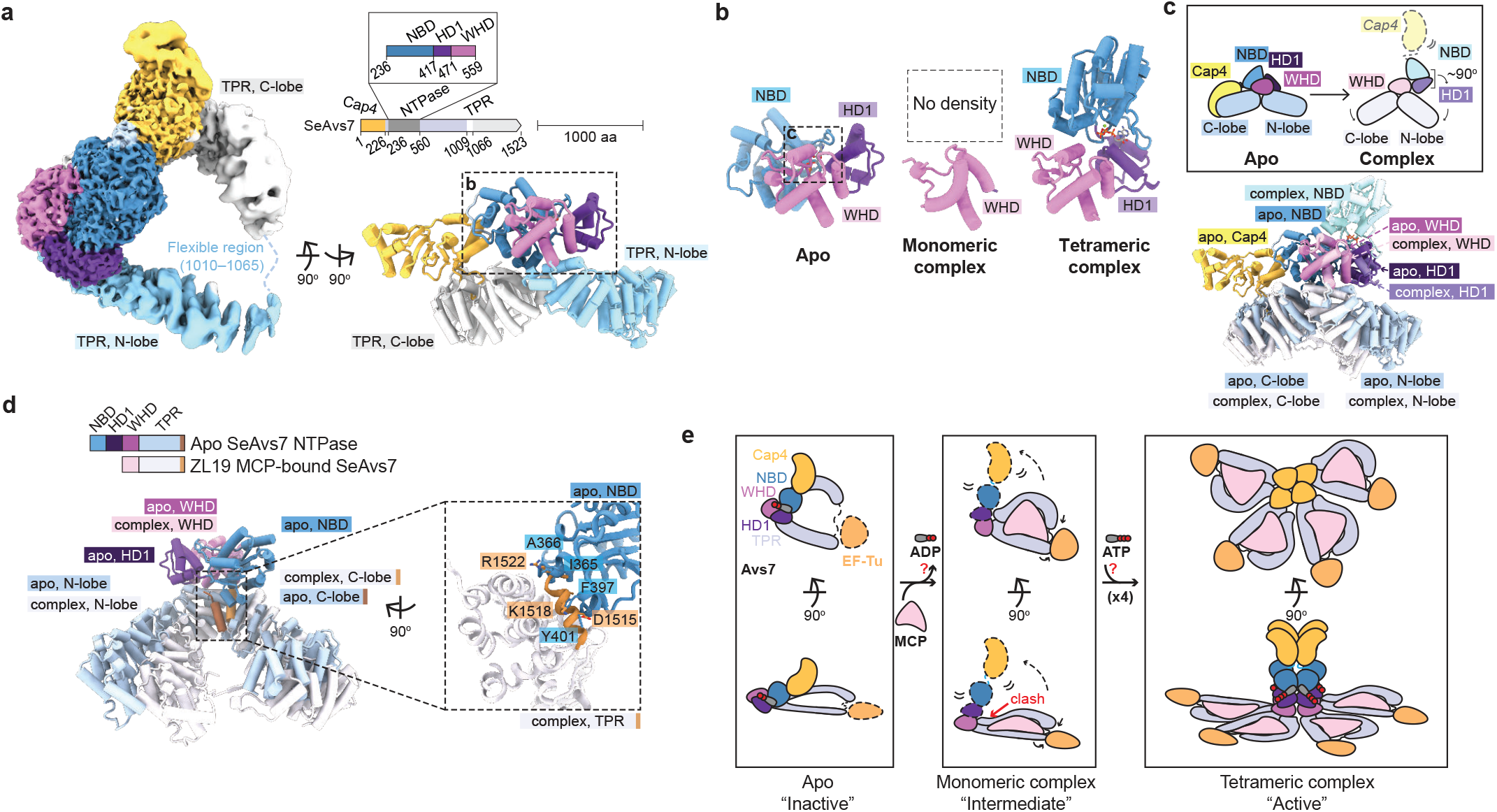
MCP binding induces large structural changes in SeAvs7. **a**, Cryo-EM map (left) and structure (right) of apo SeAvs7. **b**, Comparison of the NTPase domains from three structural states. **c**, Overlay of apo SeAvs7 and a tetrameric complex protomer aligned by the WHD. A schematic of the structural rearrangement is shown in the top inset. **d**, Alternate view of the WHD-aligned overlay from (c), highlighting the steric clash between the C-terminal helix of MCP-bound SeAvs7 (orange) and the NBD of apo SeAvs7 (blue). **e**, Proposed mechanism of SeAvs7 activation during phage infection.

The apo state of SeAvs7 is maintained by extensive autoinhibitory interactions. The closed conformation of the NTPase is stabilized by its interaction with the C-lobe through charge complementarity and a network of hydrogen bonds around the partially conserved Arg367 (Extended Data Fig. 6c–d). The Cap4 nuclease domain is sequestered by extensive hydrogen bond contacts with the NBD and by engaging the C-lobe (Extended Data Fig. 6d–e). Superimposing the assembled Cap4 tetramer onto the apo structure reveals severe steric clashes with the apo C-lobe, indicating that tetramerization is incompatible with the apo state (Extended Data Fig. 6f).

Binding of ZL19 MCP triggers global conformational changes that relieve autoinhibition of SeAvs7. Superimposing the apo and tetramer structures by the WHD shows that MCP binding causes the TPR N- and C-lobes to clamp together, displacing them by ∼30 Å relative to each other (Fig. 4c–d and Extended Data Fig. 6g). This movement leads to steric clashes between the MCP-bound C-lobe and the NTPase helices of the apo form, likely disrupting the interactions that stabilize the inactive state (Fig. 4d). These include a notable clash between the C-terminal helix of the C-lobe (residues 1511–1521) and two NBD helices (residues 360–367 and 396–401). The Cterminal helix is present across Avs7 homologs and has a conserved length (Extended Data Fig. 6h).

Based on these observations, we propose a stepwise mechanism for the activation and assembly of Avs7 complexes (Fig. 4e). In the absence of MCP, Avs7 remains in an autoinhibited, ADP-bound resting state. Upon phage infection, the Avs7 sensor binds a newly synthesized MCP monomer, strengthened by EF-Tu, clamping the two lobes of the sensor. These conformational changes destabilize the ADP-bound state of the NTPase, likely promoting ADP release and increased flexibility of the N-terminal domains. Finally, ATP binding stabilizes the NTPase, driving tetramerization and enabling Cap4-mediated DNA cleavage to prevent phage replication.

### Thirteen additional Avs families are activated by diverse phage proteins

To investigate the mechanisms of diverse STAND families beyond Avs7, we cloned representative proteins from 30 additional sensor structural clusters (Extended Data Fig. 1d–f, Supplementary Fig. 2b, and Supplementary Table 6), focusing on *Enterobacteriaceae* homologs, when possible, to facilitate experimental reconstitution in *E. coli*. To identify triggers, we conducted a genetic screen with a barcoded plasmid library of 600 genes from 36 diverse phages and viruses, including ssRNA, ssDNA, and tailed and tail-less dsDNA phages (Supplementary Table 10). We transformed the library into STAND-expressing *E. coli* and deep-sequenced the barcodes to quantify cell death triggered by individual phage genes (Fig. 5a).

Thirteen STAND families, which we denote Avs8 and Avs10–21 (because Avs6 and Avs9 now refer to other SAMD9-like families^33^), strongly induced cell death in response to specific phage proteins (Extended Data Fig. 7 and Supplementary Fig. 11). Notably, the identified targets encompassed most of the core structural and replicative proteins of tailed phages^45^ (Fig. 5b). Avs8 (PD-λ-4)^14^ and Avs10 (Erebus/Hypnos/bNACHT64)^16,17^ sensed MCPs (consistent with recent findings for Avs8 (ref.^46^)), whereas Avs12–18 recognized seven distinct structural proteins: portal adaptor, tail nozzle, head-tail connector, tail terminator, tail tube protein (TTP), tail assembly chaperone (TAC), and tape measure protein (TMP), respectively (Extended Data Fig. 7). In contrast, Avs19–21 sensed replicative proteins: family A DNA polymerases, the RecA-type ATPase domain in both the primase-helicase and RecA, and single-stranded DNA annealing protein (SSAP), respectively (Extended Data Fig. 7). Avs11 recognized the portal protein despite lacking sequence and structural similarity to the previously characterized portal-sensing Avs4 (Extended Data Fig. 7 and Supplementary Fig. 2). In all cases, mutating or deleting the effector domains abolished cell death (Supplementary Fig. 12).

**Figure 5.**
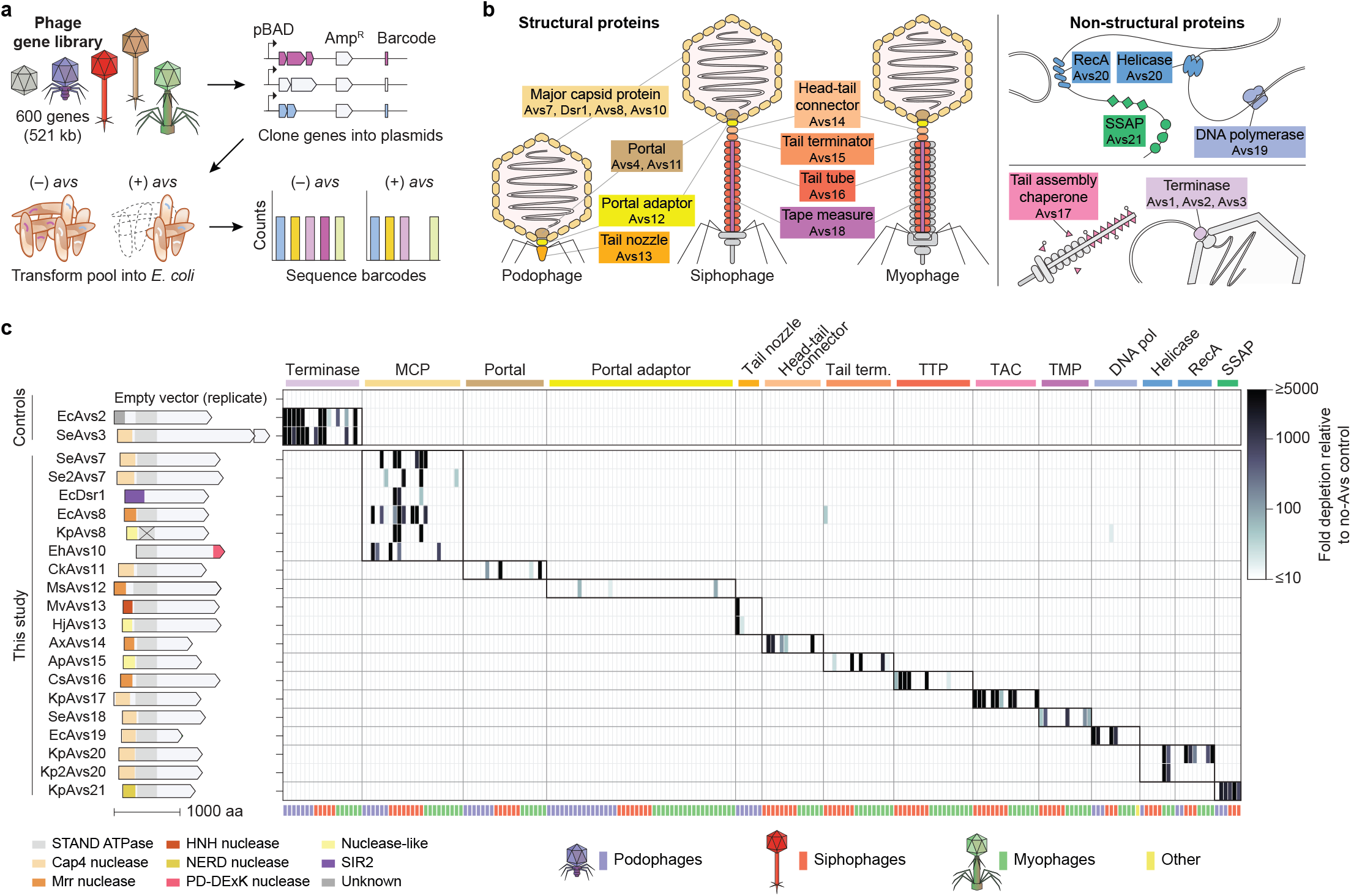
Thirteen additional Avs families are activated by core phage structural and replicative proteins. **a**, Avs activation screen using an expanded plasmid library expressing 600 genes from 36 phages and archaeal viruses (see Extended Data Fig. 7). **b**, Functional roles of phage proteins recognized by Avs families. **c**, Heatmap of Avs-mediated toxicity induced by diverse cognate target proteins. MCP, major capsid protein; TTP, tail tube protein; TAC, tail assembly chaperone; TMP, tape measure protein; SSAP, ssDNA annealing protein.

To evaluate the specificity of the identified interactions, we assembled a panel of 218 target homologs, combining hits from the genetic screen with 60 newly cloned additions to ensure at least 6 diverse homologs per target protein. We quantified all 4,360 pairwise interactions between this panel and 18 Avs representatives via deep sequencing, and selectively tested an additional 27 genes (Supplementary Fig. 14). Consistent with the initial screen, each Avs was triggered by homologous proteins from different phages, with minimal cross-reactivity against non-cognate targets (Fig. 5c). Within each target family, Avs proteins recognized a broad range of homologs, in many cases with little to no sequence similarity, but sharing the same core fold (Extended Data Fig. 8 and Supplementary Fig. 15). For instance, Avs21 could detect both ERF-family and RecT-family SSAPs, which have virtually no sequence similarity^47^. The conservation of Avs target proteins mirrors the overall synteny of structural genes across phage genomes (Supplementary Fig. 13).

Supporting these findings, most tested Avs families mediated anti-phage defense in *E. coli* against at least one of 12 tested coliphages. Specificities generally aligned with their identified genetic triggers, with a few exceptions (Extended Data Fig. 9a–b and Supplementary Fig. 16). Anti-phage activity was also detected for four additional sensor structural clusters (Avs22–24 and bNACHT20) for which no phage-encoded activators were identified in our screen (Extended Data Fig. 9a and Supplementary Fig. 16). Further investigation is needed to determine how these Avs families sense phage infection.

### Avs proteins recognize core phage structural patterns

To determine how Avs8 and Avs10–21 engage their targets, we generated AlphaFold3 or AlphaFold-multimer models, obtaining confident predictions (ipTM 0.71–0.91) for 11 of the 13 complexes (Fig. 6, Extended Data Fig. 10, and Supplementary Fig. 17). In each case, the Avs TPR-like sensor bound a single copy of the target protein through an extensive, shape-complementary interface. *In vitro* reconstitution of two complexes (KpAvs20–HdK1 RecA and MvAvs13–PhiV-1 tail nozzle) biochemically confirmed direct binding and subsequent Avs oligomerization (Fig. 6f, m).

**Figure 6.**
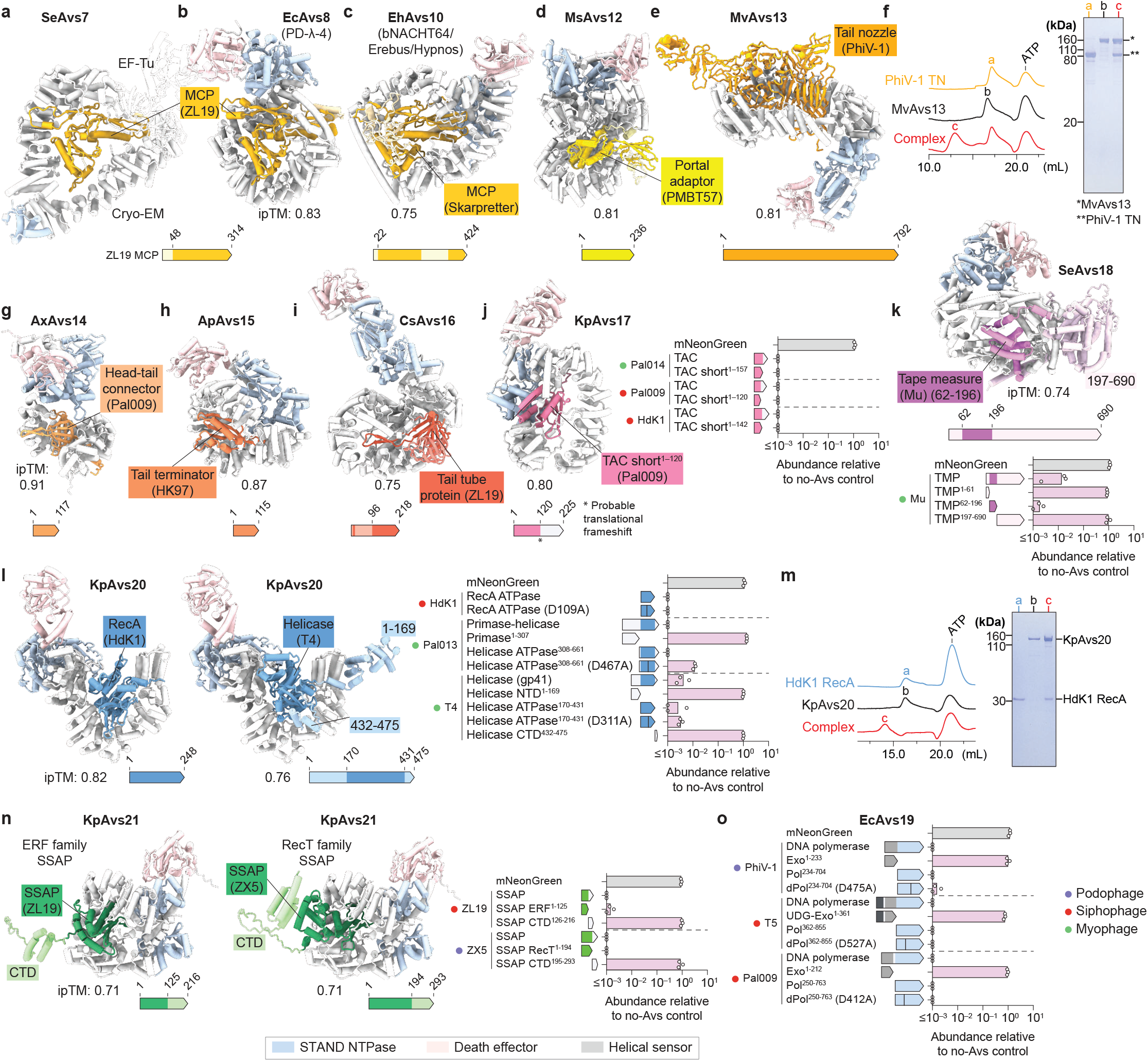
Avs proteins recognize core structural patterns through shape complementarity across extended binding interfaces. **a–n**, AlphaFold2/3 models of Avs proteins binding the major capsid protein (MCP) (**a–c**), portal adaptor (**d**), tail nozzle (**e**), head-tail connector (**g**), tail terminator (**h**), tail tube protein (**i**), tail assembly chaperone (**j**), tape measure protein (**k**), helicase/RecA-fold ATPase (**l**), and ssDNA annealing proteins (SSAPs) (**n**), accompanied by quantification of Avs activation by truncated or catalytically inactivated target proteins. Bars represent the mean of three barcode replicates. **f, m**, Size-exclusion chromatograms and SDS-PAGE analysis of MvAvs13–PhiV-1 tail nozzle (**f**) and KpAvs20–HdK1 RecA (**m**) complexes are shown. **o**, EcAvs19 activation by truncated or catalytically inactivated DNA polymerases. ipTM, interface predicted template modeling score (0.0–1.0); TN, tail nozzle. ipTM values refer to protein–protein pairs only and exclude interactions with ligands (*e.g*., ATP and Mg^2+^).

The Avs sensors consistently recognized the conserved core folds of their targets, leaving variable extensions unbound (Extended Data Fig. 8, 10, and Supplementary Fig. 15). For example, despite their different sensor architectures, Avs7, Avs8, and Avs10 all bind the penton/hexon-forming interface of the MCP A-domain (Fig. 6a–c). Avs12–15 envelop the globular folds of their single-domain targets^48^ (Fig. 6d–e, g–h). Avs16 forms extensive contacts with the conserved β-hairpin of the TTP, which constitutes the interface between stacked hexameric rings of the tail tube (Fig. 6i). Avs17 binds the N-terminal core of the TAC, a region shared by both long and short isoforms generated through programmed translational frameshifting^49^ (Fig. 6j). For SeAvs18, models indicate binding to a central ∼130-residue α-helical segment of TMPs, which PROMALS3D^50^ independently identified as the most conserved region. This segment was necessary and sufficient to activate cell death (Fig. 6k). Core fold recognition extended to replicative proteins: Avs20 binds the RecA-type ATPase domain that is present in both primase-helicase and RecA^51^, independent of ATPase catalytic activity (Fig. 6l), and Avs21 recognizes the N-terminal ssDNA-binding domain of both ERF-family and RecT-family SSAPs (Fig. 6n).

Although high-confidence models could not be generated for CkAvs11–portal and EcAvs19–DNA polymerase complexes, predictions of apo Avs proteins suggest autoinhibition that might be released by target binding (Extended Data Fig. 9c–d). The core polymerase domain was necessary and sufficient to activate EcAvs19, independent of the polymerase catalytic aspartate (Fig. 6o).

Altogether, our structural modeling and mutational analyses demonstrate that Avs families recognize the conserved core structures of highly divergent target proteins (Fig. 6, Extended Data Fig. 10, and Supplementary Fig. 16). This recognition appears to target monomers, as structural modeling showed major steric clashes with their oligomerized or substrate-bound forms (Supplementary Fig. 18).

## Discussion

Here we present a systematic study of the diversity and activation of STAND defense systems across prokaryotes. Together with the previously characterized terminase-sensing Avs1–3 (ref.^5^), our findings demonstrate that dedicated Avs families monitor most of the conserved core proteins of tailed phages. At least 12 of the 14 investigated families engage their targets through broad shape-complementary interfaces that envelop the target core, a mechanism that distinguishes Avs sensors from most other known protein-sensing defense systems. For instance, three prevalent families (Avs7, Avs8, and Avs10) wrap around the HK97-fold MCP, contrasting systems such as Lit protease^52,53^ and CapRel^54,55^, which detect MCP through short sequence motifs.

Six of the 13 identified targets—tail nozzle, portal adaptor, head-tail connector, tail terminator, tail assembly chaperone, and tape measure protein—have not been previously known to trigger defense. For three replicative proteins—primase-helicase, RecA, and SSAP—phage escape mutations have been previously identified^56-59^, but the mechanisms of sensing remained unknown and could be indirect. For instance, the Hachiman/AbpAB defense system detects genome integrity rather than replicative proteins themselves^60^. Altogether, our findings substantially expand the repertoire of phage proteins that directly activate prokaryotic defense systems.

Our structural and biochemical analyses of SeAvs7 reveal a previously unknown mechanism of host factor repurposing in anti-phage defense. The structural role of EF-Tu in the Avs7 complex contrasts with its involvement in the functioning of other defense systems such as Lit protease^61-63^ and type IV Thoeris^64^ systems, where EF-Tu is cleaved to arrest translation. Instead, this mechanism is reminiscent of the ribosomal protein S15 in the type V-K CRISPR-associated transposon (CAST) complex^65^, which facilitates DNA mobilization. Another relevant case of EF-Tu repurposing is its recruitment (along with EF-Ts) as an accessory subunit by the RNA-dependent RNA polymerase of *Leviviridae* family phages^66,67^. By co-opting highly abundant and conserved host proteins, both defense systems and mobile elements appear to achieve a reduction in their own protein investment without substantially perturbing the host proteome^68^. These observations parallel those for several eukaryotic NLRs that bind host factors, undergoing spatial or conformational changes upon infection^27,69^. Alternatively or additionally, given that T4 MCP binds EF-Tu^63^, Avs7 might have evolved to exploit a pre-existing interaction between certain MCPs and EF-Tu, a mechanism that parallels sensing of DnaJ perturbation by bNACHT25 (ref.^8^).

Our findings do not exclude the existence of other, noncanonical activating mechanisms for the Avs families described here. Indeed, alternative activators have been identified for CapRel^55^ and Avs2 (ref.^6^). Avs activation also might require additional host factors beyond their phage target, as exemplified by EF-Tu recruitment by Avs7 and DnaJ perturbation by bNACHT25 (ref.^8^). Further studies are required to elucidate the full repertoire of these activation mechanisms. Nonetheless, the conservation and broad structural complementarity of the identified targets with the Avs sensor domains suggest these interfaces are favored during the host-virus arms race, enabling sensing of a broad range of phages while avoiding autoimmunity.

Beyond the 13 targets identified here, the full sensory repertoire of prokaryotic STANDs remains an open question. Additional STAND families might detect other essential phage components that are more variable or lineage-specific, such as small terminase subunit, tail sheath, minor tail proteins, and baseplate hub, or other uncharacterized proteins^6,7,9^. Furthermore, it remains a possibility that some sense molecules other than proteins. Our phylogenetic analysis and structural classification of the STANDs provide a resource for future investigation.

In addition to their fundamental role in phage defense, the mechanism of Avs sensors offers insights for therapeutic innovation. The discovery that bacterial defense systems actively monitor conserved, essential phage structures exposes potential vulnerabilities that could be exploited for off-the-shelf therapeutic phage design. Conversely, the finding that small phage proteins can trigger Avs-mediated cell death suggests the possibility of developing novel antimicrobials that artificially activate these pathways in antibiotic-resistant pathogens. Altogether, our findings reveal that structure-based pattern recognition of the core phage proteome is a major defense strategy in prokaryotes, highlighting sophisticated adaptations in the host-virus arms race.

## Materials and Methods

### Bioinformatics analysis of STAND NTPases

To systematically identify defense-related STAND NTPases in prokaryotes, we curated an initial set of 656 diverse STAND ‘seed’ sequences (Supplementary Table 1). This set included representatives of all previously predicted prokaryotic defense-related STANDs^2,4,16–18^, covering both NACHT and other clades. We also included additional sequences from GenBank identified through BLAST searches using the known STANDs and the defense-associated effector proteins Cap4, Mrr, Trypco2, and metallo-β-lactamase^5^.

The 659 NTP-binding domains within the 656 seed sequences were clustered, aligned, and used as queries in PSI-BLAST^70^ searches against the NCBI clustered_nr database (non-redundant proteins clustered at 90% identity and 90% coverage) to identify a preliminary set of homologs. Footprints of these searches were then aligned with the corresponding queries and used in a second round of PSI-BLAST searches against the same database, resulting in the identification of 1,750,195 NTP-binding domains from 1,708,335 proteins. These NTP-binding domains, originating from both prokaryotes and eukaryotes, were clustered using MMSeqs2 (ref.^71^) at 75% identity and 67% coverage, yielding 680,395 clusters.

Representatives of these 680,395 clusters (one from each cluster) were selected and further clustered using MMSeqs2 at 33% identity and 67% coverage, producing 37,834 clusters. Sequences within each new cluster were aligned using MUSCLE5 (ref.^72^) and the resulting 37,834 consensus sequences^73^ were aligned using FAMSA^74^. Each character of the aligned consensus sequence was then expanded into a corresponding column of the original cluster MSA, generating a pseudo-MSA of 680,395 representative NTP-binding domain sequences.

A preliminary approximate maximum likelihood tree was reconstructed from these sequences using FastTree^75^ with gamma-distributed site rates and the WAG evolutionary model. This tree was cut at a depth of 1.5 from the tips into 17,857 clades. The domain content of the 680,395 sequences across these clades were annotated using PSI-BLAST against the NCBI CD database^76^; clades containing potential defense-related NTPases were identified by the presence of likely sensor and effector domains. This procedure identified 884 potential defense-related clades, totaling 359,879 leaves corresponding to NTP-binding domain sequences. The discarded clades mainly consisted of homologous but distantly related P-loop domains identified by PSI-BLAST, such as ABC transporters, adenylate kinases, and Clp proteases, or were singleton outliers.

This subset of 359,879 leaves was further clustered using MM-Seqs2 at 25% identity and 67% coverage, resulting in 2,488 clusters. Each cluster was aligned using MUSCLE5, and the 2,488 consensus sequences were aligned using FAMSA. Each character of the aligned consensus sequence was expanded into a corresponding column of the original cluster MSA, resulting in a pseudo-MSA of 359,879 representative NTP-binding domain sequences. A final approximate maximum likelihood tree was reconstructed from these sequences using FastTree with gamma-distributed site rates and the WAG evolutionary model. The tree was cut at a depth of 2.0 from the tips, yielding 589 final STAND clades (Supplementary Table 2). The tree was visualized using ggtree v3.10.1 in R (ref.^77^).

Each of the 589 STAND clades were classified as defense-associated if either or both of the following criteria were met: the presence of defense effector domains within the protein sequences or statistical enrichment of genomic context adjacent to known defense systems.

- The presence of effector domains was assessed using the following criteria: Clades in which at least 50% of the representative sequences contained previously annotated effector domains (detected by PSI-BLAST using CDD profiles; Supplementary Table 3) were classified as high-confidence defense clades (86 clades), and those with 10–50% of sequences containing these domains were considered low-confidence defense clades (86 clades).
- To assess the genomic context of each STAND clade, we used DefenseFinder^78^ to identify known anti-phage defense genes encoded within 5 kb of representative STAND sequences. For each clade, to determine whether the observed proximity to defense genes was significantly greater than expected by chance, we calculated a *P*-value using the binomial distribution. The null probability was set at 0.0153, which reflects the chance that a randomly selected non-defense gene would be located within 5 kb of a defense system, based on our calculated DefenseFinder statistics. Clades with a *P*-value less than or equal to 0.0001 were classified as high-confidence defense-associated clades, resulting in 131 such clades (Supplementary Fig. 1b).

This combined classification scheme yielded 198 highconfidence defense-associated STAND clades (Supplementary Table 4). To assess sensor diversity, sequence clusters within these clades (clustered at 75% identity and 67% coverage; see Fig. 1a, step 3) were expanded, resulting in 178,641 proteins. From these, C-terminal regions following the NTPase domains were extracted, and sequences shorter than 400 amino acids were excluded, yielding 34,990 sequences. These sequences were then clustered using MMseqs2 (ref.^71^) at 80% coverage and an *E*-value threshold of 10^−5^, resulting in 1,479 sequence clusters.

For each sequence cluster, a representative AlphaFold structure was either retrieved from the AlphaFold database (https://alphafold.ebi.ac.uk/)^79,80^ or predicted using AlphaFold3 (ref.^42^), using the full protein sequence along with one ADP molecule and one Mg^2+^ ion. Seventeen eukaryotic clusters were excluded, and five clusters did not yield structural models. From the remaining 1,457 representative structures, the C-terminal regions following the NTPase domains were extracted and clustered using Foldseek^81^ at 80% coverage, an *E*-value of 10^−10^, and a TM-score threshold of 0.40, resulting in 684 structural clusters (Supplementary Table 5). For visualization, 90 sensor structural clusters were selected, based on (1) each cluster containing at least five members, and (2) at least 25% of cluster members belonging to defense-associated STAND clades.

### Prophage prediction

Prophage regions were detected using PHASTEST^82^.

### Bioinformatics analysis of Avs7 and Dsr1

Avs7 homologs (Supplementary Table 7) were identified using PSI-BLAST searches against the NCBI nonredundant (nr) protein sequence database in July 2024. After filtering out truncated sequences and non-Avs7 hits, a final set of 3,272 verified Avs7 sequences was obtained. These sequences were then clustered at 90% identity using MMseqs2 (ref.^71^), resulting in 1,757 clusters, and one representative from each cluster was selected for subsequent analysis. A multiple sequence alignment (MSA) of these representatives, excluding their variable N-terminal domains, was generated using MAFFT v7.520 (ref.^83^) with global pairwise alignment (--maxiterate 1000 --globalpair). The alignment was trimmed with trimAl 1.2 (ref.^84^) using a gap threshold of 0.25 (-gt 0.25). Phylogenetic trees were constructed from the trimmed MSAs using IQ-TREE 1.6.12 (ref.^85^) with the LG+G4 model and 2000 ultrafast bootstrap replicates (parameters -nstop 500 -safe -nt 4 -bb 2000 -m LG+G4). N-terminal effectors were annotated based on HHpred^86^ predictions, and taxonomic classifications were retrieved from the NCBI taxonomy database. The tree was visualized using iTOL^87^.

To identify additional defense proteins that share the Avs7 sensor domain, PSI-BLAST searches were performed against the NCBI nr protein sequence database downloaded in February 2024. The sensors from two Avs7 representatives (OJX47172.1 and WP_079981830.1) were used as queries for ten and nine iterations, respectively, with a maximum *E*-value threshold of 0.001 and a minimum sequence coverage of 70%. These searches yielded an initial list of 10,249 proteins. Given the high number of Dsr1 homologs in this list, we conducted an additional PSI-BLAST search using the sensor domain of EcDsr1 (WP_029488749.1) for 16 iterations, which added 45 unique proteins.

A round of curation to remove partial proteins and false positives yielded 9,097 proteins containing an Avs7 sensor domain (Supplementary Table 8). These sensor domains were extracted and clustered at 95% sequence identity and 98% coverage using MMseqs2 (ref.^71^). An MSA of 4,969 representative sequences was generated using MAFFT v7.525 (ref.^83^) with global pairwise alignment. The alignment was trimmed by trimAl 1.5 (ref.^84^) with a gap threshold of 0.25. A phylogenetic tree was constructed from the trimmed alignment using IQ-TREE 2.3.6 (ref.^88^) with the LG+G4 model and 2000 ultrafast bootstrap replicates. Protein domains were predicted using HHpred^86^, and for split proteins, adjacent sequences were also analyzed. Taxonomic classifications were retrieved from the NCBI taxonomy database, and the tree was visualized using the R package ggtree v3.10.1 (ref.^77^).

### Cloning

Genes were chemically synthesized (Twist Bioscience or Integrated DNA Technologies) or amplified by polymerase chain reactions from genomic DNA using Q5 (New England Biolabs) or Phusion Flash (ThermoFisher) polymerases. Plasmids were constructed with Gibson assembly and transformed into *E. coli* strains Stbl3 or Mach1 (ThermoFisher). *E. coli* colonies were grown in 1.8 mL Terrific Broth overnight at 37 °C with 100 µg/mL carbenicillin or 25 µg/mL chloramphenicol, as appropriate. Plasmids were isolated using Qiagen miniprep buffers and 96-well DNA & RNA binding plates (Epoch Life Science). The full sequences of all plasmids were verified with Tn5 tagmentation and Illumina sequencing, as previously described^4,89^.

STAND NTPases and mutants were cloned into a low-copy pACYC184-like plasmid backbone with a chloramphenicol resistance gene (Supplementary Table 6). For genes from *Enterobacteriaceae*, the native promoter and native DNA sequence was retained in most cases (66 systems), with the exception that the lac promoter was used if the gene was separated by less than 50 bp from the preceding open reading frame (ORF) in the genome, or if the promoter could be not synthesized (4 systems). Genes outside of *Enterobacteriaceae* (40 systems) were codon optimized for expression in *E. coli* K-12 and cloned after a lac promoter.

Phage genes and mutants were cloned downstream of an arabinose-inducible pBAD promoter into a pBR322-like plasmid backbone with an ampicillin resistance gene and a 20 bp barcode sequence unique to each clone. One to three clones for each construct, each with a distinct barcode, were retained for inclusion in the phage gene library (Supplementary Tables 10 and 13).

### Phage samples

Phages T4, T5, PhiV-1, P1, Lambda, M13, MS2, Qβ, PhiX, and PR772 were obtained from the American Type Culture Collection (ATCC). Phages HdK1, PTXU04, and PMBT57 were obtained from DSMZ. Phages Pal001–014 were isolated from wastewater treatment samples from Palo Alto, California. Wastewater samples were passed through a 0.22 µm syringe filter (Cytiva Whatman), 10^2^-, 10^3^-, or 10^4^-fold diluted in water, and plated on top agar containing *E. coli* K-12 strains ATCC25404 or BW25113 ΔiscR (ref.^90^), similar to phage plaque assays. After overnight incubation at 37 °C, phages from individual plaques were isolated from the top agar.

Phages used in plaque assays were amplified by incubation at 37 °C with *E. coli* K-12 (ATCC25404) or *E. coli* BW25113 in Terrific Broth at log phase (OD_600_ ∼ 0.2−0.5), without shaking, for up to 12 h or until lysis. Lysates were passed through a 0.22 µm syringe filter, and the filtrate was stored at 4 °C for subsequent use.

### Phage plaque assays

*E. coli* BW25113 was used in phage plaque assays in Fig. 2b and 3j, and Extended Data Fig. 2b, 2g, 3g, and 5e. *E. coli* K-12 (ATCC 25404) was used in phage plaque assays in Extended Data Fig. 9a–b and Supplementary Fig. 16, except for those related to Avs21, for which *E. coli* Stbl3 was used instead due to the toxicity of Avs21 expression in *E. coli* K-12 (ATCC25404). *E. coli* cultures were grown to saturation in Terrific Broth at 37 °C with orbital shaking at approximately 220 rpm. To 50 mL top agar (10 g/L tryptone, 5 g/L yeast extract, 10 g/L NaCl, 7 g/L agar) was added chloramphenicol (to a final concentration 25 µg/mL) and 1 mL *E. coli* culture, and the mixture was poured into an empty 15 cm Petri dish and left at room temperature for 10–30 min to solidify. Ten-fold serial dilutions of phage in water were then spotted on each plate, with 3 µL per spot. Plates were incubated at 37 °C for approximately 20 h and photographed with a Nikon D7500 DSLR camera in a dark room above a white backlight. Images were scaled, rotated, cropped, and converted to grayscale using custom ImageJ Macro Language and Python scripts.

### SeAvs7 affinity purification

Plasmids encoding N-terminal TwinStrep-tagged SeAvs7 or SeAvs7^Δ561–1523^ under an araBAD promoter were transformed into *E. coli* K-12 (ATCC25404) cells. A single colony was inoculated in 20 mL TB with carbenicillin (100 µg/mL) and grown at 37 °C for 16 h. The overnight culture was diluted 1:100 into 2 L fresh TB containing carbenicillin (100 µg/mL) and arabinose (0.2% w/v), and incubated at 37 °C with vigorous shaking for 1.5 h. When OD_600_ reached 0.5−0.8, *Escherichia* phage T1 was added at a multiplicity of infection (MOI) of 2, and cultures were incubated at 37 °C for an additional 30 minutes without shaking. Cells were harvested by centrifugation (4,000 rpm, 10 min, 4 °C).

Cell pellets were resuspended in 5 mL of lysis buffer (50 mM Tris-HCl pH 7.5, 0.5 M NaCl, 5% glycerol, 1 mM ATP, 5 mM MgCl_2_, 1 mM DTT) and lysed by sonication. Insoluble fractions were removed by centrifugation (20,000 xg, 20 min, 4 °C). Clarified lysate mixed with 100 µL of Strep-Tactin SuperFlow Plus resin (Qiagen), and the resin was washed four times with 1 mL of lysis buffer. Bound proteins were eluted three times with 100 µL elution buffer (0.2 M EPPS pH 8.5, 5 mM desthiobiotin). Eluates were analyzed by SDS-PAGE and mass spectrometry (See Proteomics sample preparation, LC-MS analysis, and Proteomics quantification).

### Proteomics sample preparation

Pulldown eluates were digested with Lys-C and trypsin (10 ng/µL each) in 200 mM EPPS pH 8.5 at 37 °C overnight with shaking at 1,200 rpm. Neat acetonitrile was added to each digest to a final concentration of 30% (v/v), and peptides were labeled with 62.5 µg TMTpro-16 reagent^91^ for 1 h at 25 °C with shaking at 900 rpm. Twenty percent of each TMT-labeled sample was pooled, acidified, and diluted with 1% formic acid to pH <3 and a final acetonitrile concentration <5%. The combined sample was desalted with StageTips, dried in a SpeedVac, and reconstituted in 5% acetonitrile / 5% formic acid.

### LC-MS analysis

A total of 1.3 µg of peptides was analyzed by liquid chromatography-mass spectrometry (LC-MS). Peptides were separated on an Aurora Ultimate column 25cm × 75µm with 1.5 µm media and analyzed on an Orbitrap Ascend Tribrid mass spectrometer (ThermoFisher) coupled to a Vanquish Neo UHPLC (ThermoFisher) using a 55-min gradient (5−35.6% acetonitrile, 0.125% formic acid) at a 400 nL/min flow rate. Data-dependent acquisition was used with a mass range of m/z 400−1600. MS1 resolution was set at 120,000, with 50 ms maximum injection time and a 50% normalized automatic gain control (AGC) target. The top 10 most abundant species with charge states 2−6 were subjected to fragmentation. MS2 was performed with standard AGC, 35% normalized collisional energy (NCE), a 120 ms maximum ion injection time, and a 30 s dynamic exclusion window. Fragment ions were further analyzed using multi-notch synchronous precursor selection(SPS)-MS3 (ref.^92^) with 45% NCE to quantify TMT reporter ions.

### Database searching

Database searching was conducted with the Comet algorithm^93^ on Masspike, an in-house search engine described previously^93,94^. Searches used a concatenated target-decoy database comprising proteins from *E. coli* ATCC 25404, *Escherichia* phage T1, the SeAvs7 expression plasmid, common contaminants (*e.g*., keratins, trypsin), and their reversed sequences. Parameters were 25 ppm precursor mass tolerance, 1.0 Da fragment mass tolerance, fully tryptic specificity (K/R) with up to three missed cleavages, and variable Met oxidation (+15.9949 Da). For TMT experiments, TMTpro (+304.2071 Da) on Lys residues and peptide N termini was set as a static modification. Peptide-spectrum matches were filtered by a target–decoy strategy^94–96^ to achieve peptide-level false discovery rate (FDR) <1%, considering XCorr, ΔCn, missed cleavages, peptide length, charge state, and precursor mass accuracy. Short peptides (<7 amino acids) were discarded. Proteins were assembled from identified peptides and controlled to protein-level FDR <1%, and proteins supported by fewer than four unique peptides were discarded.

### Proteomics quantification

For TMT-based quantification, peptide abundance was measured by TMT reporter ions. Reporter intensities were extracted using a 0.003 Th window centered on the theoretical m/z, selecting the most intense peak. Isotopic impurities were corrected based on the manufacturer’s specifications, and TMT signal-to-noise (S/N) values were used for quantification. Peptides were discarded if the summed S/N across 6 channels of each TMT plex was <60 or if isolation specificity was <50%. Protein abundances were computed by summing peptide S/N values. Protein quantification was normalized to ensure equal SeAvs7 protein loading across all TMT channels. SeAvs7 abundance was quantified using peptides mapping to the N-terminal 596 amino acids, including the TwinStrep tag.

### Phage genome sequencing

The genomes of all phages isolated from the City of Palo Alto were sequenced using Tn5 tagmentation and paired-end Illumina sequencing, as previously described^4^. Adapter sequences were trimmed from sequencing reads using CutAdapt with parameters --trim-n -q 20 -m 20 -a CTGTCTCTTATACACATCT -A CTGTCTCT-TATACACATCT. Trimmed reads were assembled into contigs with SPAdes v4.2.0 (ref.^97^) using the --careful option or unicycler v0.5.1 (ref.^98^) with the default option. Genes were annotated by Prokka^99^, Pharokka^100^, Phold^101^, or by comparison with a reference genome of a highly similar phage in GenBank. Isolated phages (Pal001–014) were classified according to the rules of the International Committee on the Taxonomy of Viruses (ICTV)^102,103^. The identities of core structural and replicative genes were confirmed based on HHpred^86^ predictions. Phage phylogenies in Fig. 3j and Extended Data Fig. 5f were generated by concatenating alignments of the large terminases and major capsid proteins. Phage genomes in Extended Data Fig. 5f were visualized using pyGenomeViz.

### Phage gene library construction

To construct each library, barcoded plasmids containing phage genes were pooled along with 8–12 uniquely barcoded mNeonGreen clones as internal normalization controls. The amount of each plasmid in the library was adjusted to account for variations in *E. coli* growth rates due to phage gene expression, in order to ensure balanced plasmid levels across the library after growth in *E. coli*. To determine these adjustments, an initial pilot pool of plasmids, mixed in equal volumes, was transformed into *E. coli* NovaBlue(DE3) (Millipore Sigma) containing an empty pACYC184-like vector. These cells were grown for 12 h at 37°C on minimal salt LB agar (10 g/L tryptone, 5 g/L yeast extract, 0.5 g/L NaCl, 7 g/L agar) containing 100 µg/mL carbenicillin, 25 µg/mL chloramphenicol, and 0.005% arabinose. The relative barcode abundances from plasmids isolated from surviving colonies were quantified by deep sequencing. Based on these abundances, the volume of each plasmid added to the final library was adjusted accordingly.

### Competent cell production

*E. coli* strains were cultured in ZymoBroth with 25 µg/mL chloramphenicol and made competent using a Mix & Go Transformation Kit (Zymo) according to the manufacturer’s recommended protocol.

### Co-expression toxicity screens

Co-expression screens for all Avs systems were performed in *E. coli* NovaBlue(DE3), with the exception of Avs21, which was performed in *E. coli* Stbl3 instead due to its toxicity in *E. coli* NovaBlue(DE3). *E. coli* strains containing Avs plasmids were made competent and transformed with 50 ng of the phage gene library, followed by a 30-minute incubation on ice, a 1 minute heat shock at 42 °C, and a 1 h outgrowth at 37°C in SOC (super optimal broth with catabolite repression). Transformed cells were grown on LB agar plates containing 25 µg/mL chloramphenicol, 100 µg/mL carbenicillin, and 0.01% arabinose. Cells expressing Avs2, Avs12, Avs18, and Avs21 were grown on LB agar Lennox (10 g/L tryptone, 5 g/L yeast extract, 5 g/L NaCl, 7 g/L agar), while cells expressing other Avs families were grown on LB agar with minimal salt (10 g/L tryptone, 5 g/L yeast extract, 0.5 g/L NaCl, 7 g/L agar). For each sample, a corresponding empty vector control was grown in parallel using the same strain and media. Plates were incubated for 12 h at 37 °C, and plasmids from surviving colonies were isolated by miniprep (Qiagen). Barcodes were amplified using Q5 polymerase over 8 cycles of PCR with primers containing Illumina sequencing adaptors and sample-specific 8-nt indexes. Purified amplicons were sequenced on a NextSeq 550 (Illumina) with 20 cycles for Read 1. Barcode counts for each plasmid were determined from demultiplexed FASTQ files, requiring a perfect 20 bp match.

For each sample, barcode counts were normalized by dividing them by the average of the mNeonGreen counts within that sample. For each barcode, synthetic toxicity was then quantified as *q* = (normalized count in Avs sample) / (normalized count in the empty vector (no-Avs) control). For phage genes represented by multiple barcodes, *q* values were averaged across all barcodes for that gene to obtain a single *q* value. Fold depletion was calculated as 1*/q*.

### Anion exchange chromatography

Anion exchange chromatography was performed using an ÄKTA Explorer (Cytiva) equipped with a HiTrap Q HP column (Cytiva). A linear NaCl gradient (0–1 M) in presence of 20 mM Tris-HCl pH 7.5 was applied to purify proteins. Fractions were monitored by UV absorbance at 214nm, 260nm, and 280 nm, and by SDS-PAGE. Desired fractions were concentrated using Amicon Ultra centrifugal filters (Millipore).

### Size-exclusion chromatography

Size-exclusion chromatography (SEC) was performed using an ÄKTA Explorer (Cytiva) equipped with a Superose 6 10/300 GL column (Cytiva). The column was pre-equilibrated with either an ATP-containing SEC buffer (20 mM Tris-HCl pH 7.5, 0.2 M NaCl, 0.1 mM ATP, and 2 mM MgCl_2_) or an ATP-free SEC buffer (20 mM Tris-HCl pH 7.5, 0.2 M NaCl). Fractions were monitored by UV absorbance at 214, 260, 280 nm and SDS-PAGE. Desired fractions were concentrated using Amicon Ultra centrifugal filters (Millipore).

### Protein purification

Plasmids encoding ZL19 MCP-TwinStrep (TS), TS-SUMO-SeAvs7 variants, TS-EcEF-Tu-His6, TS-SUMO-MvAvs13, TS-SUMO-PhiV-1 tail nozzle, TS-SUMO-KpAvs20, or TS-SUMO-HdK1 RecA were transformed into *E. coli* Rosetta (DE3) (Fisher Scientific) or BL21 (DE3) (New England Biolabs). Transformed cells were plated on LB agar and incubated overnight at 37 °C. Single colonies were inoculated into Terrific Broth (TB) with 100 µg/mL carbenicillin and grown overnight at 37 °C. Starter cultures were diluted 1:100 into fresh TB and grown at 37 °C to an OD_600_ of 0.8. Protein expression was induced with 0.05−0.1 mM isopropyl-D-1-thiogalactopyranoside (IPTG), and cultures were incubated at 22 °C for 16 h with ∼220 rpm orbital shaking. Cells were harvested by centrifugation, and pellets were stored at −80 °C.

To purify proteins other than EcEF-Tu, pellets were resuspended in lysis buffer (50 mM Tris-HCl pH 7.5, 500 mM NaCl, 5% glycerol (v/v), 1 mM DTT, 1 mM ATP, 5 mM MgCl_2_, 1x cOmplete Protease Inhibitor (Sigma Aldrich)) and lysed using an LM20 Microfluidizer (Microfluidics) at 28,000 PSI. Lysates were clarified by centrifugation at 32,000 ×g for 30 minutes at 4 °C, and the supernatant was incubated with Strep-Tactin Superflow Plus resin for 1 h at 4 °C. Resin-bound proteins were washed three times with the lysis buffer on a Poly-Prep chromatography column (Bio-Rad). ZL19 MCP was eluted with a buffer containing 20 mM Tris-HCl pH 7.5, 5 mM desthiobiotin. For all other proteins, bdSENP1 protease was added for on-resin cleavage and incubated overnight at 4 °C, and proteins were eluted with 20 mM Tris-HCl pH 7.5. Proteins were further purified by anion exchange chromatography, and buffers were exchanged to 20 mM Tris-HCl pH 7.5, 0.2 M NaCl using Amicon Ultra centrifugal filters (Millipore). For nuclease activity assays, proteins were additionally polished by SEC in ATP-free SEC buffer.

For TS-EcEF-Tu-His_6_, pellets were resuspended in His–lysis buffer (50 mM Tris-HCl pH 7.5, 0.3 M NaCl, 40 mM imidazole, 5% glycerol (v/v), 1x cOmplete protease inhibitor) and lysed by microfluidizer. Clarified lysate was incubated with Ni-NTA Superflow resin (Qiagen), and resin–bound protein was washed three times with His-lysis buffer and eluted with His-elution buffer (50 mM Tris-HCl pH 7.5, 0.1 M NaCl, 500 mM imidazole, 5% glycerol). The eluates were incubated with Strep-Tactin Superflow Plus resin, which was then washed three times with lysis buffer and eluted with 5 mM desthiobiotin. The protein was further purified by anion exchange chromatography. Final polishing was performed by size-exclusion chromatography using a HiLoad 16/600 Superdex 200 pg column (Cytiva) pre-equilibrated with ATP-free SEC buffer. Protein concentration was determined by UV absorbance at 280 nm.

### In vitro DNA cleavage assay

SeAvs7 (0 or 320 nM), ZL19 MCP-TS (1.6 µM or as specified), and TS-EcEF-Tu-His_6_ (10 µM or as specified) were incubated in a reaction buffer containing 20 mM Tris-HCl (pH 7.5), 0.2 M NaCl, 1 mM ATP (or ADP or AMP-PNP as specified), and 5 mM MgCl_2_ for 15 minutes at room temperature. DNA was added at a final concentration of 5 ng/µL, and the reaction mixture was incubated at room temperature for 15 minutes or for the indicated times. DNA products were purified using QIAquick PCR purification columns (Qiagen) and analyzed by gel electrophoresis on an E-Gel EX 2% agarose gel (ThermoFisher). Gel images were acquired using a Bio-Rad VersaDoc imaging system, and the signal intensity for each lane was quantified with the ImageJ Plot Profile tool.

### Complex formation and purification

For SeAvs7 variant–ZL19 MCP complex, SeAvs7 variant (5 µM; 1 eq.) was incubated with ZL19 MCP (10 µM; 2 eq.) and EcEF-Tu (0 or 10 µM; 0 or 2 eq.) in a buffer containing 20 mM Tris-HCl (pH 7.5), 0.2 M NaCl, 1 mM ATP, 5 mM MgCl_2_, and 1 mM DTT for 1 h at room temperature. For the other Avs–phage trigger pairs, Avs protein (10 µM; 1 eq.) was incubated with the cognate phage protein (10 µM; 2 eq.) in the same buffer for 30 minutes at room temperature. Complexes were purified by SEC using an ATP-containing SEC buffer.

### Cryo-EM sample preparation

To assemble the SeAvs7 complex, Strep-Tactin–purified SeAvs7 (5 µM) was incubated with Strep-Tactin–purified ZL19 MCP-TS (10 µM) in 50 mM Tris-HCl (pH 7.5), 0.5 M NaCl, 1 mM ATP, 5 mM MgCl_2_, 1 mM DTT, and 5% glycerol (v/v) for 1 h at room temperature. The resulting complex was purified by SEC using a Superose 6 10/300 Increase GL column equilibrated with SEC buffer containing 20 mM Tris-HCl (pH 7.5), 0.2 M NaCl, 0.1 mM ATP, and 2 mM MgCl_2_. For apo SeAvs7, Strep-Tactin–purified SeAvs7 was directly separated by SEC under the same conditions. The ATP-free SEC buffer was also used for comparison in Supplementary Fig. 6b.

### Cryo-EM sample preparation and data collection

To prepare cryo-EM grids, 3 µL of EcEF-Tu-bound apo SeAvs7 (5 mg/mL) or the SeAvs7–ZL19 MCP–EcEF-Tu complex (3 mg/mL) was applied to freshly glow-discharged 300-mesh Quantifoil R1.2/1.3 holey carbon grids. The grids were vitrified in liquid ethane using a Vitrobot Mark IV (ThermoFisher) at 4 °C and 100% humidity, with blotting conditions set to a wait time of 5 seconds, a blot time of 3 seconds, and a blot force of 3. Cryo-EM data were collected on a 300 kV Titan Krios (S2C2/Stanford or cEMc/Stanford) equipped with a Falcon 4 detector (ThermoFisher) using EPU software in counting mode. For apo SeAvs7, images were acquired at a nominal magnification of 130,000×, corresponding to a physical pixel size of 0.946 Å. For the SeAvs7–ZL19 MCP–EcEF-Tu complex, images were collected at the same magnification, with a pixel size of 0.95 Å. A total of 40 movie stacks per sample were recorded, each with a total electron dose of 50 electrons per Å^2^.

### Cryo-EM data processing

For the SeAvs7–ZL19 MCP–EcEF-Tu complex, 5,805 movies were processed using cryoSPARC live v4.4 (ref.^104^), including Patch motion correction, Patch CTF estimation, template picking, and particle extraction with a box of 120 pixels and 4 × 4 binning. A total of 5,704 images with a CTF resolution better than 8 Å were selected, resulting in 3,769,341 particles. After two rounds of 2D classification, 487,281 particles were selected for ab initio reconstruction to generate initial models. Following two rounds of heterogeneous refinement, 312,068 particles (in two groups containing 205,105 and 106,963 particles) were re-extracted with a box size of 480 pixels. The 205,105 particles in the first group were subjected to one additional round of heterogeneous refinement, after which 78,090 particles were selected for non-uniform refinement, yielding a 3.43 Å resolution map of the tetrameric complex (Map A). To improve the resolution of the four SeAvs7 TPR–ZL19 MCP–EcEF-Tu interfaces, local refinement was performed individually, yielding improved resolutions of 4.01 Å (Map B), 3.78 Å (Map C), 3.77 Å (Map D), and 3.70 Å (Map E), respectively. A composite map (Map F) was generated by combining Maps A–F in Phenix. As SeAvs7 TPR–ZL19 MCP–EcEF-Tu domains were structurally similar, all particles were aligned using the volume alignment tool, resulting in 312,068 particles subjected to further 3D classification, yielding 59,867 particles. These domains were consistent with those of the SeAvs7–ZL19 MCP–EcEF-Tu monomeric complex from the second particle group (with 106,963 particles). Therefore, these particles were combined with the second particle group, resulting in a total of 166,830 particles. These underwent additional 3D classification and local refinement, producing a final 3.40 Å map (Map G). Maps A–E were sharpened by Local resolution filter, and local resolution was estimated using BlocRes.

For the apo SeAvs7 dataset, 1,209 movies were initially motion corrected using MotionCorr2 (ref.^105^). The dose-weighted images were imported into cryoSPARC v4.4 (ref.^104^), where the contrast transfer function (CTF) was estimated using Patch CTF estimation. Template-based particle picking identified 800,233 particles with a box size of 80 pixels and 4×4 binning. After one round of 2D classification, 304,831 particles were selected for ab initio reconstruction, yielding three initial models for subsequent heterogeneous refinement. The entire set of 800,233 particles was subjected to two rounds of heterogeneous refinement, resulting in 207,066 particles. This subset was re-extracted with a box of 320 pixels and further refined using non-uniform refinement, producing a final map at 2.72 Å resolution. Local resolution was estimated using BlocRes, and the map was sharpened using Local resolution filter.

### Model building and refinement

Initial models for all structures were generated using AlphaFold3 predictions and subsequently fitted into the cryo-EM map using Chimera. Iterative manual model building was performed using Coot^106^. The adjusted models were then refined using Phenix^107^ to optimize geometry and stereochemistry. Additionally, refinement was performed using e2gmm_model_refine.py^108^ in EMAN2 (ref.^109^) to achieve improved PDB validation scores. Model quality was assessed by MolProbity. Structures were visualized using PyMOL (Schrödinger) and ChimeraX^110^.

### Structural analyses

Structural models for proteins and complexes were generated with AlphaFold3 or with AlphaFold multimer in ColabFold^111^, requiring a minimum interface predicted template modeling (ipTM) score of 0.7 for Avs complexes. Structures were visualized with PyMOL (Schrödinger) or ChimeraX 1.8 (UCSF). Structure-guided alignments of homologous phage target proteins were obtained with PROMALS3D^50^, using AlphaFold3 models as inputs. Alignable regions, corresponding to the core folds of the target proteins, were used to determine pairwise sequence identity.

To assess the generalizability of Avs7–MCP–EF-Tu complex formation across Avs7 homologs, structures were predicted as a five-component complex (one each of XxAvs7, XxEF-Tu, ZL19 MCP, ATP, and Mg^2+^, where Xx represents distinct bacterial species), with AlphaFold3. Structures were clustered along with the cryo-EM structure of the SeAvs7–ZL19 MCP–EcEF-Tu monomeric complex using Foldseek multimer-cluster^112^ at 60% coverage and a TM-score of 0.70 (Supplementary Table 12). Structures that clustered with the cryo-EM reference are shown in Extended Data Fig. 5d.

## ACKNOWLEDGEMENTS

We thank Dennis Black and Bill Li for assistance with wastewater sample collection, Bharti Singal at Stanford cEMc for assistance with EM data collection, Ross Tomaino for mass spectrometry analysis (Taplin Biological Mass Spectrometry Facility, Harvard Medical School), and Gao lab members for discussions. Some of this work was performed at the Stanford-SLAC Cryo-EM Center (S2C2), which is supported by the National Institutes of Health Common Fund Transformative High-Resolution Cryo-Electron Microscopy program (U24 GM129541). K.S.M., Y.I.W. and E.V.K. were supported by the Intramural Research Program of the National Institutes of Health (NIH). The contributions of the NIH authors are considered Works of the United States Government. The findings and conclusions presented in this paper are those of the author and do not necessarily reflect the views of the NIH or the U.S. Department of Health and Human Services. C.C. is supported by a training grant from NIH Cellular and Molecular Training Grant (NIGMS, grant number 5T32GM007276). S.A.E. is supported by an NIH Genetics and Developmental Biology Training Grant (NIGMS, grant number 5T32GM141828) and by Stanford Bio-X. H.X. is supported by NIH R00AG07346103, the Shmunis Family Innovation Award in Cancer Therapeutics, the Baxter Foundation Faculty Scholars Award, and the V Scholar Award. L.F. is supported by the G. Harold and Leila Y. Mathers Foundation and NIH R35GM153424. A.G. is supported by the G. Harold & Leila Y. Mathers Foundation, the Esther Ehrman Lazard Faculty Scholars Program, and Stanford Bio-X. Phage MRC was a gift from Max Wilkinson.

## AUTHOR CONTRIBUTIONS

H.L. and A.G. conducted bioinformatics analyses of Avs7/Dsr1 and generated AlphaFold models. K.S.M., Y.I.W., E.V.K., H.L., and A.G. performed bioinformatics analyses of STAND NTPases. H.L. performed biochemical experiments, analyzed structures, and generated structural figures. H.L. and A.G. isolated and sequenced phages from wastewater. S.L.-W. and A.G. cloned Avs genes, created phage gene libraries, and performed co-expression toxicity experiments. S.L.-W., C.C., H.L., and A.G. performed phage plaque assays. H.L., S.L.-W., S.A.E., and C.C. purified proteins. J.Z. prepared grids and performed cryo-EM data collection, structure determination, and model building. J.C. and H.X. performed mass spectrometry of SeAvs7 pulldowns. L.F. oversaw cryo-EM studies. All authors analyzed data. A.G. conceived the project and supervised research. H.L., S.L.-W., C.C., and A.G. wrote the manuscript with input from all authors.

## DATA AVAILABILITY

The genome sequences of phages Pal001–013 have been deposited in GenBank under the accession numbers PX985120–PX985132. Cryo-EM maps have been deposited in the Electron Microscopy Data Bank under accession numbers EMD-73001 (Map F; composite map of the SeAvs7–ZL19 MCP–EcEF-Tu tetrameric complex), EMD-73002 (Map B), EMD-73003 (Map A), EMD-73004 (Map E), EMD-73005 (Map D), EMD-73006 (Map C), and EMD-48771 (Apo SeAvs7) and EMD-48772 (Map G; SeAvs7–ZL19 MCP–EcEF-Tu monomeric complex). The corresponding atomic models have been deposited in the Protein Data Bank under accession codes 9YIX (Map F), 9N01 (Map G) and 9N00 (Apo SeAvs7).

---

**Extended Data Fig. 1.**
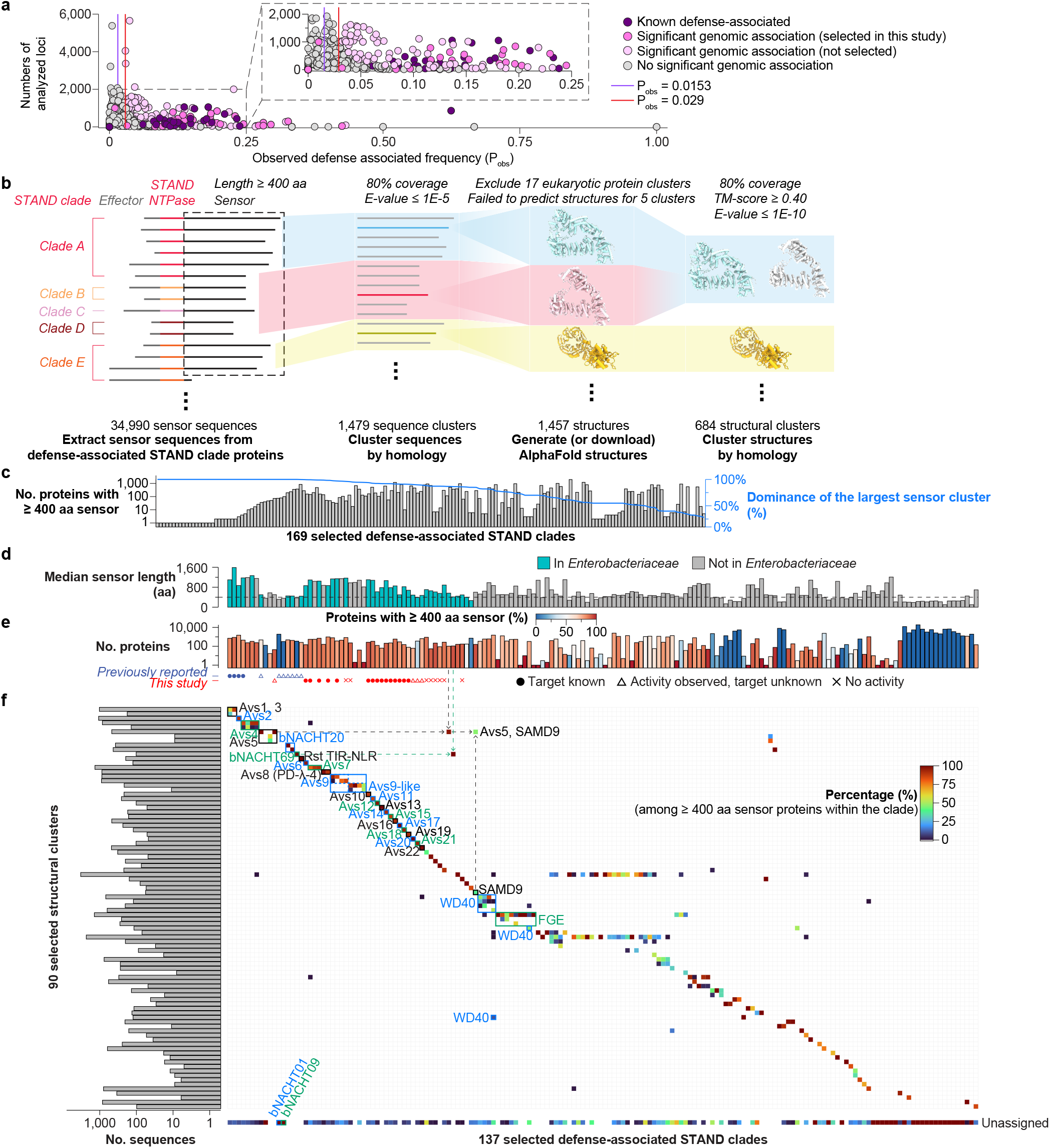
Related to Fig. 1. **a**, Statistics of defense context enrichment for the 589 identified STAND NTPase clades. **b**, Schematic workflow for the sequence- and structure-based clustering of defense-associated sensors. **c**, Number of proteins with ≥400 aa sensors per defense-associated STAND clade, alongside the percent dominance of the largest sensor cluster within each clade. **d**, Median sensor length and **e**, total protein count per selected STAND clade. **f**, Number of proteins for each of the 90 selected structural clusters (left) and their distribution across STAND clades (right). STAND clades lacking sensors ≥400 aa were excluded, and percentages were calculated using sensors ≥400 aa. FGE, formylglycine-generating enzyme.

**Extended Data Fig. 2.**
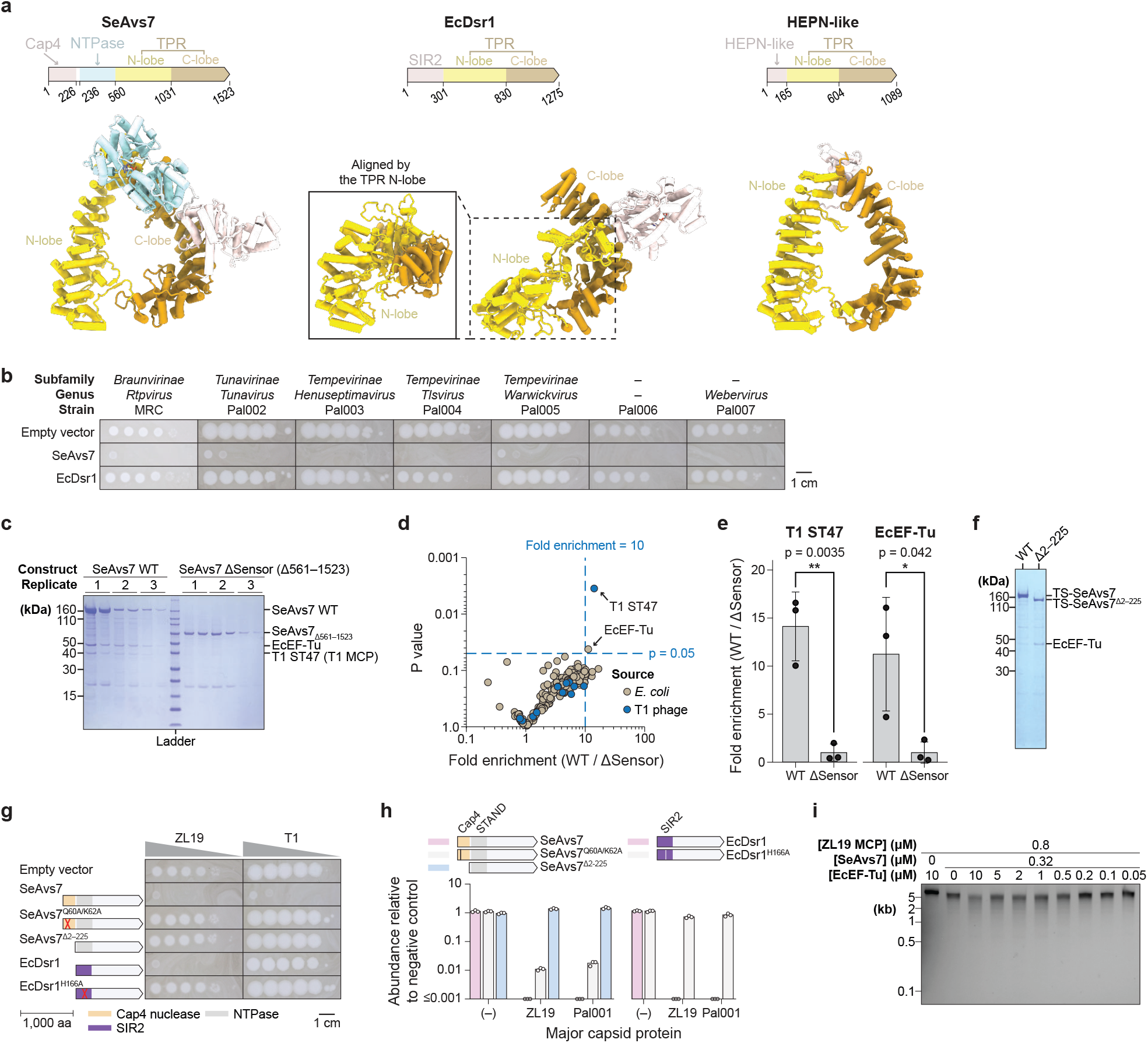
Related to Fig. 2. **a**, AlphaFold3 models of SeAvs7, EcDsr1, and a HEPN-like domain-containing protein from *Streptomyces sp*. NBC_00271, colored by domain. **b**, Plaque assays of *E. coli* expressing an empty vector, SeAvs7, or EcDsr1 against seven *Drexlerviridae* phages. **c**, SDS-PAGE analysis of pulldown samples prepared for mass spectrometry, and **d**, the corresponding volcano plot (axes in logarithmic scale). **e**, Fold enrichment of the T1 major capsid protein (T1 ST47; left) and *E. coli* elongation factor Tu (EcEF-Tu; right) co-purified with wild-type (WT) SeAvs7 compared to the SeAvs7 ΔSensor mutant. Bars represent the mean of the three independent biological replicates, and error bars indicate the standard deviation. Individual data points are shown. **f**, SDS-PAGE analysis of Twin-Strep-tagged WT SeAvs7 and SeAvs7 ΔCap4 (SeAvs7^Δ2–225^). **g**, Plaque assays of *E. coli* expressing the indicated SeAvs7 or EcDsr1 variants, challenged with *Escherichia* phages ZL19 and T1. **h**, Toxicity assays demonstrating that disruption of the SeAvs7 nuclease domain (left) or EcDsr1 NADase domain (right) abrogates cell death upon MCP co-expression. **i**, Agarose gel of *in vitro* nuclease activity assays using a linear dsDNA substrate.

**Extended Data Fig. 3.**
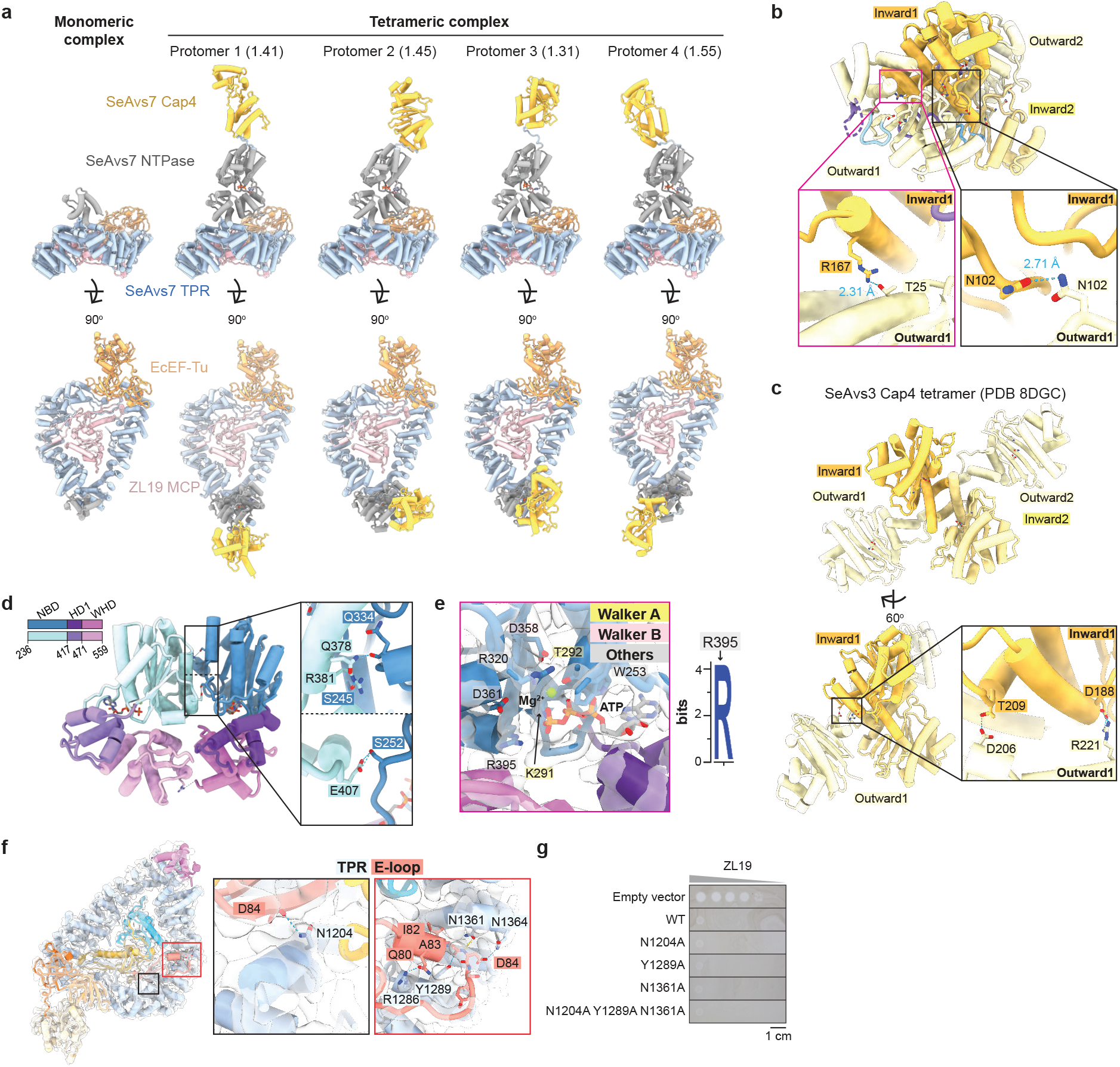
Assembly of the SeAvs7–MCP–EcEF-Tu tetrameric complex. **a**, Structural alignment of the monomeric complex with the four protomers of the tetrameric complex. Root mean square deviation (RMSD) values relative to the monomeric complex are indicated in parentheses. **b, c**, Structures of the Cap4 tetramers of SeAvs7 (**b**) and SeAvs3 (PDB 8DGC) (**c**). Insets highlight interactions between the inward-facing and outward-facing protomers. **d**, Molecular interactions between the SeAvs7 NTPase domains. **e**, ATP and Mg^2+^ bound in the active site of the NTPase domain, with the cryo-EM map displayed as a transparent surface. A sequence logo for SeAvs7 Arg395, generated from 1,757 Avs7 homologs, is also shown. **f**, Molecular interactions between the SeAvs7 TPR domain and ZL19 MCP, overlaid with the cryo-EM map (transparent surface). **g**, Plaque assays of *E. coli* expressing SeAvs7 mutants against phage ZL19.

**Extended Data Fig. 4.**
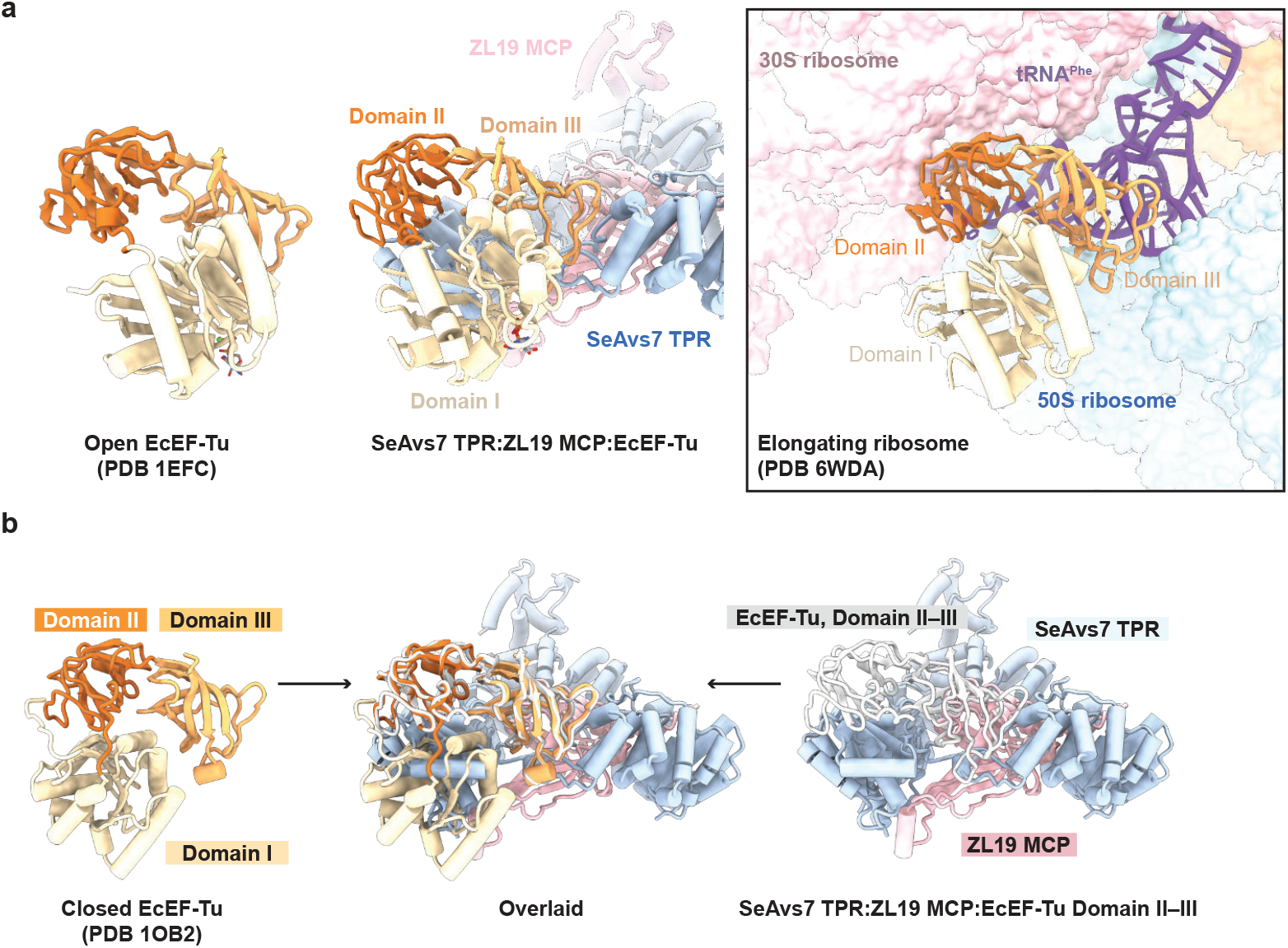
SeAvs7-bound EcEF-Tu adopts an open conformation. **a**, Comparison of the SeAvs7-bound EcEF-Tu structure (left) with a reference open conformation (PDB 1EFC). The inset highlights ribosome-bound EcEF-Tu in the open conformation (PDB 6WDA). **b**, Structures of closed EcEF-Tu (PDB 1OB2; left) and EcEF-Tu Domains II–III bound to SeAvs7 (right), together with a structural superimposition of the two (middle).

**Extended Data Fig. 5.**
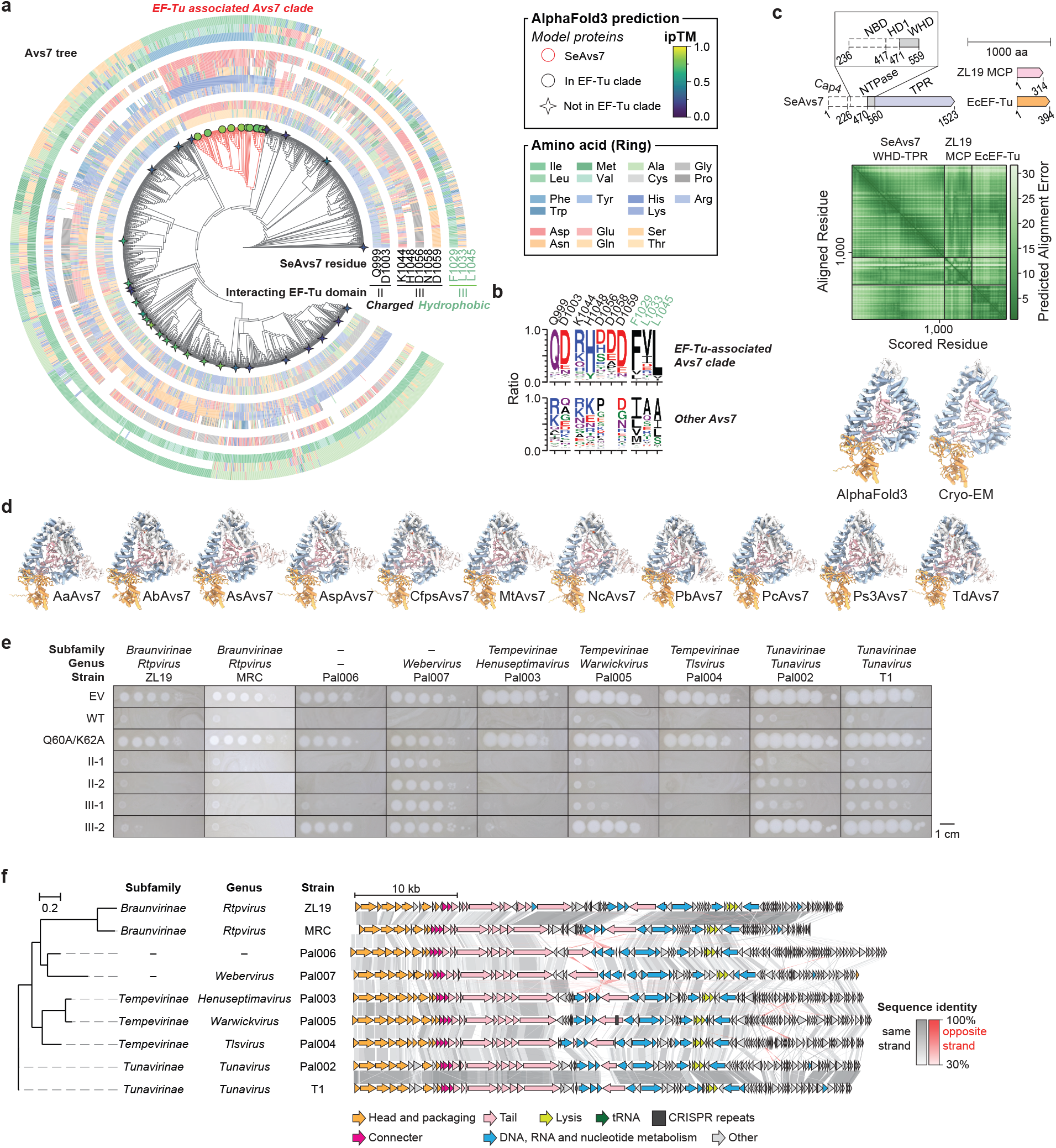
Conservation and functional validation of the Avs7–EF-Tu interaction. **a**, Phylogenetic tree of 1,757 Avs7 homologs. Color strips represent amino acids at the SeAvs7–EcEF-Tu interface. Proteins co-folded with their host EF-Tu and ZL19 MCP are annotated with circles or stars and colored by their ipTM scores. A clade comprising 192 Avs7 homologs likely to form a complex with EF-Tu is highlighted with red branches. **b**, Sequence logos of SeAvs7 residues within the Avs7–EF-Tu interface, comparing the EF-Tu-associated clade (top) with non-associated clades (bottom). **c**, AlphaFold3 model of the SeAvs7 WHD-TPR–ZL19 MCP–EcEF-Tu complex (SeAvs7–EcEF-Tu ipTM 0.85), alongside the predicted alignment error (Å). The cryo-EM structure is shown for comparison. **d**, AlphaFold3 models of 11 Avs7–ZL19 MCP–EF-Tu complexes. Corresponding ipTM scores are provided in Supplementary Table 12. **e**, Plaque assays of *E. coli* expressing SeAvs7 variants against nine *Drexlerviridae* family phages. **f**, Genome architecture and phylogeny of the nine *Drexlerviridae* phages used in (e). Genomes are shown starting with the 5′ end of the small terminase subunit gene. Arrows are color-coded by predicted function and linked by sequence identity.

**Extended Data Fig. 6.**
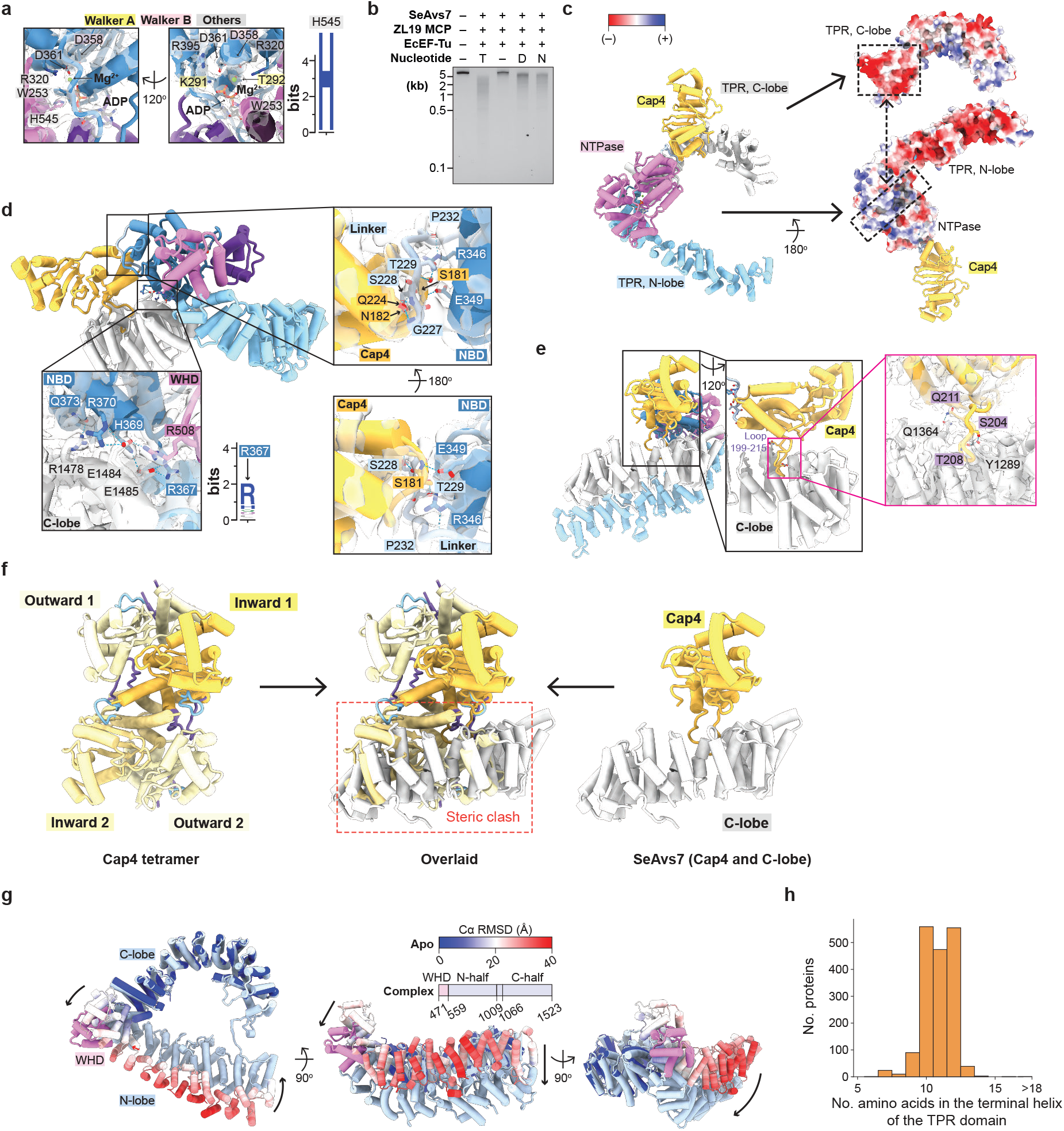
Apo SeAvs7 is inactivated by extensive interactions. **a**, Detailed views of the nucleotide-binding pocket in the apo SeAvs7 NTPase domain, with the cryo-EM map displayed as a transparent surface. A sequence logo for SeAvs7 His545 is shown. **b**, Nuclease activity assay in the presence of distinct nucleotides (T, ATP; D, ADP; N, AMP-PNP). **c**, Electrostatic surfaces of the interfaces between the NTPase domain and the N-lobe and C-lobe of the TPR domain. **d**, Interactions within the NTPase–C-lobe and NTPase–Cap4 interfaces, with a sequence logo for SeAvs7 Arg367. **e**, Molecular interactions between Cap4 and the C-lobe. **f**, Structural overlay of the SeAvs7 Cap4 tetramer and auto-inhibited apo SeAvs7 (Cap4 and C-lobe). **g**, Conformational transition of the SeAvs7 WHD and TPR domains upon binding of ZL19 MCP. Structures are colored by Cα RMSD values (apo SeAvs7 vs. monomeric complex) or by domain (monomeric complex). **h**, Histogram showing the length distribution of the TPR domain terminal helix across 1,757 Avs7 homologs.

**Extended Data Fig. 7.**
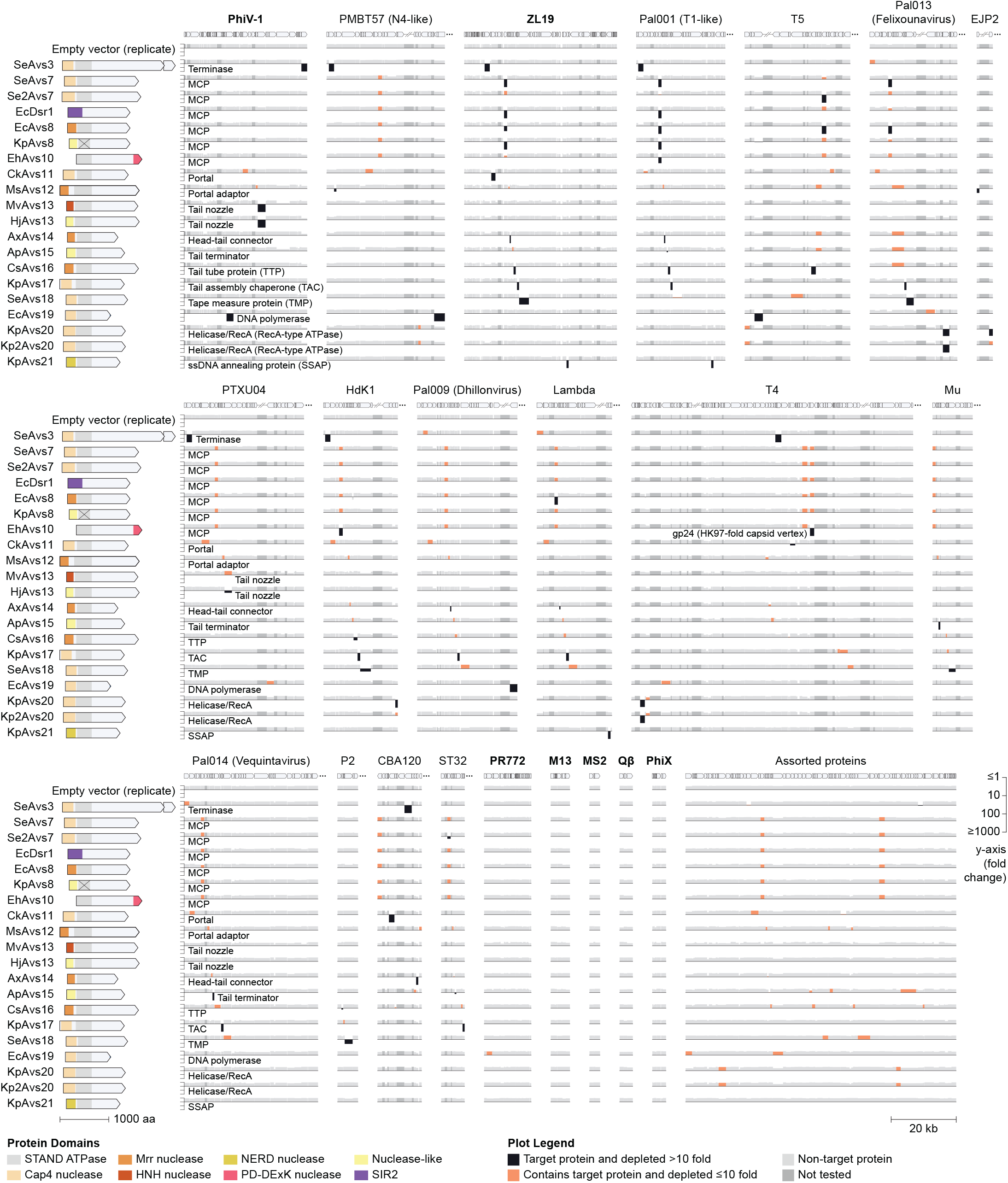
Related to Fig. 5a. Toxicity of Avs genes co-expressed with 600 phage genes in *E. coli*, quantified by barcode abundance following deep sequencing. Bold names indicate phages with complete genomes represented in the library, excluding the dark gray genes due to basal toxicity in *E. coli*. ZL19 data for SeAvs7 and EcDsr1 are replotted from Fig. 2. MCP, major capsid protein.

**Extended Data Fig. 8.**
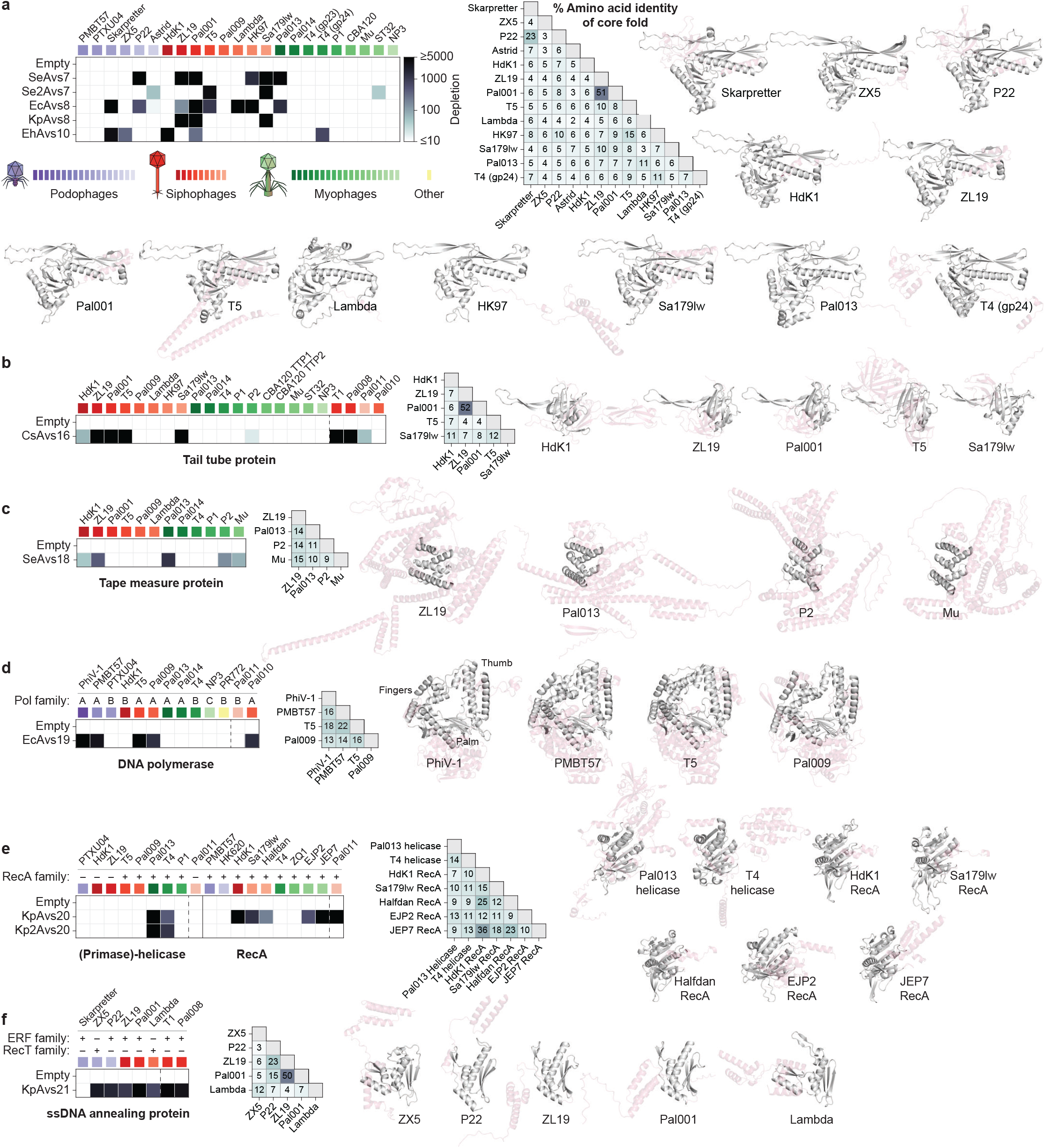
Related to Fig. 5b. Avs pattern recognition of conserved core structural folds. **a–f**, Toxicity quantification and structural comparisons of AlphaFold3 models for major capsid protein (**a**), tail tube protein (**b**), tape measure protein (**c**), DNA polymerase (**d**), primase-helicase and RecA (**e**), and ssDNA annealing protein (**f**). Toxicity data are replotted from Fig. 5b and Supplementary Fig. 14. Pairwise amino acid identities between representative core folds are shown; corresponding AlphaFold3 structures highlight these folds in dark gray. Amino acid identities below ∼5% are considered random. NP3, *Pseudomonas* phage NP3.

**Extended Data Fig. 9.**
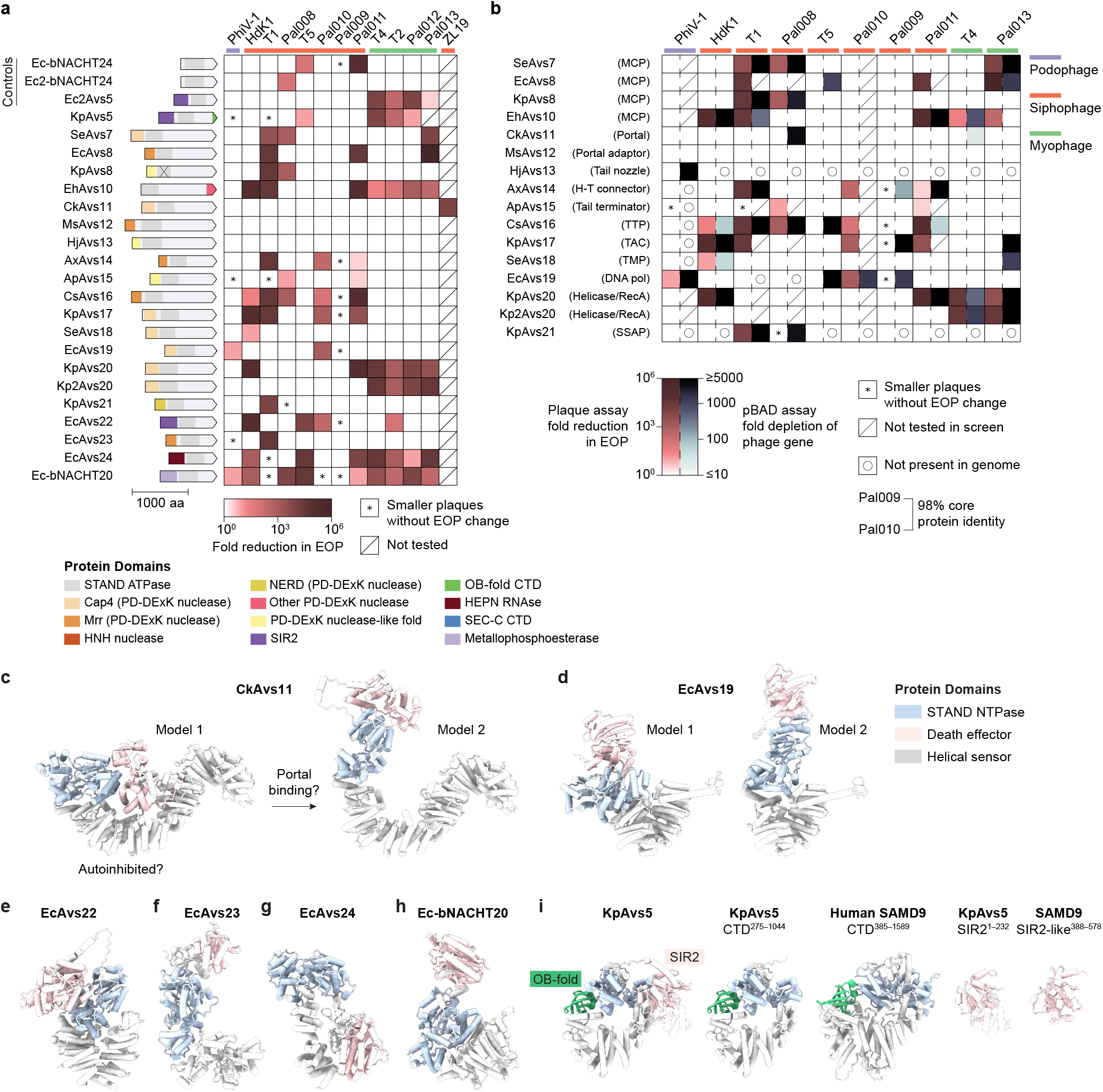
Anti-phage defense activity of Avs proteins. **a**, Efficiency of plating (EOP) of a panel of phages against Avs proteins in *E. coli*. Raw images are shown in Supplementary Fig. 16. **b**, Comparison of phage defense (EOP fold reduction) with co-expression toxicity for each Avs protein and its respective trigger. **c–h**, AlphaFold2/3 models of CkAvs11 (**c**), EcAvs19 (**d**), EcAvs22 (**e**), EcAvs23 (**f**), EcAvs24 (**g**), and Ec-bNACHT20 (**h**). **i**, Structural comparison of KpAvs5 and human SAMD9.

**Extended Data Fig. 10.**
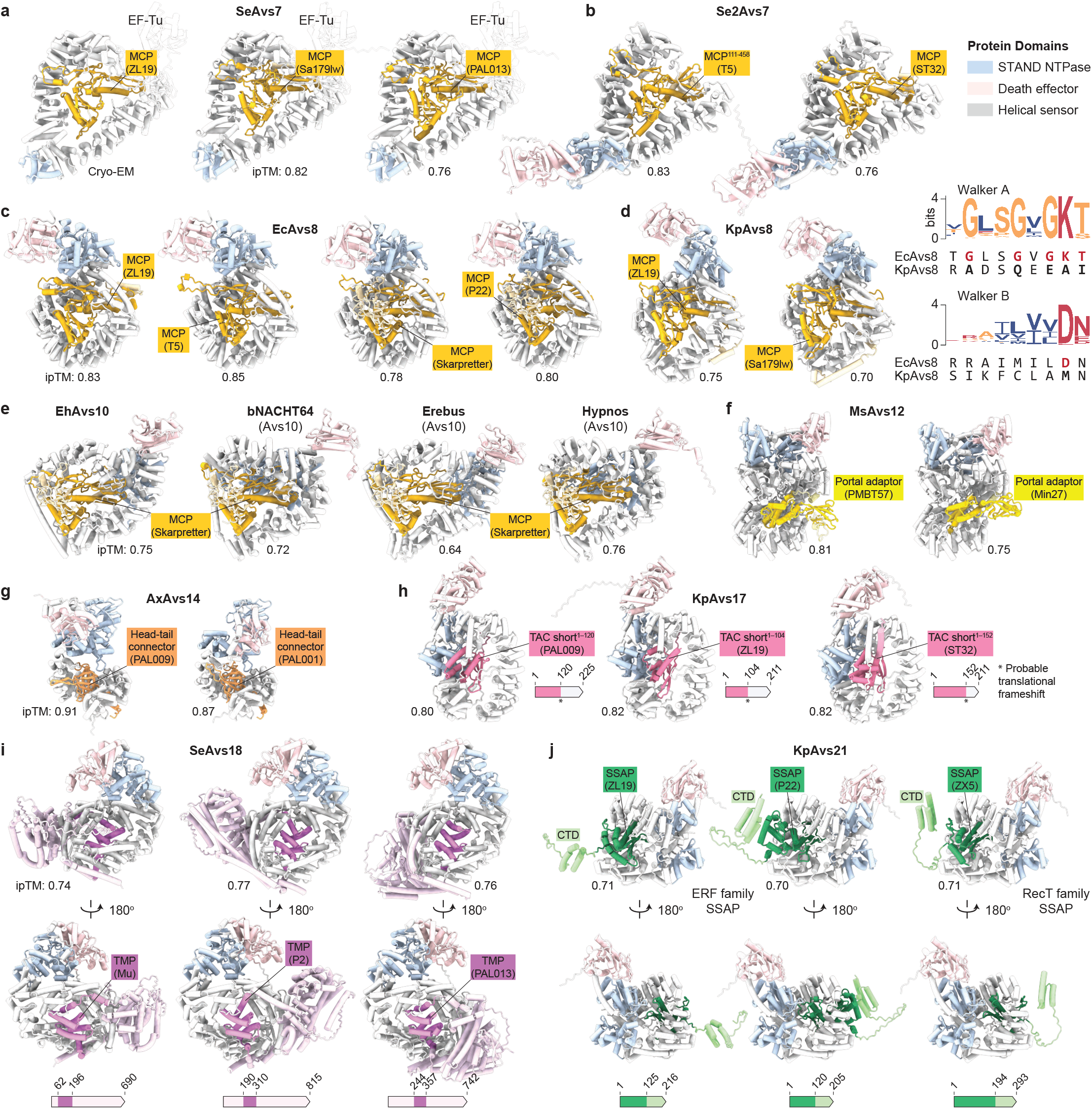
Related to Fig. 6. Additional structural models of Avs pattern recognition of phage proteins. **a–j**, AlphaFold2/3 models of SeAvs7 (**a**), Se2Avs7 (**b**), EcAvs8 (**c**), KpAvs8 (**d**), Avs10 homologs (**e**), MsAvs12 (**f**), AxAvs14 (**g**), KpAvs17 (**h**), SeAvs18 (**i**), and KpAvs21 (**j**) binding their respective target proteins. The ATPase Walker A and Walker B motifs of EcAvs8 and KpAvs8 are shown in (d); the degenerate motifs in KpAvs8 suggest a lack of ATPase activity.

## References

1. Koonin, E. V. & Aravind, L. Origin and evolution of eukaryotic apoptosis: the bacterial connection. Cell Death Differ 9, 394–404 (2002).

2. Leipe, D. D., Koonin, E. V. & Aravind, L. STAND, a class of P-loop NTPases including animal and plant regulators of programmed cell death: multiple, complex domain architectures, unusual phyletic patterns, and evolution by horizontal gene transfer. J Mol Biol 343, 1–28 (2004).

3. Jones, J. D. G., Vance, R. E. & Dangl, J. L. Intracellular innate immune surveillance devices in plants and animals. Science 354, (2016).

4. Gao, L. et al. Diverse enzymatic activities mediate antiviral immunity in prokaryotes. Science 369, 1077–1084 (2020).

5. Gao, L. A. et al. Prokaryotic innate immunity through pattern recognition of conserved viral proteins. Science 377, eabm4096 (2022).

6. Béchon, N. et al. Diversification of molecular pattern recognition in bacterial NLR-like proteins. Nat Commun 15, 9860 (2024).

7. Kibby, E. M. et al. A bacterial NLR-related protein recognizes multiple unrelated phage triggers to sense infection. bioRxiv (2024), doi:10.1101/2024.12.17.629029.

8. Conte, A. N. et al. DnaJ mediates phage sensing by the bacterial NLR-related protein bNACHT25. PLoS Biol 23, e3003203 (2025).

9. Muralidharan, A. et al. Molecular basis for anti-jumbo phage immunity by AVAST type 5. Mol Cell 86, 740–756.e9 (2026).

10. Zhao, Y. et al. The NLRC4 inflammasome receptors for bacterial flagellin and type III secretion apparatus. Nature 477, 596–600 (2011).

11. Kofoed, E. M. & Vance, R. E. Innate immune recognition of bacterial ligands by NAIPs determines inflammasome specificity. Nature 477, 592–595 (2011).

12. Chamaillard, M. et al. An essential role for NOD1 in host recognition of bacterial peptidoglycan containing diaminopimelic acid. Nat Immunol 4, 702–707 (2003).

13. Girardin, S. E. et al. Nod2 is a general sensor of peptidoglycan through muramyl dipeptide (MDP) detection. J Biol Chem 278, 8869–8872 (2003).

14. Vassallo, C. N., Doering, C. R., Littlehale, M. L., Teodoro, G. I. C. & Laub, M. T. A functional selection reveals previously undetected anti-phage defence systems in the E. coli pangenome. Nat Microbiol 7, 1568–1579 (2022).

15. Rousset, F. et al. Phages and their satellites encode hotspots of antiviral systems. Cell Host Microbe 30, 740–753.e5 (2022).

16. Kibby, E. M. et al. Bacterial NLR-related proteins protect against phage. Cell 186, 2410–2424.e18 (2023).

17. van den Berg, D. F. et al. Bacterial homologs of innate eukaryotic antiviral defenses with anti-phage activity highlight shared evolutionary roots of viral defenses. Cell Host Microbe 32, 1427–1443.e8 (2024).

18. Rousset, F. et al. A conserved family of immune effectors cleaves cellular ATP upon viral infection. Cell 186, 3619–3631.e13 (2023).

19. Makarova, K. S., Wolf, Y. I., Snir, S. & Koonin, E. V. Defense islands in bacterial and archaeal genomes and prediction of novel defense systems. J Bacteriol 193, 6039–6056 (2011).

20. Doron, S. et al. Systematic discovery of antiphage defense systems in the microbial pangenome. Science 359, eaar4120 (2018).

21. Shmakov, S. A., Makarova, K. S., Wolf, Y. I., Severinov, K. V. & Koonin, E. V. Systematic prediction of genes functionally linked to CRISPR-Cas systems by gene neighborhood analysis. Proc Natl Acad Sci 115, E5307–E5316 (2018).

22. Brenes, L. R. & Laub, M. T. E. coli prophages encode an arsenal of defense systems to protect against temperate phages. Cell Host Microbe 33, 1004–1018.e5 (2025).

23. Chou, W.-C., Jha, S., Linhoff, M. W. & Ting, J. P.-Y. The NLR gene family: from discovery to present day. Nat Rev Immunol 23, 635–654 (2023).

24. Hu, Z. et al. Structural and biochemical basis for induced self-propagation of NLRC4. Science 350, 399–404 (2015).

25. Zhang, L. et al. Cryo-EM structure of the activated NAIP2-NLRC4 inflammasome reveals nucleated polymerization. Science 350, 404–409 (2015).

26. Zhou, M. et al. Atomic structure of the apoptosome: mechanism of cytochrome c- and dATP-mediated activation of Apaf-1. Genes Dev 29, 2349–2361 (2015).

27. Wang, J. et al. Reconstitution and structure of a plant NLR resistosome conferring immunity. Science 364, (2019).

28. Liu, F. et al. Activation of the helper NRC4 immune receptor forms a hexameric resistosome. Cell 187, 4877–4889.e15 (2024).

29. Wu, Y., Sun, Y., Richet, E., Han, Z. & Chai, J. Structural basis for negative regulation of the Escherichia coli maltose system. Nat Commun 14, 4925 (2023).

30. Poon, K. K., Chu, J. C. & Wong, S. L. Roles of glucitol in the GutR-mediated transcription activation process in Bacillus subtilis: glucitol induces GutR to change its conformation and to bind ATP. J Biol Chem 276, 29819–29825 (2001).

31. Wang, Y. et al. Structural and functional characterization of AfsR, an SARP family transcriptional activator of antibiotic biosynthesis in Streptomyces. PLoS Biol 22, e3002528 (2024).

32. Koonin, E. V. & Aravind, L. The NACHT family - a new group of predicted NTPases implicated in apoptosis and MHC transcription activation. Trends Biochem Sci 25, 223–224 (2000).

33. Legrand, A. et al. Evolutionary characterization of antiviral SAMD9/9L across kingdoms supports ancient convergence and lineage-specific adaptations. Nat Ecol Evol 9, 2206–2222 (2025).

34. Mekhedov, S. L., Makarova, K. S. & Koonin, E. V. The complex domain architecture of SAMD9 family proteins, predicted STAND-like NTPases, suggests new links to inflammation and apoptosis. Biol Direct 12, 13 (2017).

35. Kaur, G., Burroughs, A. M., Iyer, L. M. & Aravind, L. Highly regulated, diversifying NTP-dependent biological conflict systems with implications for the emergence of multicellularity. Elife 9, (2020).

36. Furano, A. V. Content of elongation factor Tu in Escherichia coli. Proc Natl Acad Sci 72, 4780–4784 (1975).

37. Suhanovsky, M. M. & Teschke, C. M. Nature’s favorite building block: Deciphering folding and capsid assembly of proteins with the HK97-fold. Virology 479-480, 487–497 (2015).

38. Duda, R. L. & Teschke, C. M. The amazing HK97 fold: versatile results of modest differences. Curr Opin Virol 36, 9–16 (2019).

39. Koonin, E. V. et al. Global organization and proposed megataxonomy of the virus world. Microbiol Mol Biol Rev 84, (2020).

40. Helgstrand, C. et al. The refined structure of a protein catenane: the HK97 bacteriophage capsid at 3.44 Å resolution. J Mol Biol 334, 885–899 (2003).

41. Hryc, C. F. et al. Accurate model annotation of a near-atomic resolution cryo-EM map. Proc Natl Acad Sci 114, 3103–3108 (2017).

42. Abramson, J. et al. Accurate structure prediction of biomolecular interactions with AlphaFold 3. Nature 630, 493–500 (2024).

43. Loveland, A. B., Demo, G. & Korostelev, A. A. Cryo-EM of elongating ribosome with EF-Tu•GTP elucidates tRNA proofreading. Nature 584, 640–645 (2020).

44. Song, H., Parsons, M. R., Rowsell, S., Leonard, G. & Phillips, S. E. Crystal structure of intact elongation factor EF-Tu from Escherichia coli in GDP conformation at 2.05 Å resolution. J Mol Biol 285, 1245–1256 (1999).

45. Veesler, D. & Cambillau, C. A common evolutionary origin for tailed-bacteriophage functional modules and bacterial machineries. Microbiol Mol Biol Rev 75, 423–33, first page of table of contents (2011).

46. Nagy, T. A., Gersabeck, G. W., Conte, A. N. & Whiteley, A. T. A phage protein screen identifies triggers of the bacterial innate immune system. Nat Microbiol 11, 597–609 (2026).

47. Iyer, L. M., Koonin, E. V. & Aravind, L. Classification and evolutionary history of the single-strand annealing proteins, RecT, Redβ, ERF and RAD52. BMC Genomics 3, 8 (2002).

48. Zinke, M., Schröder, G. F. & Lange, A. Major tail proteins of bacteriophages of the order Caudovirales. J Biol Chem 298, 101472 (2022).

49. Xu, J., Hendrix, R. W. & Duda, R. L. Chaperone-protein interactions that mediate assembly of the bacteriophage lambda tail to the correct length. J Mol Biol 426, 1004–1018 (2014).

50. Pei, J.Kim, B.-H. & Grishin, N. V. PROMALS3D: a tool for multiple protein sequence and structure alignments. Nucleic Acids Res 36, 2295–2300 (2008).

51. Leipe, D. D., Aravind, L., Grishin, N. V. & Koonin, E. V. The bacterial replicative helicase DnaB evolved from a RecA duplication. Genome Res 10, 5–16 (2000).

52. Kao, C., Gumbs, E. & Snyder, L. Cloning and characterization of the Escherichia coli lit gene, which blocks bacteriophage T4 late gene expression. J Bacteriol 169, 1232–1238 (1987).

53. Bergsland, K. J., Kao, C., Yu, Y. T., Gulati, R. & Snyder, L. A site in the T4 bacteriophage major head protein gene that can promote the inhibition of all translation in Escherichia coli. J Mol Biol 213, 477–494 (1990).

54. Zhang, T. et al. Direct activation of a bacterial innate immune system by a viral capsid protein. Nature 612, 132–140 (2022).

55. Zhang, T. et al. A bacterial immunity protein directly senses two disparate phage proteins. Nature 635, 728–735 (2024).

56. Bouchard, J. D. & Moineau, S. Lactococcal phage genes involved in sensitivity to AbiK and their relation to single-strand annealing proteins. J Bacteriol 186, 3649–3652 (2004).

57. Yasui, R., Washizaki, A., Furihata, Y., Yonesaki, T. & Otsuka, Y. AbpA and AbpB provide anti-phage activity in Escherichia coli. Genes Genet Syst 89, 51–60 (2014).

58. Depardieu, F. et al. A eukaryotic-like serine/threonine kinase protects Staphylococci against phages. Cell Host Microbe 20, 471–481 (2016).

59. Stokar-Avihail, A. et al. Discovery of phage determinants that confer sensitivity to bacterial immune systems. Cell 186, 1863–1876.e16 (2023).

60. Tuck, O. T. et al. Genome integrity sensing by the broad-spectrum Hachiman antiphage defense complex. Cell 187, 6914–6928.e20 (2024).

61. Yu, Y. T. & Snyder, L. Translation elongation factor Tu cleaved by a phage-exclusion system. Proc Natl Acad Sci 91, 802–806 (1994).

62. Georgiou, T. et al. Specific peptide-activated proteolytic cleavage of Escherichia coli elongation factor Tu. Proc Natl Acad Sci 95, 2891–2895 (1998).

63. Bingham, R., Ekunwe, S. I., Falk, S., Snyder, L. & Kleanthous, C. The major head protein of bacteriophage T4 binds specifically to elongation factor Tu. J Biol Chem 275, 23219–23226 (2000).

64. Rousset, F. et al. TIR signaling activates caspase-like immunity in bacteria. Science 387, 510–516 (2025).

65. Schmitz, M., Querques, I., Oberli, S., Chanez, C. & Jinek, M. Structural basis for the assembly of the type V CRISPR-associated transposon complex. Cell 185, 4999–5010.e17 (2022).

66. Kidmose, R. T., Vasiliev, N. N., Chetverin, A. B., Andersen, G. R. & Knudsen, C. R. Structure of the Qβ replicase an RNA-dependent RNA polymerase consisting of viral and host proteins. Proc Natl Acad Sci 107, 10884–10889 (2010).

67. Takeshita, D. & Tomita, K. Assembly of Qβ viral RNA polymerase with host translational elongation factors EF-Tu and -Ts. Proc Natl Acad Sci 107, 15733–15738 (2010).

68. Harvey, K. L., Jarocki, V. M., Charles, I. G. & Djordjevic, S. P. The diverse functional roles of elongation factor Tu (EF-Tu) in microbial pathogenesis. Front Microbiol 10, 2351 (2019).

69. Sharif, H. et al. Structural mechanism for NEK7-licensed activation of NLRP3 inflammasome. Nature 570, 338–343 (2019).

70. Schäffer, A. A. et al. Improving the accuracy of PSI-BLAST protein database searches with composition-based statistics and other refinements. Nucleic Acids Res 29, 2994–3005 (2001).

71. Steinegger, M. & Söding, J. MMseqs2 enables sensitive protein sequence searching for the analysis of massive data sets. Nat Biotechnol 35, 1026–1028 (2017).

72. Edgar, R. C. Muscle5: High-accuracy alignment ensembles enable unbiased assessments of sequence homology and phylogeny. Nat Commun 13, 6968 (2022).

73. Esterman, E. S., Wolf, Y. I., Kogay, R., Koonin, E. V. & Zhaxybayeva, O. Evolution of DNA packaging in gene transfer agents. Virus Evol 7, veab015 (2021).

74. Deorowicz, S., Debudaj-Grabysz, A. & GudysÂ, A. FAMSA: Fast and accurate multiple sequence alignment of huge protein families. Sci Rep 6, 33964 (2016).

75. Lefort, V., Desper, R. & Gascuel, O. FastME 2.0: A comprehensive, accurate, and fast distance-based phylogeny inference program. Mol Biol Evol 32, 2798–2800 (2015).

76. Marchler-Bauer, A. et al. CDD/SPARCLE: functional classification of proteins via subfamily domain architectures. Nucleic Acids Res 45, D200–D203 (2017).

77. Yu, G., Smith, D. K., Zhu, H., Guan, Y. & Lam, T. T.-Y. GGTREE: An R package for visualization and annotation of phylogenetic trees with their covariates and other associated data. Methods Ecol Evol 8, 28–36 (2017).

78. Tesson, F. et al. Systematic and quantitative view of the antiviral arsenal of prokaryotes. Nat Commun 13, 2561 (2022).

79. Jumper, J. et al. Highly accurate protein structure prediction with AlphaFold. Nature 596, 583–589 (2021).

80. Varadi, M. et al. AlphaFold Protein Structure Database in 2024: providing structure coverage for over 214 million protein sequences. Nucleic Acids Res 52, D368–D375 (2024).

81. van Kempen, M. et al. Fast and accurate protein structure search with Foldseek. Nat Biotechnol 42, 243–246 (2023).

82. Wishart, D. S. et al. PHASTEST: faster than PHASTER, better than PHAST. Nucleic Acids Res 51, W443–W450 (2023).

83. Katoh, K., Misawa, K.Kuma, K.-I. & Miyata, T. MAFFT: a novel method for rapid multiple sequence alignment based on fast Fourier transform. Nucleic Acids Res 30, 3059–3066 (2002).

84. Capella-Gutiérrez, S., Silla-Martínez, J. M. & Gabaldón, T. trimAl: a tool for automated alignment trimming in large-scale phylogenetic analyses. Bioinformatics 25, 1972–1973 (2009).

85. Nguyen, L.-T., Schmidt, H. A., von Haeseler, A. & Minh, B. Q. IQ-TREE: a fast and effective stochastic algorithm for estimating maximum-likelihood phylogenies. Mol Biol Evol 32, 268–274 (2015).

86. Zimmermann, L. et al. A completely reimplemented MPI Bioinformatics Toolkit with a new HHpred Server at its core. J Mol Biol 430, 2237–2243 (2018).

87. Letunic, I. & Bork, P. Interactive Tree Of Life (iTOL) v5: an online tool for phylogenetic tree display and annotation. Nucleic Acids Res 49, W293–W296 (2021).

88. Minh, B. Q. et al. IQ-TREE 2: New models and efficient methods for phylogenetic inference in the genomic era. Mol Biol Evol 37, 1530–1534 (2020).

89. Picelli, S. et al. Tn5 transposase and tagmentation procedures for massively scaled sequencing projects. Genome Res 24, 2033–2040 (2014).

90. Baba, T. et al. Construction of Escherichia coli K-12 inframe, single-gene knockout mutants: the Keio collection. Mol Syst Biol 2, 2006.0008 (2006).

91. Li, J. et al. TMTpro reagents: a set of isobaric labeling mass tags enables simultaneous proteome-wide measurements across 16 samples. Nat Methods 17, 399–404 (2020).

92. McAlister, G. C. et al. MultiNotch MS3 enables accurate, sensitive, and multiplexed detection of differential expression across cancer cell line proteomes. Anal Chem 86, 7150–7158 (2014).

93. Eng, J. K., Jahan, T. A. & Hoopmann, M. R. Comet: an open-source MS/MS sequence database search tool. Proteomics 13, 22–24 (2013).

94. Huttlin, E. L. et al. A tissue-specific atlas of mouse protein phosphorylation and expression. Cell 143, 1174–1189 (2010).

95. Elias, J. E. & Gygi, S. P. Target-decoy search strategy for increased confidence in large-scale protein identifications by mass spectrometry. Nat Methods 4, 207–214 (2007).

96. Peng, J., Elias, J. E., Thoreen, C. C., Licklider, L. J. & Gygi, S. P. Evaluation of multidimensional chromatography coupled with tandem mass spectrometry (LC/LC-MS/MS) for large-scale protein analysis: the yeast proteome. J Proteome Res 2, 43–50 (2003).

97. Bankevich, A. et al. SPAdes: a new genome assembly algorithm and its applications to single-cell sequencing. J Comput Biol 19, 455–477 (2012).

98. Wick, R. R., Judd, L. M., Gorrie, C. L. & Holt, K. E. Unicycler: Resolving bacterial genome assemblies from short and long sequencing reads. PLoS Comput Biol 13, e1005595 (2017).

99. Seemann, T. Prokka: rapid prokaryotic genome annotation. Bioinformatics 30, 2068–2069 (2014).

100. Bouras, G. et al. Pharokka: a fast scalable bacteriophage annotation tool. Bioinformatics 39, btac776 (2023).

101. Bouras, G. et al. Protein structure informed bacteriophage genome annotation with phold. bioRxiv (2025), doi:10.1101/2025.08.05.668817.

102. Grigson, S. R., Giles, S. K., Edwards, R. A. & Papudeshi, B. Knowing and naming: phage annotation and nomenclature for phage therapy. Clin Infect Dis 77, S352–S359 (2023).

103. Adriaenssens, E. & Brister, J. R. How to name and classify your phage: an informal guide. Viruses 9, (2017).

104. Punjani, A., Rubinstein, J. L., Fleet, D. J. & Brubaker, M. A. cryoSPARC: algorithms for rapid unsupervised cryo-EM structure determination. Nat Methods 14, 290–296 (2017).

105. Zheng, S. Q. et al. MotionCor2: anisotropic correction of beam-induced motion for improved cryo-electron microscopy. Nat Methods 14, 331–332 (2017).

106. Emsley, P. & Cowtan, K. Coot: model-building tools for molecular graphics. Acta Crystallogr D Biol Crystallogr 60, 2126–2132 (2004).

107. Adams, P. D. et al. PHENIX: a comprehensive Python-based system for macromolecular structure solution. Acta Crystallogr D Biol Crystallogr 66, 213–221 (2010).

108. Chen, M. Building molecular model series from heterogeneous CryoEM structures using Gaussian mixture models and deep neural networks. Commun Biol 8, 798 (2025).

109. Tang, G. et al. EMAN2: an extensible image processing suite for electron microscopy. J Struct Biol 157, 38–46 (2007).

110. Pettersen, E. F. et al. UCSF ChimeraX: Structure visualization for researchers, educators, and developers. Protein Sci 30, 70–82 (2021).

111. Mirdita, M. et al. ColabFold: making protein folding accessible to all. Nat Methods 19, 679–682 (2022).

112. Kim, W. et al. Rapid and sensitive protein complex alignment with Foldseek-Multimer. Nat Methods 22, 469–472 (2025).

