## Supplementary Figures 1-18 for "Diverse bacterial pattern recognition receptors sense the core phage proteome"

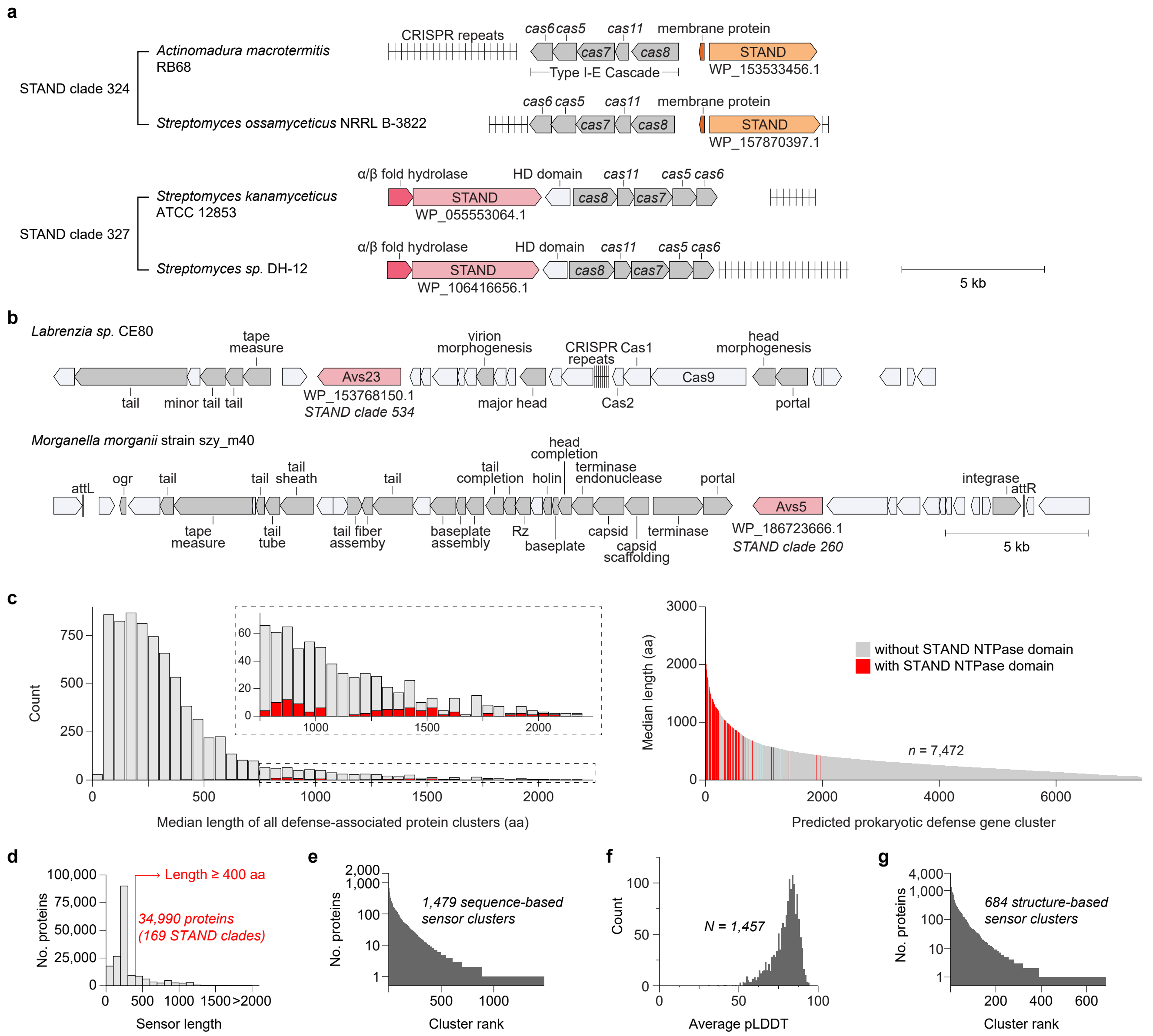


**Supplementary Fig. 1 | Related to Fig. 1. a**, Representative genomic loci of CRISPR-associated STANDs of unknown function in Actinomycetota. **b**, Representative genomic loci of prophage-encoded Avs5 and Avs23. **c**, Median lengths of 7,472 defense-associated gene clusters predicted previously (Gao et al. *Science* **369**, 1077–1084 (2020)), with STAND NTPases highlighted in red. **d**, Histogram of the sensor domain lengths for proteins within defense-associated STAND clades ($n$ = 178,614). **e**, Number of proteins within each of the 1,479 sequence-based sensor clusters. **f**, Distribution of average pLDDT scores for the AlphaFold models of 1,457 sensors. **g**, Number of proteins within each of the 684 structure-based sensor clusters.

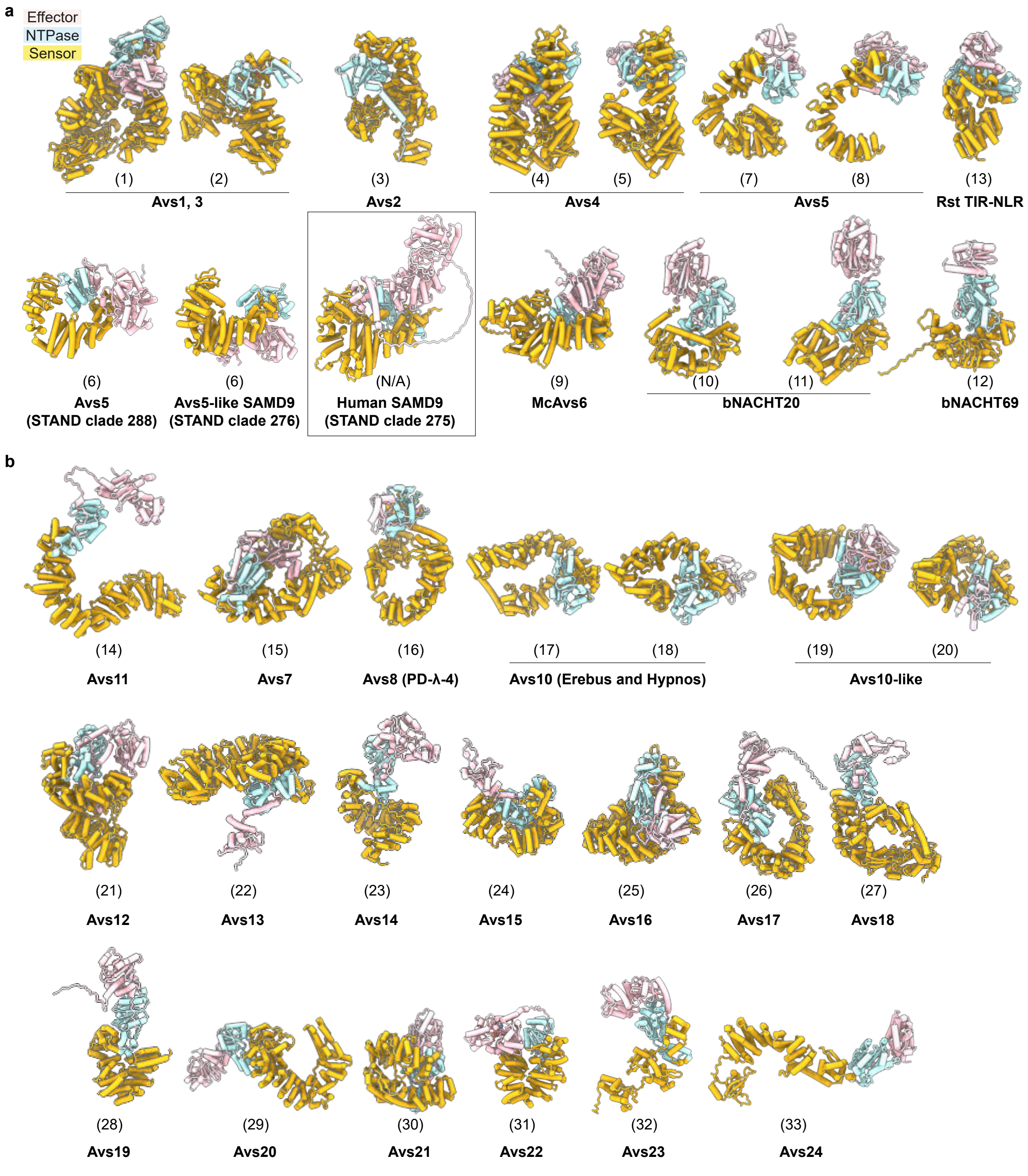


**Supplementary Fig. 2 | Related to Fig. 1. a, b**, AlphaFold3 models of representatives of structural clusters that were previously characterized (**a**) or investigated in the current study (**b**). Sensor structural cluster numbers are shown in parentheses. Human SAMD9 is shown in the inset for comparison.

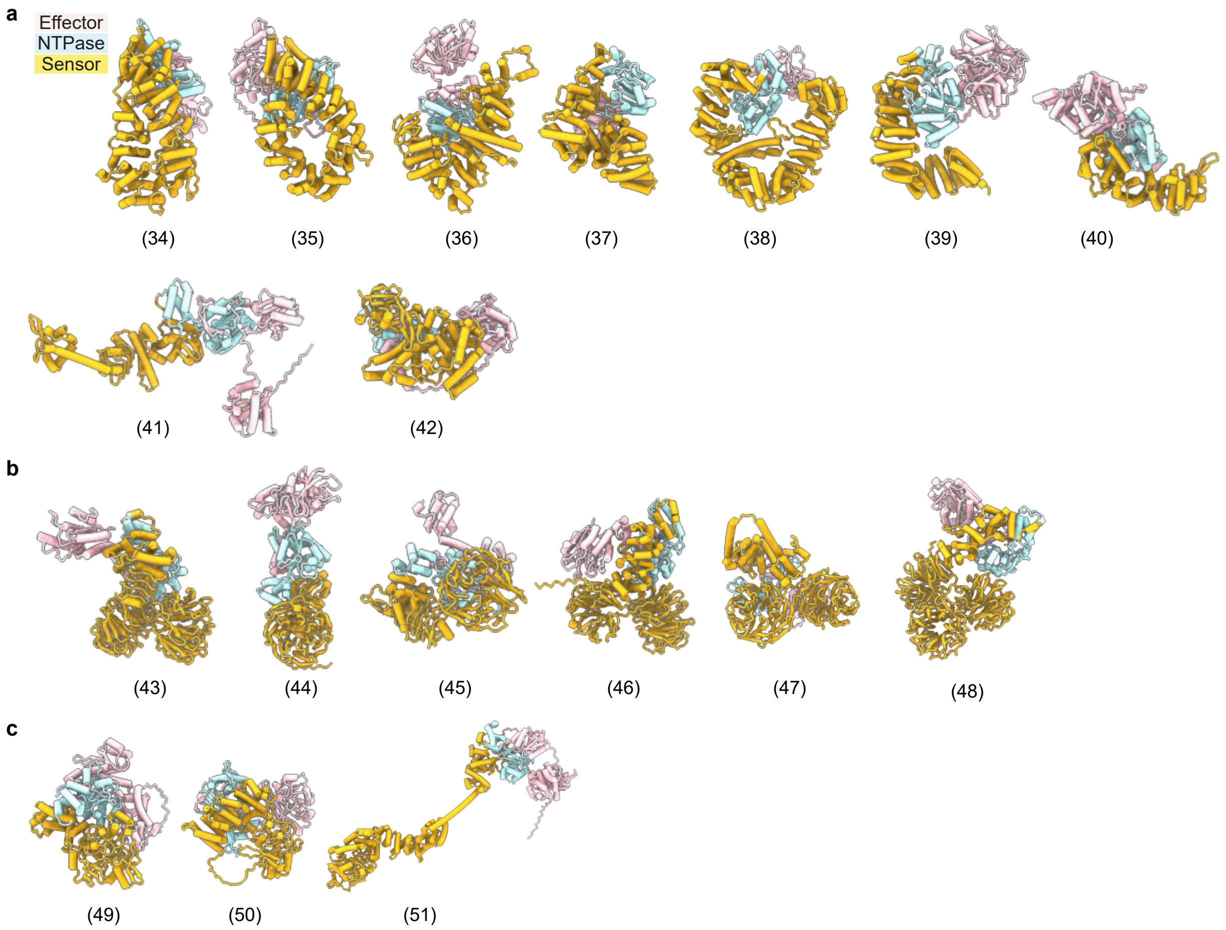


**Supplementary Fig. 3 | Related to Fig. 1. a**, AlphaFold3 models of representative sensors in *Enterobacteriaceae*. **b, c**, AlphaFold3 models of representative sensors containing a WD40 domain (**b**) or a formylglycine-generating enzyme (FGE) domain (**c**). Sensor structural cluster numbers are shown in parentheses.

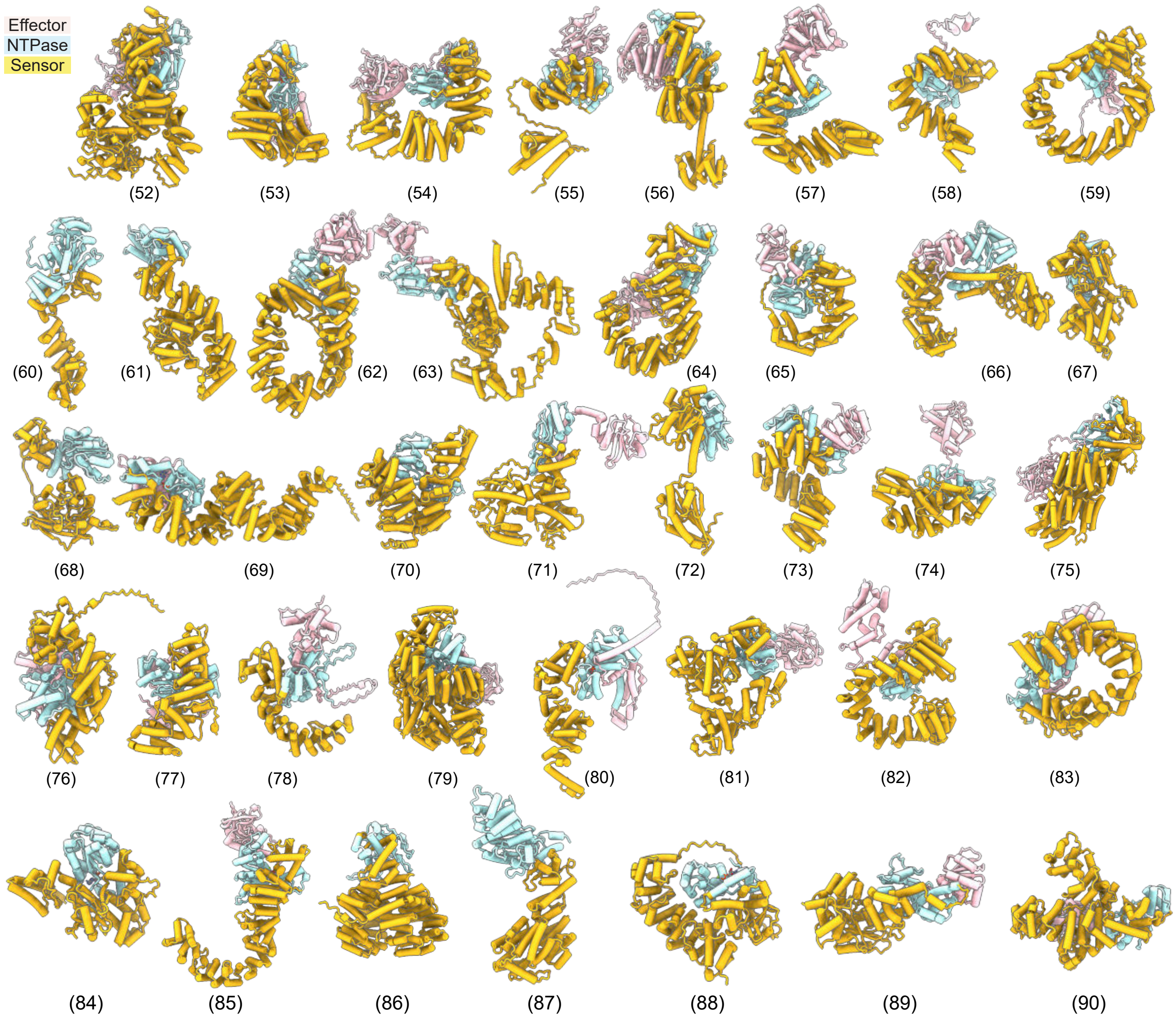


**Supplementary Fig. 4 | Related to Fig. 1.** AlphaFold3 models of 39 additional representative sensors. Sensor structural cluster numbers are shown in parentheses.

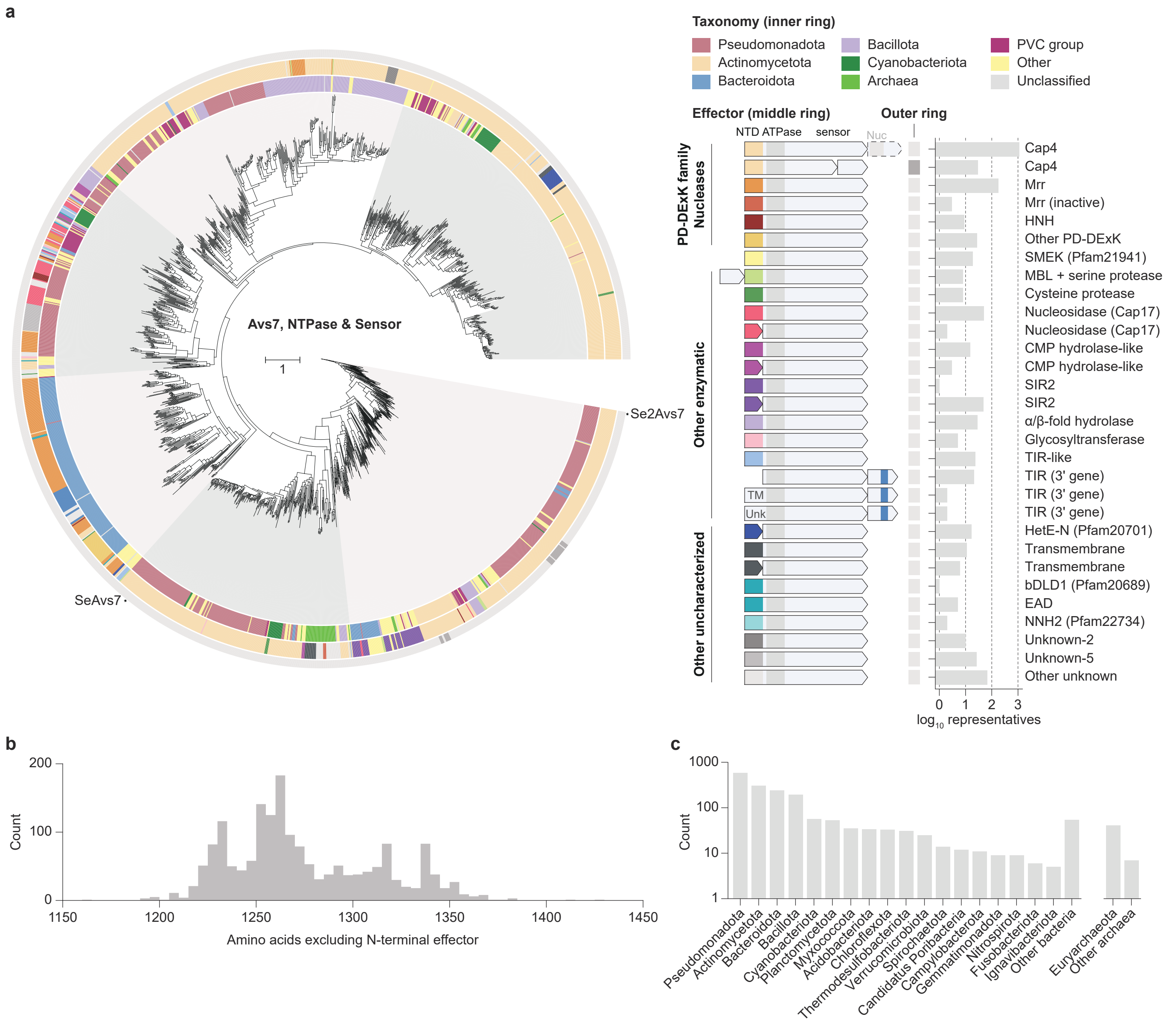


**Supplementary Fig. 5 | Prevalence and diversity of Avs7 homologs. a**, Maximum likelihood tree based on the NTPase and C-terminal domains of 1,757 representative Avs7 homologs. The inner ring indicates taxonomy, and the middle and outer rings represent the effector and sensor domains, respectively. MBL, metallo-β-lactamase; EAD, effector-associated domain; TM, transmembrane domain; Unk, unknown. **b**, Histogram of the combined length of the NTPase and C-terminal domains across Avs7 homologs. **c**, Abundance of Avs7 homologs across bacterial and archaeal phyla.

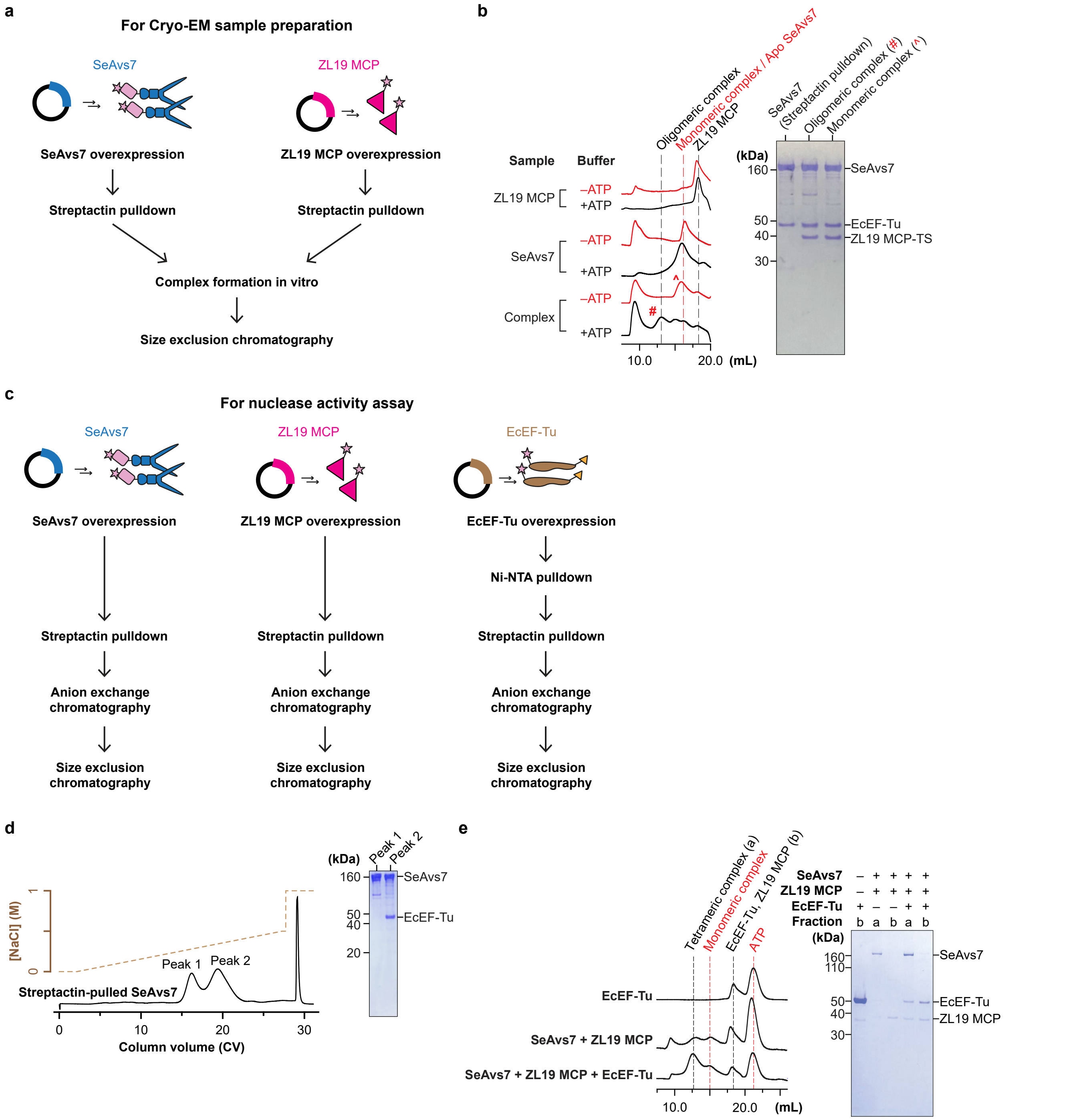


**Supplementary Fig. 6 | SeAvs7 binds ZL19 MCP and EcEF-Tu independently of ATP.** **a**, Schematic of the SeAvs7–ZL19 MCP complex purification workflow for structural analysis. **b**, Size-exclusion chromatograms of Strep-Tactin-purified SeAvs7, ZL19 MCP, and their *in vitro* assembled complex, with SDS-PAGE analysis of selected fractions. Dashed lines indicate expected retention volumes. **c**, Schematic of the SeAvs7, ZL19 MCP, and EcEF-Tu purification workflow for *in vitro* nuclease assays. **d**, Anion-exchange chromatograms of purified SeAvs7 and SDS-PAGE analysis of two peaks. **e**, Size-exclusion chromatograms and SDS-PAGE analysis of complex formation with purified proteins used for nuclease assays.

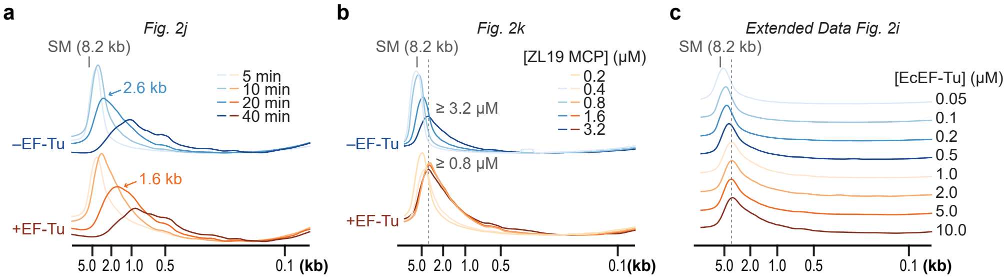


**Supplementary Fig. 7 | Quantification of in vitro cleavage kinetics. a–c**, Densitometric quantification of substrate DNA degradation from raw agarose gels. The most frequent fragment lengths (**a**) or the maximum fragment shift (**b, c**) observed are indicated. SM, substrate double-stranded DNA (8.2 kb).

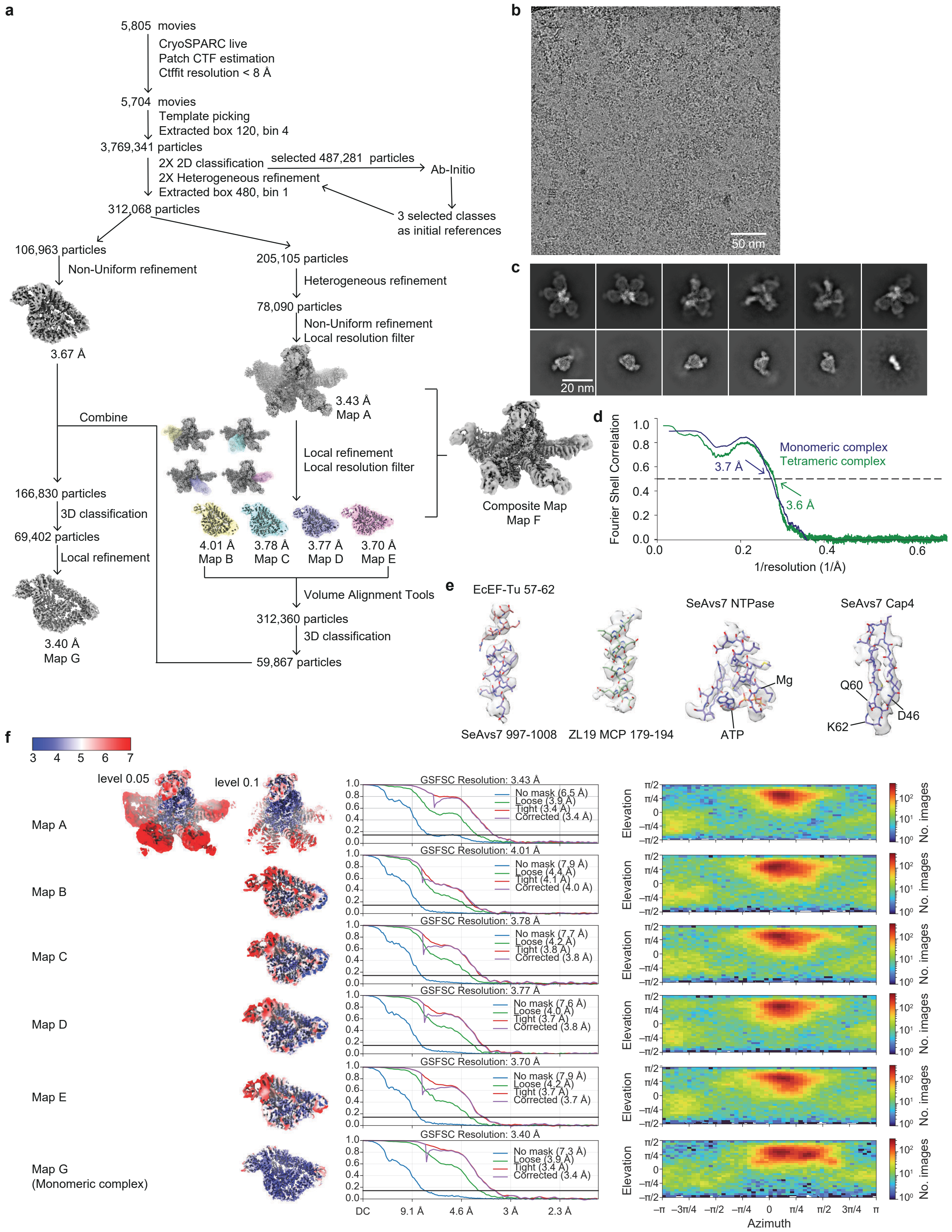


**Supplementary Fig. 8 | Cryo-EM data processing and validation for the SeAvs7–ZL19 MCP–EcEF-Tu complex. a**, Data processing workflow. **b**, Representative cryo-EM micrograph. **c**, Representative 2D class averages. **d**, Map-to-model Fourier shell correlation (FSC) curve. **e**, Representative cryo-EM density from the composite map overlaid with the atomic model. **f**, Quality assessment metrics, including local resolution maps (left), gold-standard FSC curves (middle), and angular particle distributions (right).

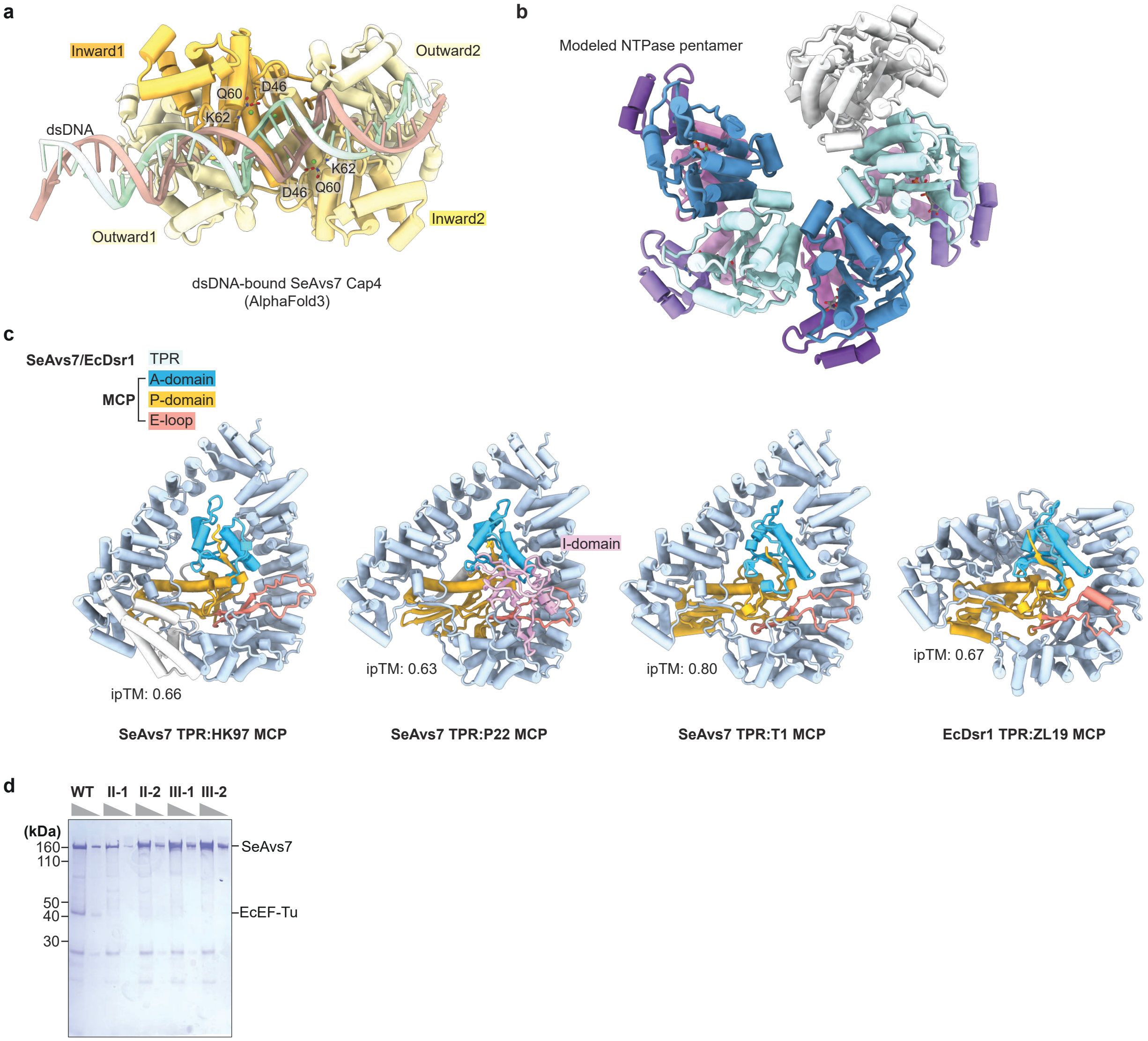


**Supplementary Fig. 9 | Related to Fig. 3. a**, AlphaFold3 model of the SeAvs7 Cap4 tetramer bound to a double-stranded DNA substrate and four $Mg^{2+}$ ions. Interface predicted template modeling (ipTM) scores between the DNA strands and Cap4 monomers range from 0.54 to 0.60. **b**, A model of an SeAvs7 NTPase domain pentamer generated from a structural alignment to the NTPase tetramer. **c**, AlphaFold3 models of the SeAvs7 and EcDsr1 TPR–MCP complexes, with ipTM scores indicated. **d**, SDS-PAGE analysis of wild-type and mutant SeAvs7 purified from *E. coli*.

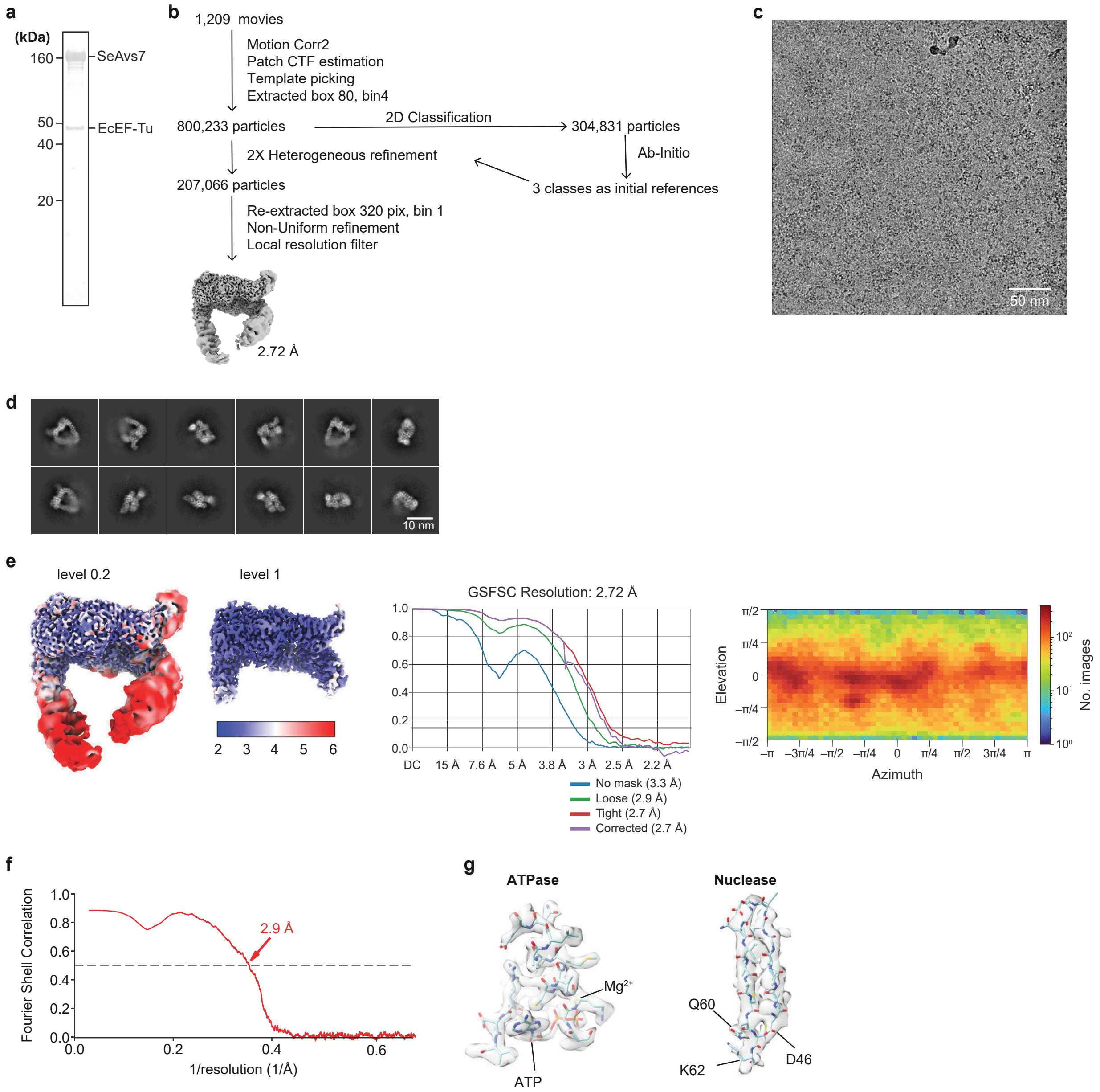


**Supplementary Fig. 10 | Cryo-EM data processing and validation for apo SeAvs7. a**, SDS-PAGE analysis of SeAvs7 co-purified with endogenous EcEF-Tu for cryo-EM grid preparation. **b**, Data processing workflow. **c**, Representative cryo-EM micrograph. **d**, Representative 2D class averages. **e**, Quality assessment metrics, including the local resolution map (left), gold-standard FSC curve (middle), and angular particle distribution (right). **f**, Map-to-model FSC curve. **g**, Representative cryo-EM density overlaid with the atomic model.

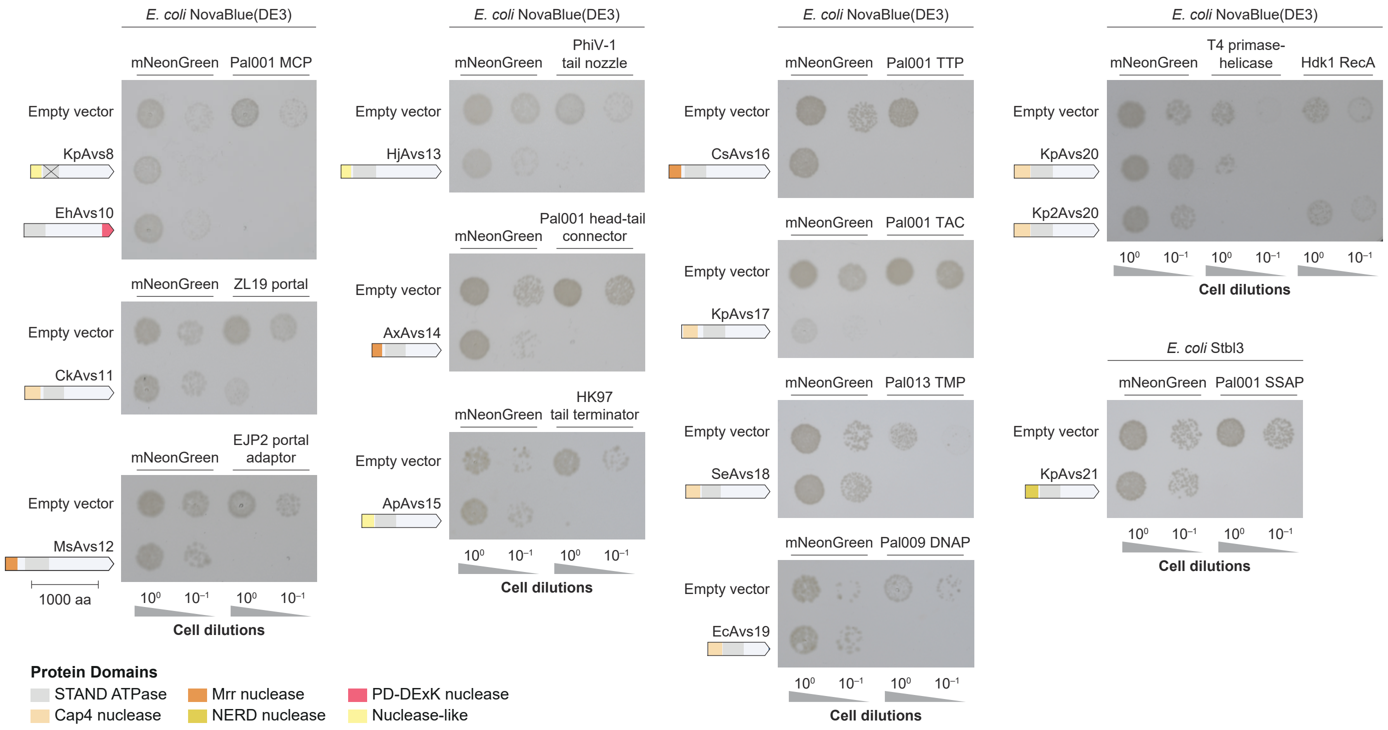


**Supplementary Fig. 11 | Co-expression of Avs–trigger pairs induces toxicity in *E. coli*.** Representative photographs of *E. coli* co-transformation assays with Avs proteins and phage triggers identified in Fig. 5.

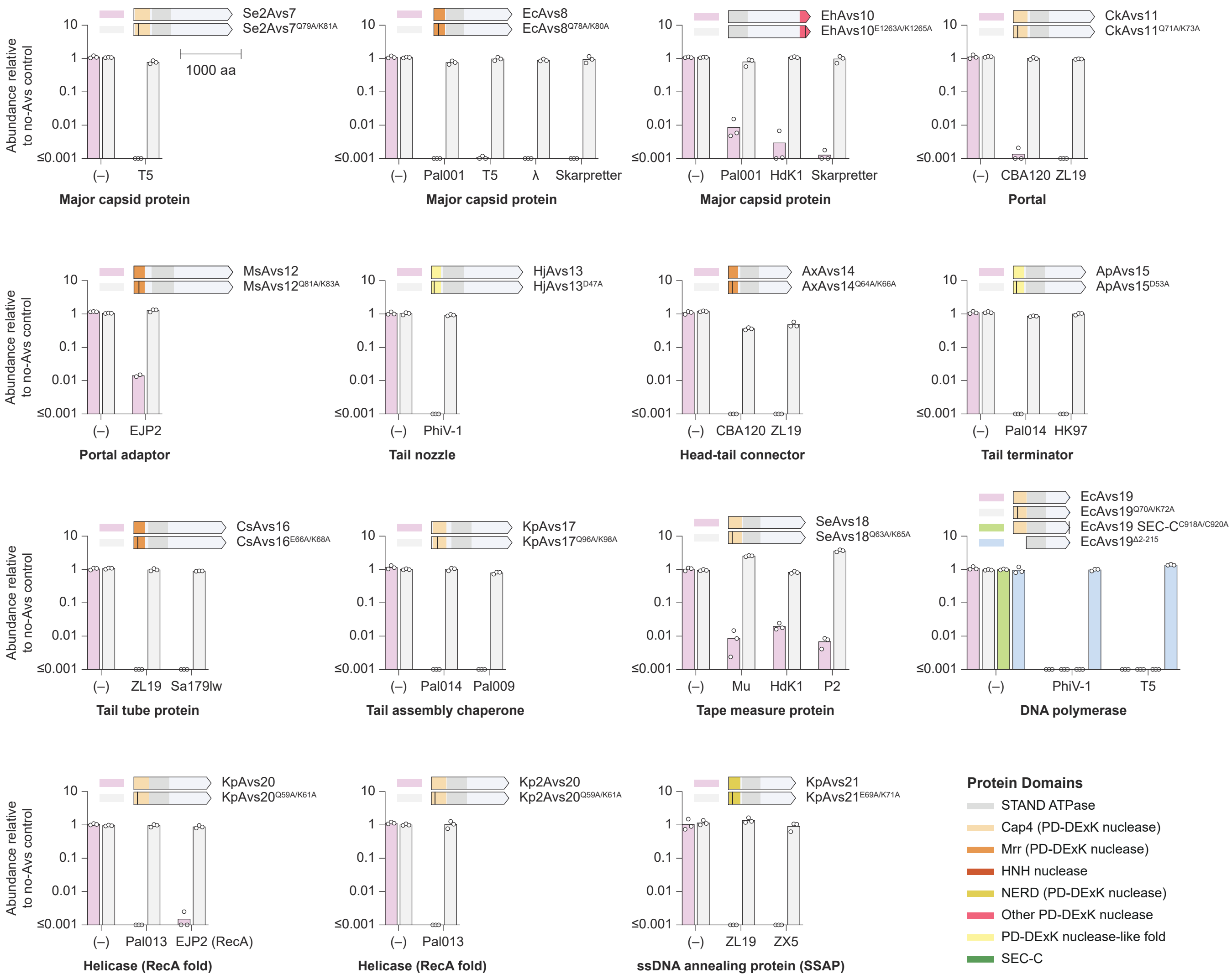


**Supplementary Fig. 12 | Avs effector domains are required for cellular toxicity.** Toxicity of wild-type Avs and effector domain mutants co-expressed with their cognate phage triggers in *E. coli*.

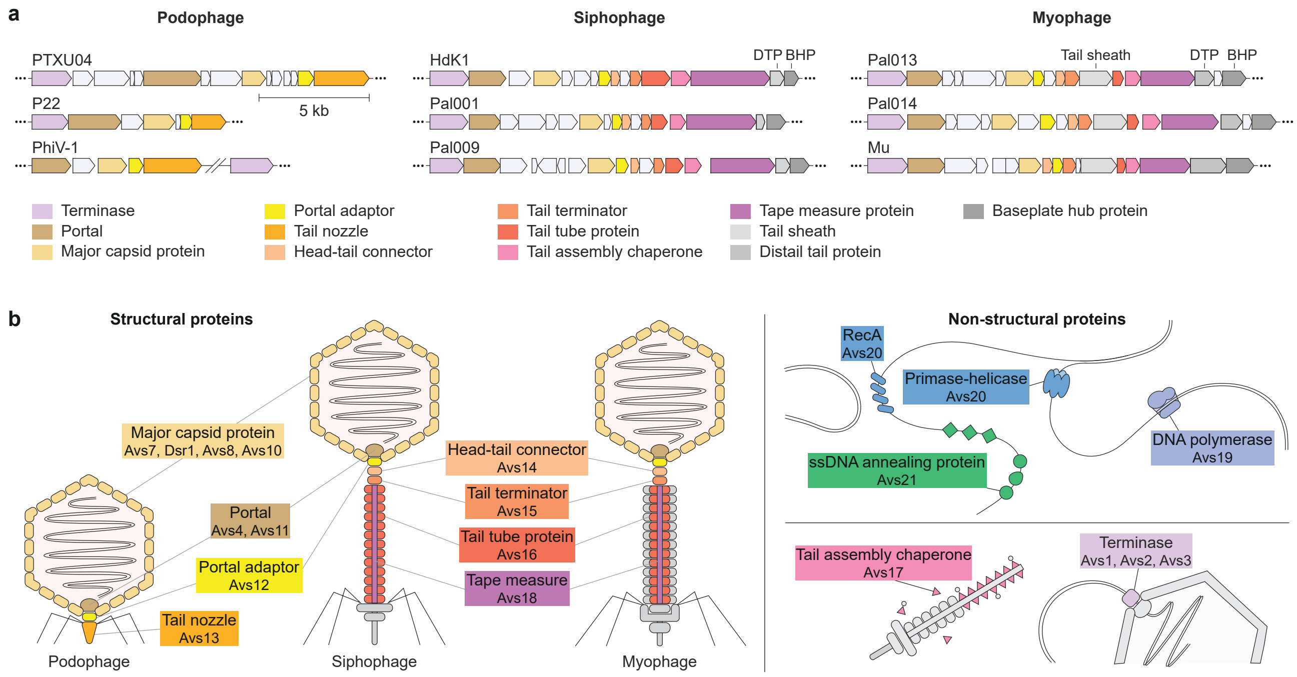


**Supplementary Fig. 13 | Avs targets are conserved across diverse tailed phages. a**, Genomic context of Avs targets across different representative phages. **b**, Schematic of the functional roles of the structural and replicative phage proteins recognized by distinct Avs families (Fig. 5c). DTP, distal tail protein; BHP, baseplate hub protein.

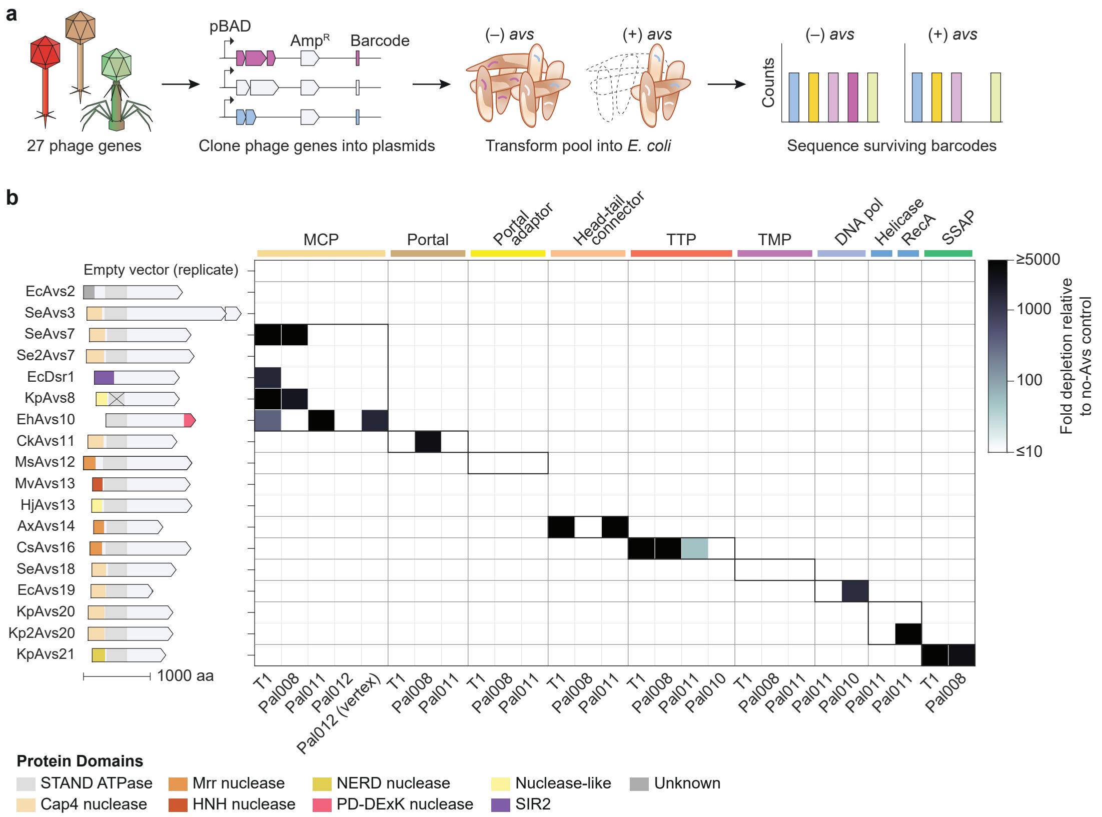


**Supplementary Fig. 14 | Expanded co-expression toxicity screen with an additional 27 phage genes.**

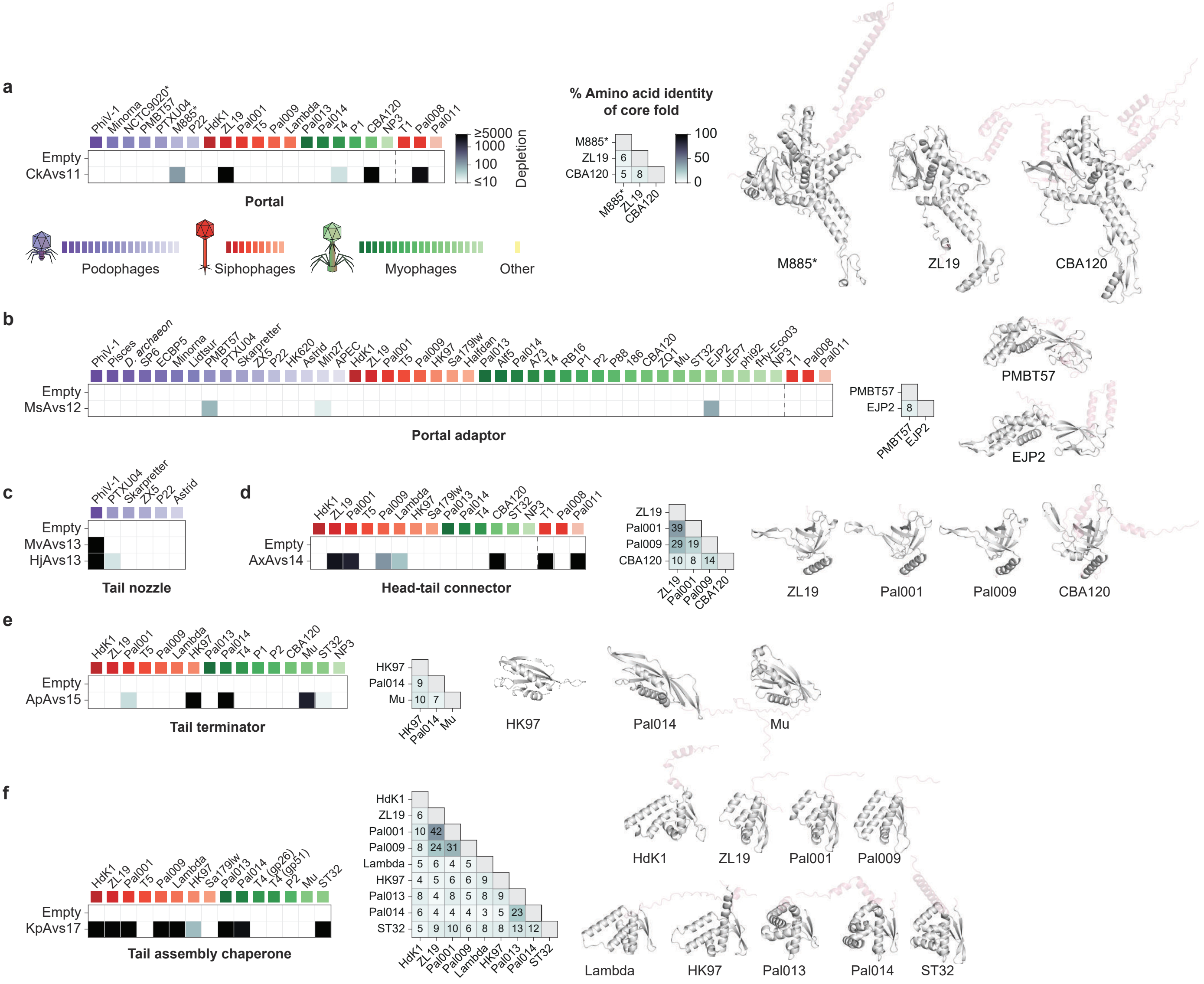


**Supplementary Fig. 15 | Avs pattern recognition of conserved core structural folds (related to Fig. 5b). a–f**, Toxicity quantification and structural comparisons of AlphaFold3 models for the portal (**a**), portal adaptor (**b**), tail nozzle (**c**), head-tail connector (**d**), tail terminator (**e**), and tail assembly chaperone (**f**). Toxicity data are replotted from Fig. 5b and Supplementary Fig. 14. Pairwise amino acid sequence identities between representative core folds are indicated; corresponding AlphaFold3 structures highlight these folds in dark gray. Sequence identities below ~5% are considered random. NP3, *Pseudomonas* phage NP3. Asterisks (*) denote prophages.

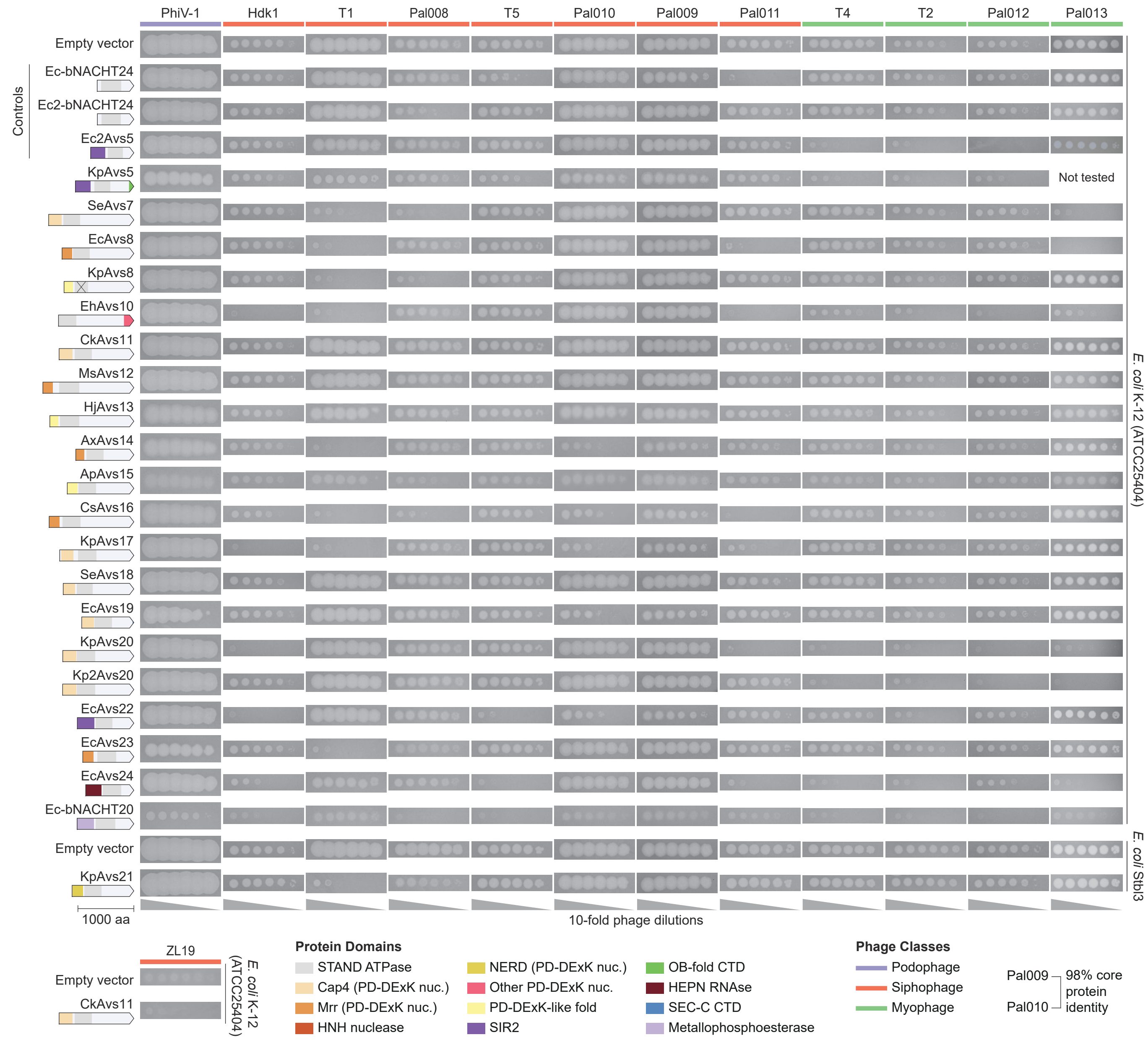


**Supplementary Fig. 16 | Anti-phage activity of Avs proteins in *E. coli*.** Plaque assays assessing Avs-mediated defense against a panel of 12 coliphages. Spots represent 10-fold serial phage dilutions (left to right). The cross mark on KpAvs8 indicates the natural absence of conserved nucleotide-binding residues in its STAND NTPase domain. Sequence identities for Pal009 and Pal010 are based on core structural and replicative proteins.

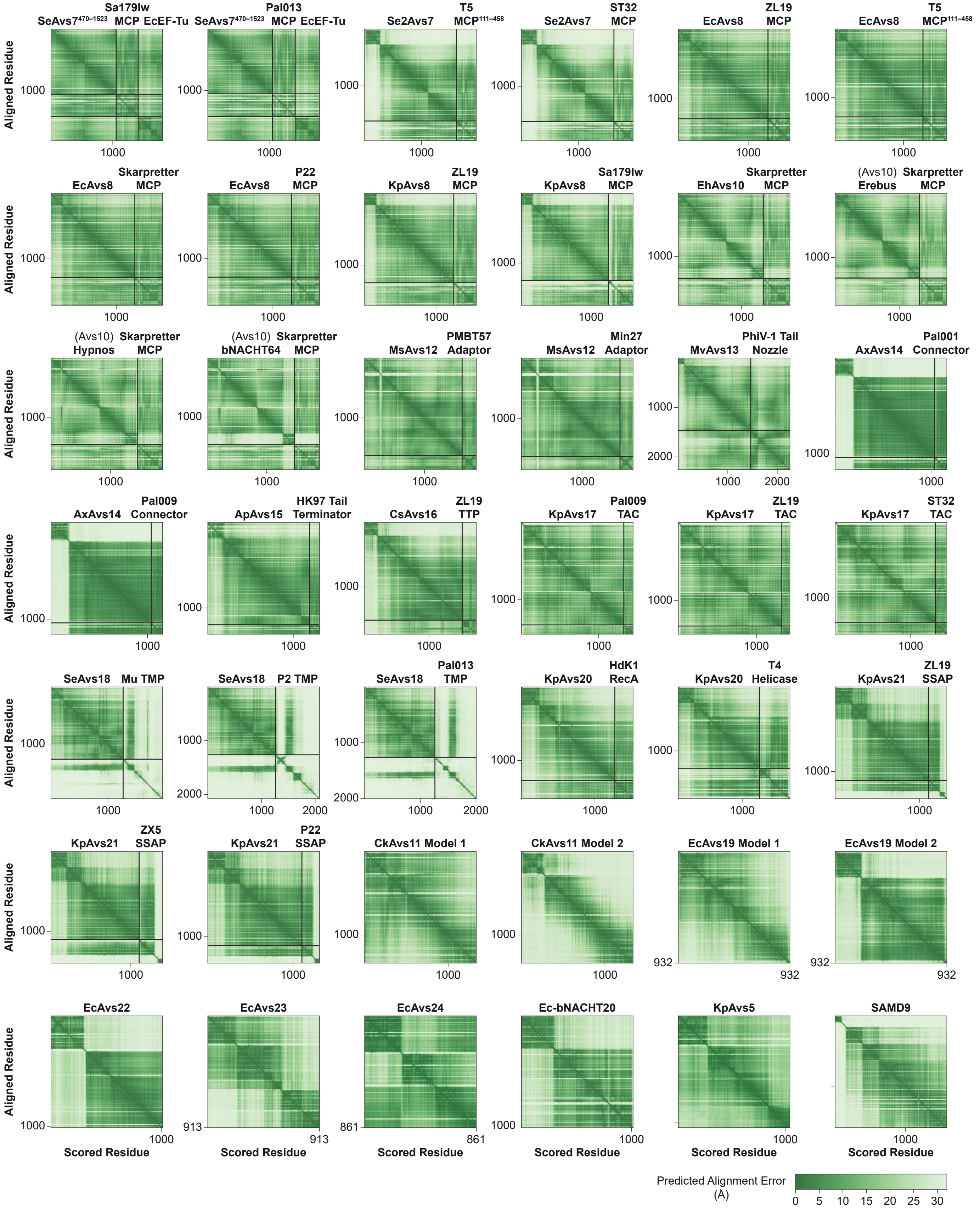


**Supplementary Fig. 17 | Quality of AlphaFold structural models.** Predicted alignment error (PAE) heatmaps for the AlphaFold2 and AlphaFold3 models presented in Fig. 6 and Extended Data Figs. 9–10.

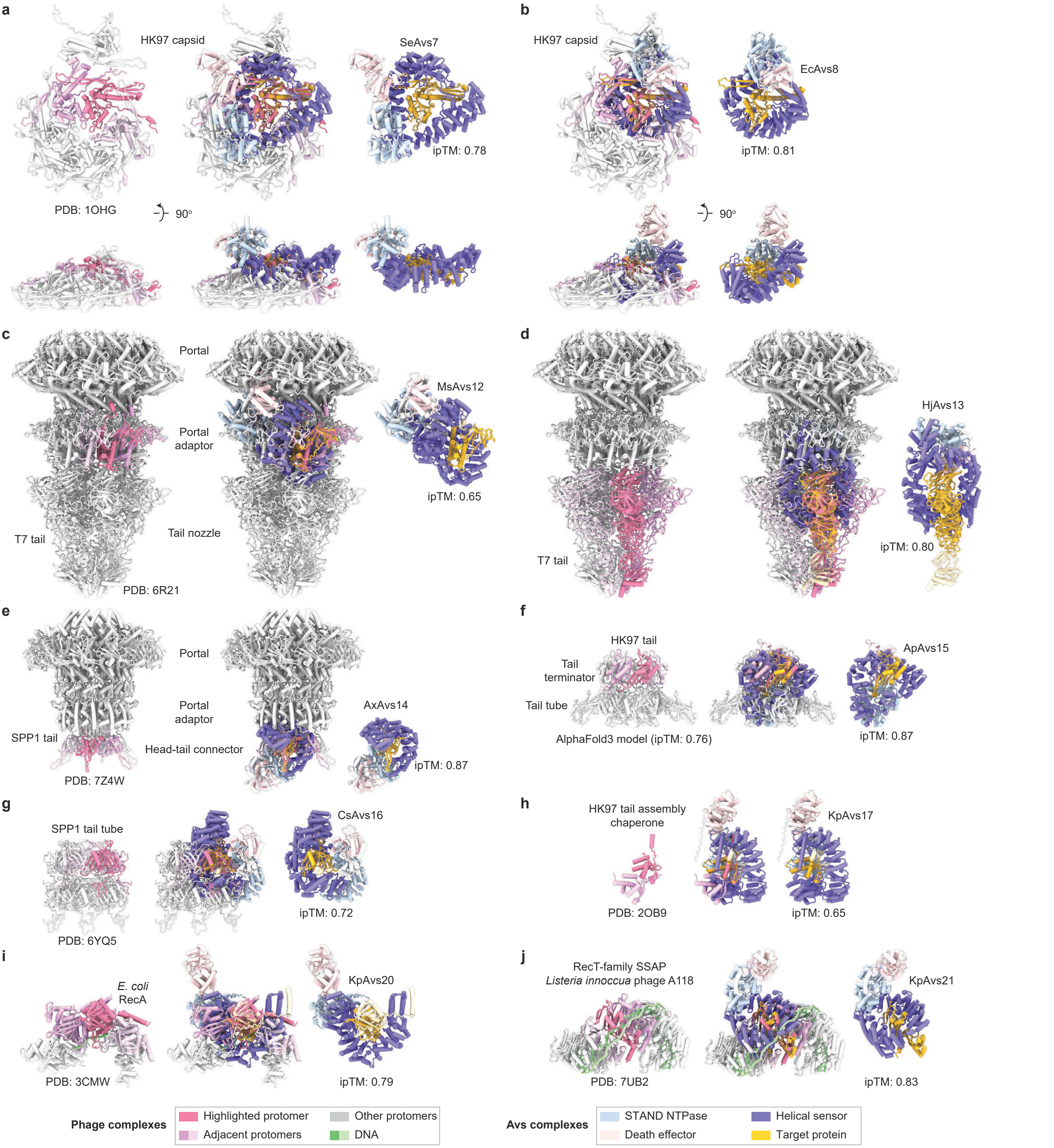


**Supplementary Fig. 18 | Avs binding is sterically incompatible with target protein oligomerization. a–j**, Structures of the HK97 capsid hexon (**a**, **b**), portal adaptor ring (**c**), complete tail nozzle (**d**), head-tail connector ring (**e**), tail terminator complex (**f**), tail tube (**g**), tail assembly chaperone (**h**), DNA-bound RecA (**i**), and DNA-bound ssDNA-annealing protein (**j**) overlaid with AlphaFold3 models of the respective Avs bound to a target monomer. The tail terminator complex is an AlphaFold3 model; all other target assemblies are experimental structures from the Protein Data Bank. ipTM, interface predicted template modeling score (0.0–1.0).
